# Evaluating the bioactivities of natural and synthetic fungal tropolone sesquiterpenoids in various cell lines

**DOI:** 10.1101/2023.07.19.549646

**Authors:** Timna C. Bergmann, Max Menssen, Carsten Schotte, Russell J. Cox, Cornelia Lee-Thedieck

**Author notes:** Prof. Dr. Cornelia Lee-Thedieck, Leibniz University Hannover, Institute of Cell Biology and Biophysics, Herrenhäuserstr. 2, 30419 Hannover.,2606.

## Abstract

Fungal specialized metabolites are known for their potent biological activities, among which tropolone sesquiterpenoids (TS) stand out for their diverse bioactivities. Here, we report bioactive effects of the recently discovered TS compounds 4–hydroxyxenovulene B and 4– dihydroxy norpycnidione, and the structurally related 4-hydroxy norxenovulene B and xenovulene B. Inhibition of metabolic activity after TS treatment was observed in Jurkat, PC–3 and FAIK3–5 cells, whereas MDA-MB-231 cells were unresponsive to treatment. Structurally similar epolones were shown to induce erythropoietin (EPO). Therefore, FAIK3-5 cells, which can naturally produce EPO, were applied to test the compounds in this regard. While no effect on EPO production in FAIK3-5 cells could be demonstrated, effects on their proliferation, viability, and morphology were observed depending on the presence of tropolone moieties in the molecules. Our study underlines the importance of relevant cell models for bioactivity testing of compounds with unknown mechanisms of action.

## INTRODUCTION

Natural products and their structural analogues have long been valuable sources of medical products due to their diverse chemical architectures and structural complexity. These active ingredients are categorized into primary and secondary (or specialized) metabolites ^1, 2^. Specialized metabolites are structurally diverse and are often optimized for specific biological functions. Advances in screening, isolation, and characterization methods for these metabolites have led to the discovery and development of novel drugs against various diseases ^3–6^ including cancer ^7, 8^, as well as treatments for drug-resistant infections ^9, 10^. However, the number of unexplored compounds, especially in fungi, remains high ^11^, despite recent progress in screening and characterization strategies. The potential of specialized metabolites to target persistent medical care challenges remains meaningful ^12^.

One promising family of bioactive fungal specialized metabolites are tropolone sesquiterpenoids (TS). TS compounds are a unique class of fungal meroterpenoids with a conserved core structure consisting of a humulene-derived macrocycle and pendent tropolones connected via one or two dihydropyran rings ^13^ (Figure 1). In 1993 the bistropolone pycnidione **1** ^14^ and its diastereomer eupenifeldin **2** ^13^ were the first reported TS compounds. They display a wide variety of diverse biological activities. For example, reported bioactivities of pycnidione (**1**), first isolated from an unidentified *Phoma* sp. ^14^, include anticancer activities ^15–17^, as well as weak antibacterial ^18^ and antimalarial properties ^15, 18, 19^. Pycnidione (**1**) is part of the “epolone family” of TS, known for their oxygen-independent induction of erythropoietin (EPO) gene expression in EPO-specific reporter cell lines ^20, 21^. Similarly, the structurally related TS compound eupenifeldin (**2**), isolated from *Eupenicillium brefeldianum,* acts as an antitumor agent against human HTC colon carcinoma cell lines ^13^, and further cytotoxic effects were recorded against several human cell lines ^22, 23^. Eupenifeldin (**2**) also displays antifungal properties (vs. *Candida albicans*) and antimalarial activity (vs. *Plasmodium falciparum* K1) ^22^.

**Figure 1:**
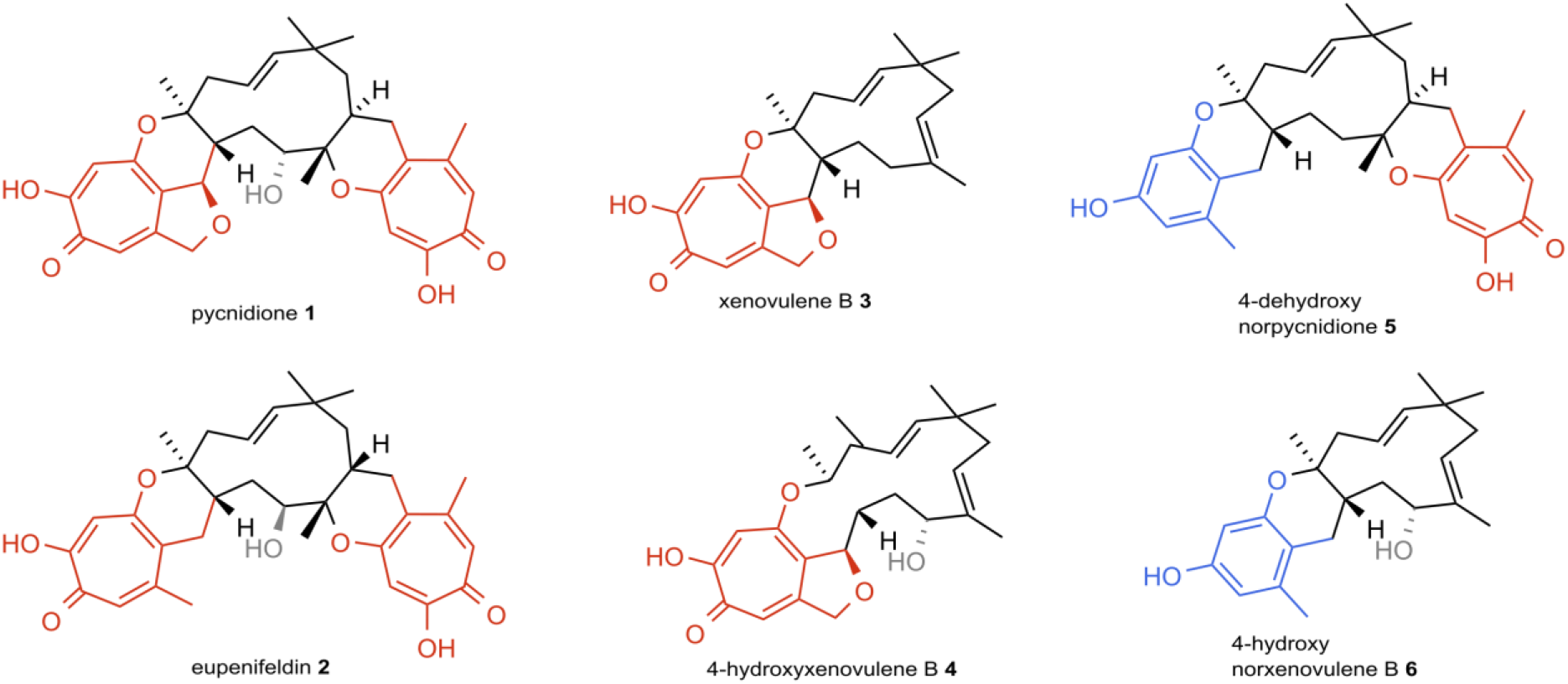
Stereochemistry of selected tropolone sesquiterpene (TS) compounds (**1**-**5**) and a control compound (**6**) lacking a tropolone moiety ^24^. TS compounds share a typical chemical structure, with a conserved 11-membered humulene-derived macropcyclic core structure connected to the pendent tropolone moieties via one or two dihydropyran rings. Key features are highlighted in red (= polyketide derived tropolones), blue (= benzopyranyl moiety) and grey (C-10 hydroxylation).

Chemical diversity in TS metabolizing fungi is due to diversity in fungal biosynthetic gene clusters (BGCs), which encode metabolite synthesis. The exact determination of the stereochemistry of TS has been difficult and has only recently been accurately reassigned ^24, 25^. Redesigning and refactoring known BGCs and effective heterologous expression in engineered hosts like *Aspergillus oryzae* NSAR1 can accelerate the synthesis process of TS compounds and create platforms for new compound discovery. Combining genes of known BGCs can create new specialized metabolites with specific characteristics and bioactivities ^26^. Recently, Schotte et al. described the successful isolation and identification of several TS compounds ^27^, including xenovulene B (**3**), first reported in 1997 ^28^ and fully characterized by Schor et al. ^29^, 4-hydroxyxenovulene B (**4**), as well as the two hitherto unknown compounds 4-dehydroxy norpycnidione (**5**) and 4-hydroxy norxenovulene B (**6**) by recombining genes of three key TS BGC in *A. oryzae* NSAR1 ^27, 29^. To date, no biological activity of these compounds has been reported in literature. Over the last decades a variety of bioactivities of other TS compounds, most of which belong to the eupenifeldin series, has been discovered ^30–34^, and given the high potency of structurally related TS compounds, similar effects can be expected by compounds **3**, **4**, **5** and **6**. Particularly pycnidione (**1**) and eupenifeldin (**2**) are well characterized and were shown to exhibit versatile bioactivities in different cell lines ^13, 15–17, 21–23^. However, early drug discovery studies often used genetically engineered (cancer)-cell lines that cannot accurately reflect physiological conditions, or rely on indirect measurements, putatively leading to false-positive results due to assay interferences ^35^. In particular, the counterintuitive phenomenon of signal activation by luciferase inhibitor molecules can increase signal strength indistinguishable from transcriptional activation of the reporter gene [57]. In addition, the malignant transformation that cells must undergo to become cancerous affects the cell’s protein function and DNA methylation patterns, which in turn influence the cells’ response to chemical compounds ^36^. Thus, different cell lines are expected to have different sensitivities towards the bioactivity of novel compounds. However, identifying appropriate cell models for bioactivity testing studies is particularly difficult when the putative bioactivities and underlying mechanisms of action have not yet been discovered. In this case, it is advantageous to test compounds in a diverse group of cancerous and non-cancerous cell lines to explore the potential bioactivities in dependence of cell type specific and transcriptional regulation. This also means that sustainable and simplified cell models for *in vitro* studies are desirable, especially when studying the compound coordinated induction of proteins such as EPO, which are tightly regulated at the transcriptional ^37^ and translational ^38^ level. Recently, Imeri et al. introduced the immortalized clonal cell line FAIK3-5, which was derived from mouse renal EPO-producing (REP) cells and can produce EPO for at least 30 passages ^39^. By using this cell line to test the bioactivity of the newly discovered TS and related compounds ^27^, we are closer to a physiological model than when using genetically engineered reporter cell lines, while the cells also meet the necessary requirements for toxicological and cell proliferation studies.

In this study we aimed to evaluate the bioactivities of the recently discovered tropolone sesquiterpenoids **4** and **5**, as well as the structurally related compounds **3** and **6**, for which no bioactive effects have been reported yet. For this purpose, their effects on metabolic activity were tested in cancer cell lines originating from pancreatic (PC–3), lymphoid (Jurkat) or breast cancer tissue (MDA–MB–231) as well as the non–cancerous renal FAIK3–5 cell line and compared to the well characterized compounds **1** and **2**, which served as positive controls. FAIK3-5 cells are suitable to assess the full range of activities described for similar compounds. Therefore, they were chosen for a detailed analysis of the compounds’ influence on oxygen-independent EPO induction, cell proliferation, viability, and morphology. With this we are able to gain a better understanding of the biological functions of TS moieties, which could support synthetic biology in redesigning TS compounds to meet medical needs by increasing their efficacy while mitigating unwanted side effects.

## RESULTS

### Tropolone-bearing TS compounds are bioactive in cancerous and non–cancerous cell lines

For preliminary identification of the bioactive potential of the individual TS compounds **1**–**6**, their influence on the metabolic activity of MDA–MB–231, PC–3 and Jurkat cancer cell lines derived from different cancer entities was evaluated using the CellTiter - Blue Cell® Viability Assay. Additionally, murine FAIK3 – 5 cells were tested, which provide an interesting model for studies of epolones and other TS compounds, as the cells originate from renal interstitial fibroblasts, the primary physiological production site of EPO.

The half maximal effective concentration (EC_50_) values for individual compounds vary between the different cell lines, showing a wide range of effective concentrations (**Figure 2** and Supplementary Figure **S1**). For example, the bistropolone controls exhibit EC_50_ values ranging from 0.20±0.07 (mean±SD) µM (pycnidione **1**) and 0.07±0.04 µM (eupenifeldin **2**) in Jurkat cells to 16.65±3.14 µM and 16.45±3.14 µM in MDA–MB-231 cells, respectively. The breast cancer cell line MDA–MB–231 appears to have an intrinsic resistance to the compounds used, as EC_50_ values varied between 16.28±2.80 µM and 18.58±3.31 µM for all compounds, which is in the same range as observed for the DMSO solvent control. Overall, the bistropolones **1** and **2** tend to be more potent as they have a higher bioactivity, i.e., lower EC_50_ values compared to the monotropolones **3** and **4**.

**Figure 2.**
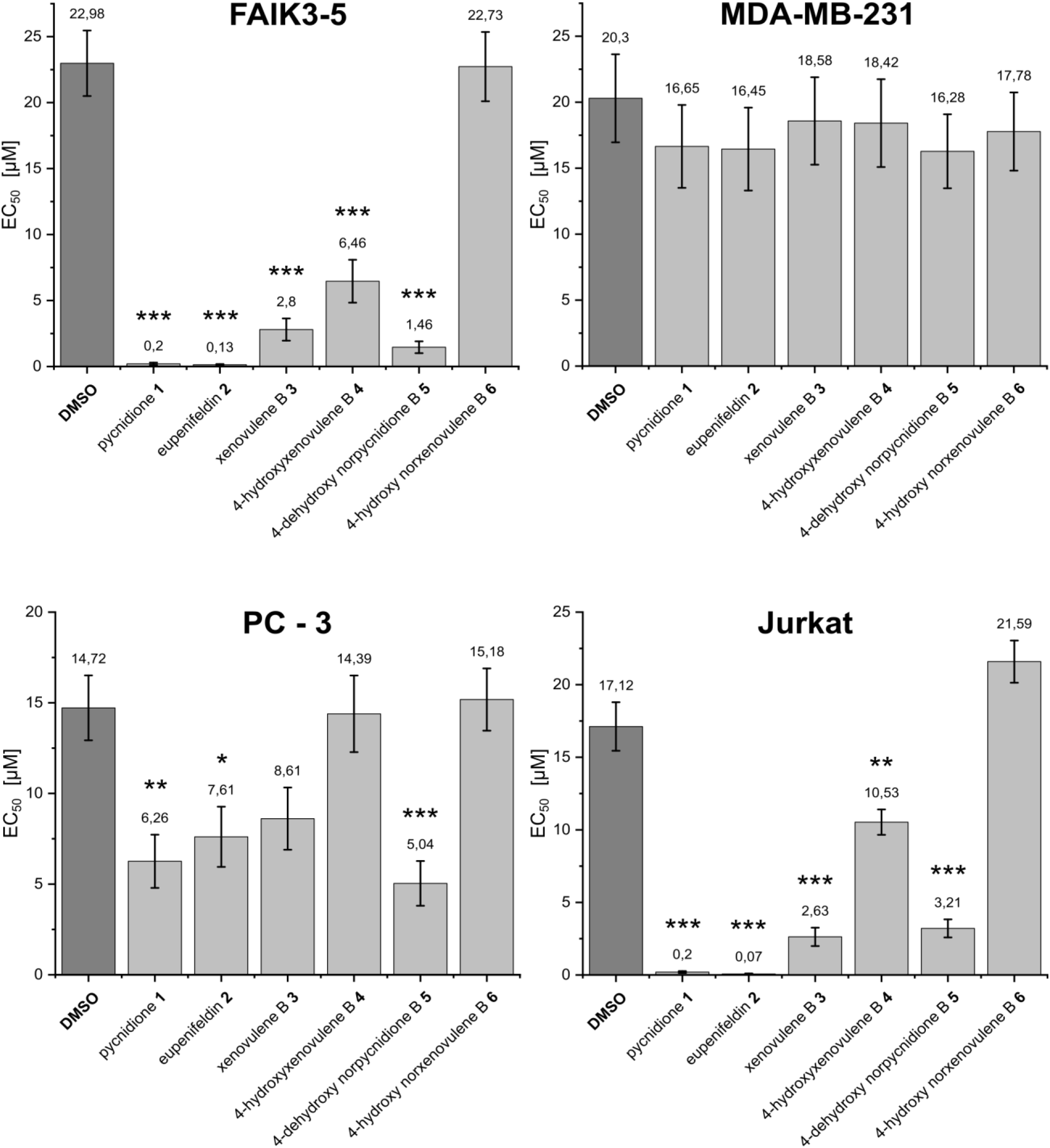
EC_50_ of the tropolone sesquiterpenoids (TS) pycnidione (**1**), eupenifeldin (**2**), xenovulene B (**3**), 4-hydroxyxenovulene B (**4**), 4-dehydroxy norpycnidione (**5**) and tropolone-lacking compound 4-hydroxy norxenovulene B (**6**) and their respective DMSO vehicle controls in murine FAIK3–5 cells and the cancer cell lines MDA–MB–231, PC–3 and Jurkat. Numbers above each graph bar indicate the calculated mean EC_50_ of three independent experiments (N = 3), with error bars corresponding to the calculated standard errors. Pairwise mean comparisons were calculated to show statistically significant differences between calculated EC_50_ of DMSO vehicle controls and TS compounds treated samples (see also Supplementary Figure **S1 b**). Significances range from 0.05-0.01 (*), <0.01-0.001 (**) to <0.001 (***). Further details on modelling and EC_50_ derivation are given in the statistical supplementary statS1.

To assess the magnitude of the actual effects of the compounds on the cell lines, mean comparisons of the EC_50_ values of the compounds and DMSO controls were calculated for each cell line, to distinguish between statistically significant cell responses to the compounds used and cytotoxic side effects of the solvent (Supplementary Figure S1b). For MDA–MB-231 cells no differences between the solvent and compound treatment were observed. Thus, MDA–MB-231 cells did not respond to the administered substances and the observed decrease in metabolic activity could be attributed solely to the toxicity of DMSO. For the other cell lines, the tropolone moieties present in the molecule appeared to be the determining factor for their bioactivity, as the tropolone-lacking compound **6** showed no significant effect in any cell line compared to the solvent controls. This assumption was further supported by the finding that the bistropolone controls **1** and **2** could be significantly distinguished from DMSO effects in all responding cell lines, whereas monotropolones appeared to have an attenuated effect on the tested metabolic activity. The effect was particularly evident in PC-3 cells where compound **5** was the only monotropolone showing a statistically significant effect on pancreatic cancer cells that was distinct from the effect of the solvent.

Based on these results, we describe the first bioactivities of the monotropolones xenovulene B (**3**), 4-hydroxyxenovulene B (**4**) and 4-dehydroxy norpycnidione (**5**) in Jurkat and FAIK3–5 cells. PC - 3 cells were less sensitive to these monotropolones, with only compound **5** having a significant effect on cell metabolism, whereas renal FAIK3 - 5 and lymphoid Jurkat cells were the most sensitive to the compounds applied. MDA–MB-231 cells, known to be chemoresistant to diverse anticancer agents, show a primary resistance to TS compounds in general.

### Tropolone sesquiterpenoid compounds have minimal effects on erythropoietin protein production in REP – derived cells

One feature of specific TS compounds comprising to the so-called epolone-family, is the hypoxia-independent induction of EPO reporter gene activity and protein expression that has been described for several EPO–reporter cell lines ^20, 21^. As we were able to verify the bioactivities of the newly discovered TS compounds **3** - **5** in several cancer cell lines and non–cancerous cells, we aimed to evaluate their potential to induce EPO production in a cell model that is able to naturally produce EPO and is thus more physiologically accurate than reporter cell lines. To assess, if the utilized compounds can also stimulate natural EPO production in REP-derived FAIK3-5 cells *in vitro*, EPO protein content after TS–treatment was measured via ELISA. In accordance with former studies on pycnidione **1** activity ^20, 21^ and preliminary experiments (see Supplementary Figures **S 2**–**S 4**), FAIK3 – 5 cells were treated with 6 µM or 12 µM of TS compounds and the respective DMSO vehicle controls. For more information on the evaluation of optimal compound concentrations for treatment of FAIK3–5 cells, see Materials and Methods section.

We found variability between biological replicates and no statistically significant effect of the compounds on EPO protein production (**Figure 3 a**). One possible reason could be epigenetic silencing of EPO synthesis by hypermethylation of EPO promoter and regulatory gene sequences^40^. To counteract this, samples were treated with 1 µM of the DNA methyltransferase inhibitor 5-Azacytidine (5-Aza) for six days. As the vehicle control DMSO itself might support hypermethylation events within the cells ^41^, the effectiveness of 5-Aza treatment in enhancing EPO production of FAIK3-5 cells was also assessed in DMSO vehicle controls (0.6 % (*v/v*) and 1.2 % (*v/v*)) that showed an increase in measured basic EPO concentrations (**Figure 3 b**).

**Figure 3:**
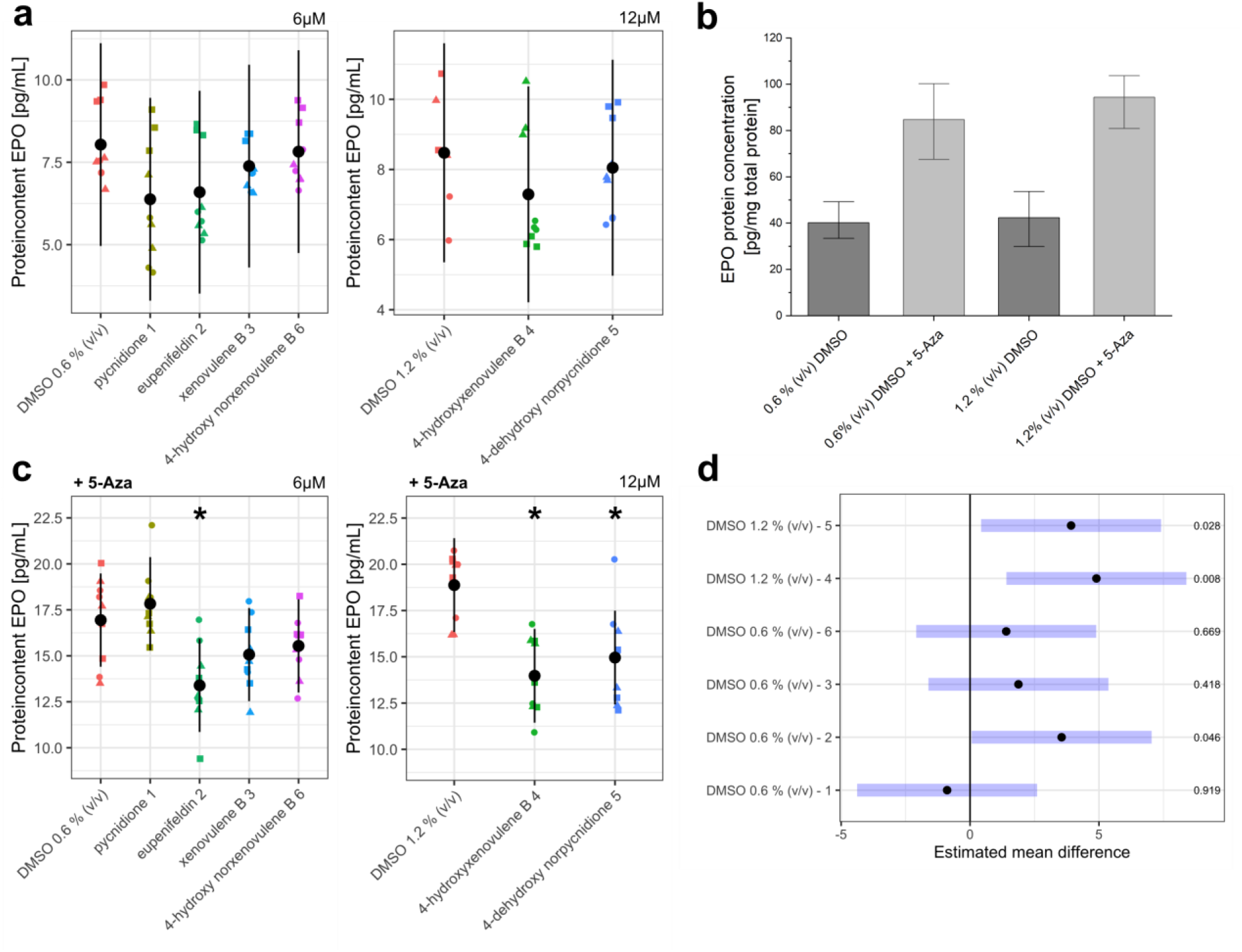
Erythropoietin (EPO) content of FAIK3-5 samples treated with the tropolone sesquiterpenoids (TS) pycnidione (**1**), eupenifeldin (**2**), xenovulene B (**3**), 4-hydroxyxenovulene B (**4**), 4-dehydroxy norpycnidione (**5**) and tropolone-lacking compound 4-hydroxy norxenovulene B (**6**) and their respective vehicle controls (DMSO0.6 % (*v/v*), DMSO 1.2 % (*v/v*)). **(a)** EPO protein concentrations measured in three biological replicates (●=1,▴=2, ▪=3) of FAIK3-5 cells treated with 6 µM of TS compounds **1**, **2**, **3**, tropolone-moiety lacking compound **6** and their respective DMSO vehicle control (0.6 % (*v/v*)) or 12 µM of TS compounds **4**, **5**, and their respective DMSO vehicle control (1.2 % (v/v)) without pre-treatment of 5-Azacytidine (5-Aza). Black dots are indicating the model based least square means and their pointwise 95 % confidence intervals (black bars). **(b)** Measurement of basic EPO protein content in FAIK3-5 DMSO vehicle controls (0.6 % (*v/v*) and 1.2 % (*v/v*) with and without 5-Aza treatment for 6 days. Error bars indicate the standard deviation of mean EPO concentrations. **(c)** EPO protein concentrations measured in three biological replicates (●=1,▴=2, ▪=3) of FAIK3-5 cells treated with 6 µM of TS compounds **1**, **2**, **3**, tropolone-moiety lacking compound **6** and their respective DMSO vehicle control (0.6 % (*v/v*)), or 12 µM of TS compounds **4**, **5**, and their respective DMSO vehicle control (1.2 % (v/v)) with pre-treatment of 5-Azacytidine (+5-Aza). Black dots are indicating the model based least square means and their pointwise 95% confidence intervals (black bars). Treatments marked with **\*** are statistically significantly different from their respective vehicle controls. **(d)** Pairwise mean comparisons showed statistically significant differences in EPO protein concentrations 4-hydroxyxenovulene B (**4**), 4-dehydroxy norpycnidione (**5**) and eupenifeldin (**2**) compared to their respective vehicle controls. Black dots indicate back transformed means of measured EPO concentrations and their 95% confidence intervals (blue bars). Respective p-values for each calculation are displayed on the right side. The estimated contrasts and corresponding confidence intervals are given in the statistical supplementary statS2 and statS3.

However, additional treatment with TS compounds showed no enhancing effect on EPO protein synthesis after 5-Aza treatment compared to the respective vehicle controls (**Figure 3 c**). Treatment with 4-hydroxyxenovulene B (**4**) (p = 0.008), 4-dehydroxy norpycnidione (**5**) (p = 0.028) and eupenifeldin (**2**) (p = 0.046) decreased the measured EPO protein content compared to the vehicle control, yet, the effect of eupenifeldin (**2**) on 5-Aza treated FAIK3-5 cells was considered as marginal (**Figure 3 d**).

Summing up, our results showed that the utilized TS compounds did not enhance the basal EPO production of 7.5±1.5 pg/mL in the REP cell line model. Hypomethylation of the FAIK3-5 cells with 5-Aza elevated EPO levels about 2-fold to 15.8±2.7 pg/mL in the cell lysates of treated and untreated FAIK3-5 cells and their controls, but treatment of hypomethylated cells with TS compounds did not induce EPO production additionally.

### Structurally related tropolone sesquiterpenoids promote similar morphological alterations in FAIK3-5 cells

Physiologically, REP cells display specific morphological features that resemble neuronal cells with protrusions extending in multiple directions, and a stellate or arboroid shape ^37, 42^. Similar cell shapes have been described for EPO–producing FAIK3–5 and Replic cells ^39, 43^. However, the conversion of functional REPs cells into morphologically distinct myofibroblasts has also been reported for primary REP cells as well as for the used FAIK3–5 cell lines ^39, 44^, which, in addition to epigenetic regulation, could be a cause for their lack of response to TS compounds. For this reason and because general bioactivity of the compounds has been observed, we aimed to assess the influence of the monotropolone controls **1** and **2**, the novel bistropolones **3**–**5** and the tropolone–lacking compound **6** on cell morphology of FAIK3–5 cells. In particular, the analysis was performed to provide evidence that morphological features can have predictive value in early stages of compound characterization, because substantial alterations in cell morphologies are frequently observed during screening studies of natural or synthetic compounds ^45, 46^. FAIK3-5 cells were treated for 24 hours with 6 µM or 12 µM of the tested TS compounds to investigate their influence on morphology and cell growth (**Figure 4 a**)

**Figure 4:**
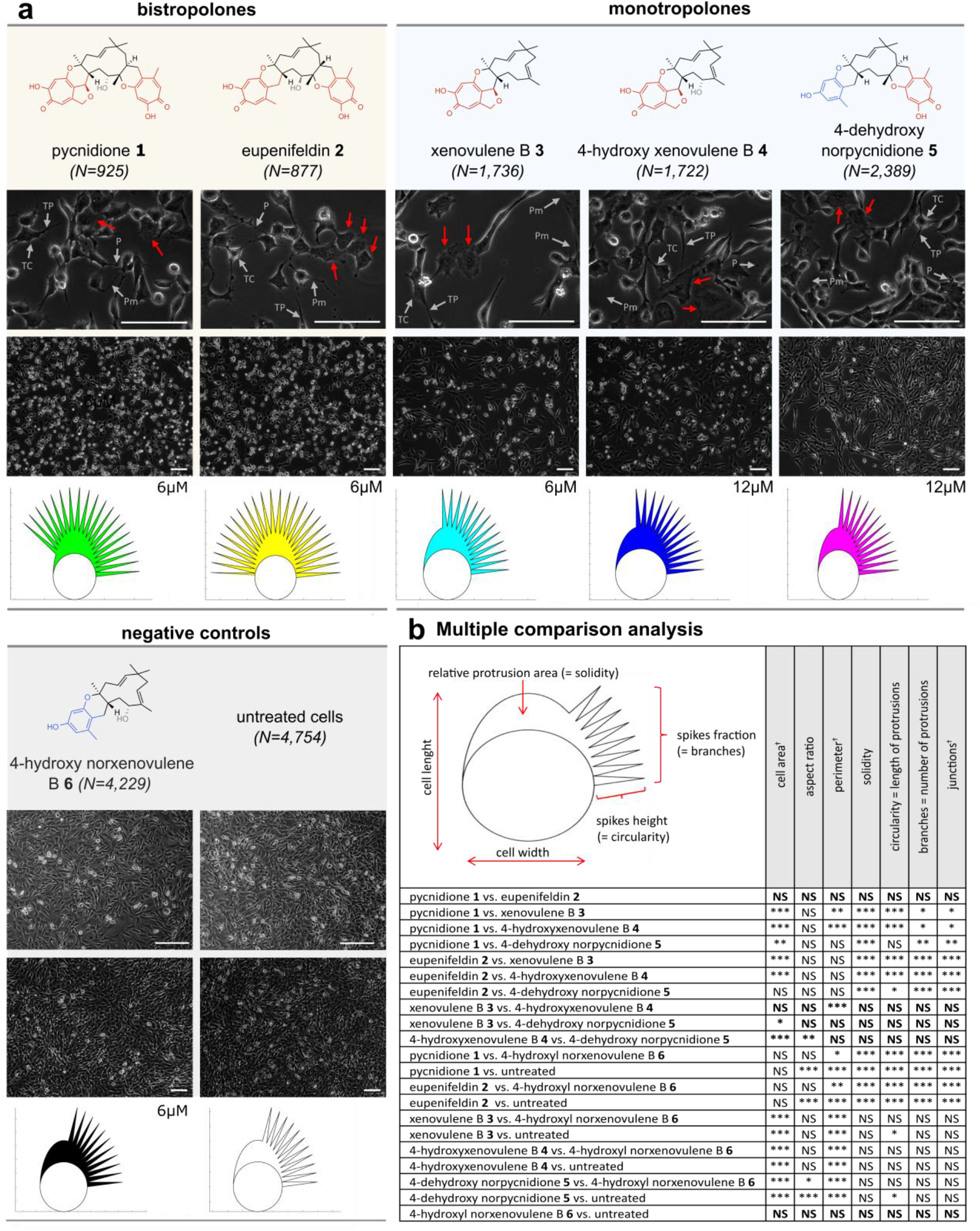
Alterations of cell morphology of FAIK3-5 cells after 24 h of treatment with the TS compounds pycnidione (**1**), eupenifeldin (**2**), xenovulene B (**3**), 4-hydroxyxenovulene B (**4**), 4-dehydroxy norpycnidione (**5**), the tropolone-lacking compound 4-hydroxy norxenovulene B **6** and untreated controls. **(a)** Brightfield images of samples treated for 24 h with compound **1**-**5** and negative controls (compound **6** and untreated cells). Error bars = 100 µm. Red arrows indicate enlarged and flat cells that were visible to varying degrees in each tropolone-treated sample. All samples treated with TS - compounds (**1** - **5**) show the occurrence of telocytes (TC) with characteristic telopodes (TP) containing of podomers (Pm) and podons (P). TS treated cells and negative controls were additionally visualized in *PhenoPlot*, in which quantifiable cell features such as cell width and length, relative area of protrusions as well as their proportion and size (spike fraction and spike height) are displayed for each sample condition. **(b)** Automated cell segmentation was conducted by using the deep-learning algorithm *Cellpose* ^48^. Multiple comparison analysis for analyzed shape descriptors and morphometric parameters (cell area, aspect ratio, perimeter, relative protrusion area as well as length and number of protrusions/branches and branch junctions) were measured on n=194 to n=1,773 cells per image and a total of n = 16,632 cells from the different test conditions from three independent experiments in Fiji (version 1.53c). Statistical analysis of data was conducted with R (version 4.2.1). All parameters marked with (†) are not depicted in *PhenoPlot* graphs. Significances range from p>0.05 (NS), 0.05-0.01 (*), <0.01-0.001 (**) to <0.001 (***). The estimated contrasts and corresponding confidence intervals are given in the statistical supplementary statS4 and statS5.

Treatment with the bistropolone positive controls (**1**, **2**) induced a neuron-like morphology in FAIK3-5 cells, with a high incidence of cells with typical morphological characteristics of telocytes, with long extensions or "telopodes" of tens to hundreds of nm, composed of ultrathin parts called "podomers" with dilated parts known as "podons" that can connect FAIK3-5 cells. While treatment with monotropolones (**3**, **4** and **5**) generally resulted in an epithelial-like cell morphology, telopodes were also present after monotropolone treatment, but to a much lesser extent. Furthermore, we observed the emergence of flat and enlarged cells in all treatment conditions with the tropolone-containing compounds **1**-**5**. Notably, treatment with the tropolone-lacking compound 4-hydroxy norxenovulene B (**6**) and the negative controls yielded no alteration in cell morphology or cell confluence in comparison to untreated FAIK3-5 cells, suggesting that tropolone moieties are responsible for changes in cell phenotype and cell growth. To quantify the observed morphological differences, the morphometric analysis was deepened by applying the deep learning-based cell segmentation algorithm *Cellpose* to the obtained microscopy images and subsequent multiple comparison tests of the treatment means on shape descriptors and morphometric parameters (cell area, aspect ratio (AR), perimeter, relative protrusion area (=*solidity*), length of protrusions (=*circularity*) as well as number of protrusions and branches) (**Figure 4 b**). The results of the analysis were visualized with the Matlab (Version R2021b) toolbox PhenoPlot ^47^. Visualized plots concur with the conducted multiple comparison analysis. Treatment with structurally similar compounds led to similar morphologies. Accordingly, the multiple comparison analysis revealed no statistically significant differences (p>0.05) between tested shape descriptors and morphometric parameters within the group of samples treated either with bistropolones (**1**, **2**) or negative controls (compound **6** and untreated samples). Cells treated with the monotropolone 4-dehydroxy norpycnidione (**5**), which belongs to the xenovulene-/pycnidione-type TS, characterized by the trans-fusion at C-8/C-9 position of the central α-humulene derived macrocycle, show a higher morphological equivalence to cells treated with the structurally distinct TS compounds xenovulene B (**3**) and 4-hydroxyxenovulene B (**4**) than to any of the other investigated compounds.

In summary, we observed a high morphological similarity of cells treated either with mono- or bistropolones and a lack of biological responses after 4-hydroxy norxenovulene B (**6**) treatment. This finding indicates that the observed alterations in morphology relate to the tropolone moieties present on the molecules, which seem to be responsible for the observed bioactivities.

### Tropolone moieties induce a downshift of cell proliferation and increased cytotoxicity upon dual-substitution

Besides a strong morphological correlation between phenotypes of cells treated with mono– or bistropolones, we observed in the course of the experiments to study cell morphologies that the visible cell density and thus cell growth appeared to be decreased in samples treated with bioactive TS compounds (**Figure S 3**). Therefore, we analyzed cytotoxicity and effects of TS compounds on proliferation in CellTrace™ Violet (CTV) stained FAIK3-5 cells, aiming to determine if the observed changes in cell morphology are accompanied by proliferative inhibition, as previously described for lung cancer cells treated with the positive control pycnidione (**1**) ^17^.

The proliferative inhibition of selected compounds was evaluated by TS treatment of the CTV-stained cells with 4 µM, 6 µM, 8 µM and 12 µM for 24 h and subsequent analysis of the mean fluorescence intensity (MFI) decrease resulting from cell proliferation, with higher MFI signals corresponding to lower proliferation. (**Supplementary Figure S 4**). We observed increased MFI values compared to untreated samples indicating proliferation inhibition of FAIK3-5 cells after treatment with the positive controls **1** and **2**, and compounds **3** and **5** after treatment with all applied concentrations compared to the untreated controls. For 4-hydroxyxenovulene B (**4**), a decrease in proliferation was observed with increasing compound concentrations, with the highest decline at a compound concentration of 12 µM.

Treatment with all utilized TS compounds at concentrations which showed the highest effects, led to a significantly decreased proliferation in treated cells compared to the untreated control (p<0.001), as shown by the low ratio of MFI values of day 1/day 3 (**Figure 5 a**) and respective shifts to higher fluorescence intensities visible in the histograms (**Figure 5 b**). Corresponding to previous experiments, the tropolone moiety lacking compound 4-hydroxy norxenovulene B (**6**) had no measurable effect on FAIK3-5 proliferation (p=0.951) and showed a statistically significant difference in MFI ratios of day 1/day 3 compared to all samples treated with TS compounds **1-5** (p<0.001, **Figure 5 c**). Moreover, as there were no detectable differences between the different tropolone-bearing compounds, these results strongly indicate a key role of the tropolone-moiety in proliferation inhibition, independent of its single or dual substitution to the central 11-membered α-humulene-derived core structure of the compounds.

**Figure 5:**
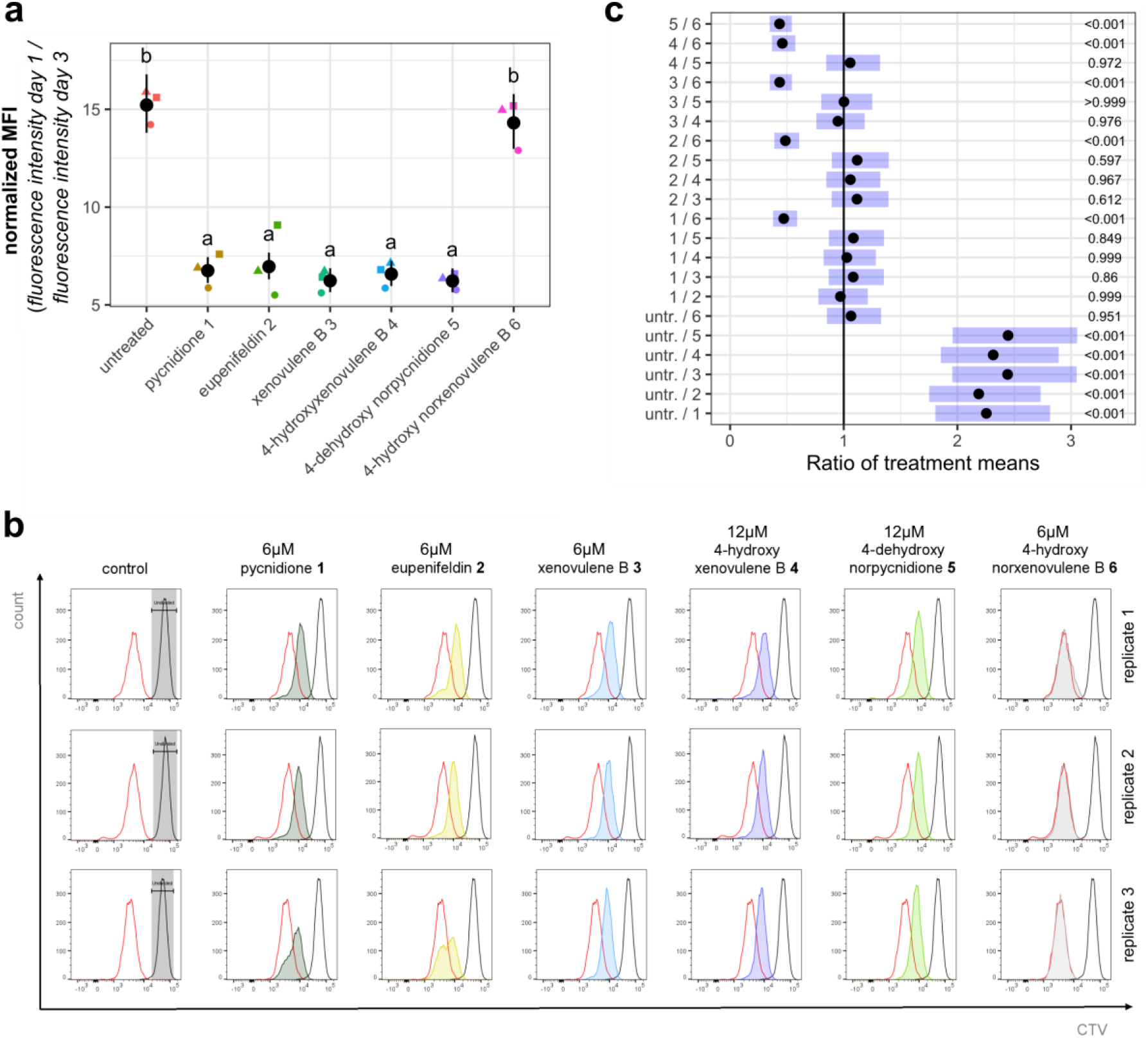
Proliferation decrease of FAIK3-5 cells after treatment with the positive controls pycnidione (**1**) and eupenifeldin (**2**), and with xenovulene B (**3**), 4-hydroxyxenovulene B (**4**) and 4-dehydroxy norpycnidione (**5**) compared to untreated controls and tropolone moiety lacking compound 4-hydroxy norxenovulene B (**6**). Statistics were calculated based on normalized values of obtained ln-transformed mean fluorescence intensities (MFIs) directly after the CellTrace™ Violet (CTV) staining of the undivided population and 24h post compound treatment. **(a)** The graphical overview shows ratios of MFI values obtained on day 1 and day 3 of three biological replicates (●=1, ▴=2, ▪=3) after treatment with compounds **1**-**6** and the untreated control sample, as well as their modelled means (black dots) and pointwise 95% confidence intervals. Treatments marked with the same letter do not differ statistically significantly from each other (α=0.05). **(b)** Histograms and obtained fluorescence intensities of CTV stained FAIK3-5 cells show MFI peaks and values of starting intensities of undivided cell populations at d1 (grey background), control peaks of untreated samples at d3 (red) and MFI peaks of the respective compounds used for treatment (dark green = pycnidione (**1**), yellow = eupenifeldin (**2**), light blue = xenovulene B (**3**), dark blue = 4-hydroxyxenovulene B (**4**), light green = 4-dehydroxy norpycnidione (**5**), 4 grey = 4-hydroxy norxenovulene B (**6**)). **(C)** Pairwise mean comparisons showed statistically significant differences in MFI ratios of day 1/day 3 between all TS compounds treated cells compared to untreated controls and compound 4-hydroxy norxenovulene B (**6**). Black dots indicate back transformed means of calculated ratios and their 95% confidence intervals (blue bars). Respective p-values for each calculation are displayed on the right side. The estimated contrasts and corresponding confidence intervals are given in the statistical supplementary statS7.

To exclude potential solvent effects, we analyzed the effects of DMSO vehicle controls (0.4 % (*v/v)*, 0.6 % (*v/v*), 0.8 % (*v/v*) and 1.2 % (*v/v*)), and 4-hydroxy norxenovulene B (**6**) in the concentrations used for compound treatment (4 µM, 6 µM, 8 µM and 12 µM) on FAIK3-5 cells (**Supplementary Figure S 5**). Statistically significant effects on cell proliferation were observed in samples treated with 1.2 % (*v/v*) DMSO or 8 - 12 µM 4-hydroxy norxenovulene B (**6**) compared to the untreated FAIK3-5 cells, but not between the vehicle control and tropolone lacking compound **6**, that were used as negative controls. However, the fold changes compared to the untreated control were rather small in both the vehicle control (1.09 ± 0.04 after 1.2 % (*v/v*) treatment), and tropolone-lacking compound **6** treated samples (1.05 ± 0.05 after 8 µM treatment and 1.08 ± 0.04 after 12 µM treatment) when compared to samples treated with TS compounds (**Supplementary Figure S 4b**) at the respective concentrations.

To investigate the potential cytotoxicity of compounds **3** - **6** and vehicle controls, their effects on FAIK3-5 cell survival was evaluated via SYTOX™ AADvanced™ staining and compared to the positive controls **1** and **2**, which are known to be cytotoxic ^13, 15–17, 22, 23^. Significances were calculated via pairwise mean comparison analysis on logit scale based on odds ratios calculated from the number of living and dead cells per analyzed sample (sample size=10,000 cells) after 24 h of treatment. These ratios represent the chance to obtain live cells in a specific sample compared to the chance of obtaining dead cells in the respective correlated sample.

To evaluate the effect of negative controls on cell survival, FAIK3-5 cells were treated with different concentrations of the DMSO vehicle control (0.4% (*v/v*) – 1.2% (*v/v*)) and the trolopone lacking compound **6** (4µM – 12µM), as described before. Although the mean comparisons showed significant differences in DMSO and compound **6** treated samples compared to untreated controls, no enhanced effect after treatment with higher concentrations was observed, and the cytotoxic effect was small (**Supplementary Figure S 6**). Comparing the effects of the different analyzed compounds on cytotoxicity, we observed a statistically significant increase in the proportion of dead FAIK3-5 cells upon treatment with the positive controls pycnidione (**1**) (p < 0.001) and eupenifeldin (**2**) (p < 0.001), as well as with xenovulene B (**3**) (p = 0.012) and 4-hydroxyxenovulene B (**4**) (p = 0.004), compared to the untreated controls (**Figure 6**). 4-dehydroxy norpycnidione (**5**) showed no such effects. By investigating the effect on cell death compared to the tropolone lacking compound 4-hydroxy norxenovulene B (**6**) instead of untreated samples, FAIK3-5 cells exhibited a highly increased proportion of dead cells after treatment with the bistropolones **1** and **2** (p < 0.001) that served as positive controls, but not after treatment with the monotropolones **3, 4** and **5**. Thus, the inclusion of a second tropolone moiety to the TS-core structure seemed to have an additional cytotoxic effect, emphasized by the statistically significantly lower amount of detected dead cells in samples treated with xenovulene B (**3**) or dehydroxy norpycnidione (**5**) compared to pycnidione (**1**) (p=0.036 or <0.001, respectively). In summary, four of the tested compounds led to cytotoxic effects which appeared to be stronger when the molecules contained two tropolone moieties.

**Figure 6:**
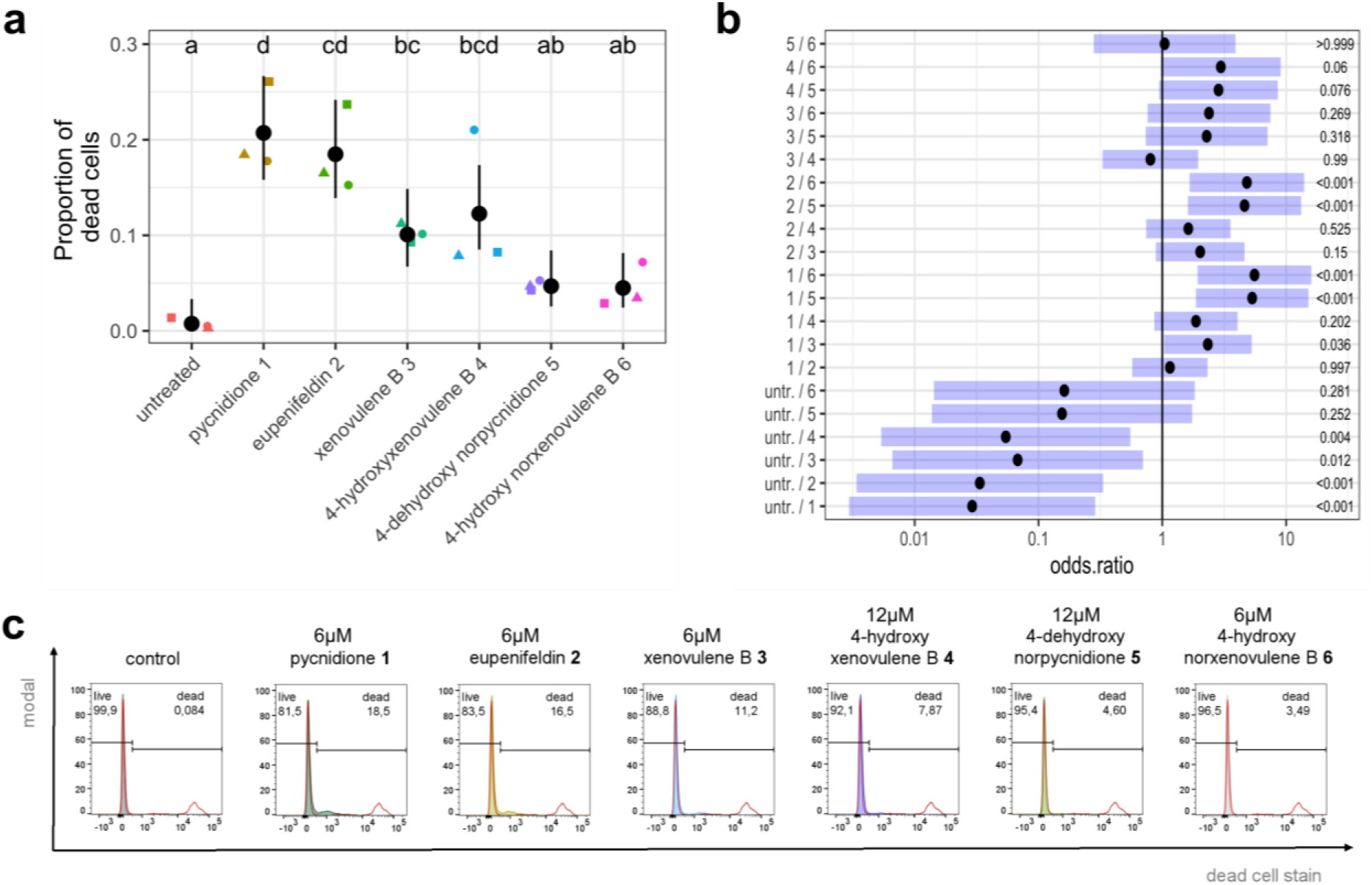
Cytotoxic effects after treatment with 6 µM of the positive controls pycnidione (**1**), eupenifeldin (**2**), and with xenovulene B (**3**), and 12 µM of 4-hydroxyxenovulene B (**4**) and 4-dehydroxy norpycnidione (**5**) on FAIK3-5 cells. As negative controls untreated cells and 6 µM of the tropolone moiety lacking compound 4-hydroxy norxenovulene B (**6**) were used. Statistical analysis was conducted based on the calculation of the average proportion of dead cells per samples (samples size = 10.000 cells). **(a)** Proportions of dead cells (black dots) and their pointwise 95 % confidence intervals (error bars) of three biological replicates (●=1, ▴ = 2, ▪ = 3). Treatments marked with the same letter do not differ statistically significantly from each other (α = 0.05). **(b)** Multiple comparisons of calculated odds ratios (black dots) and simultaneous 95 % confidence intervals (blue bars, details see statistical supplementary statS9) showing differences of cell survival between samples treated with tropolone moiety containing compounds (**1** – **5**) compared to untreated samples or tropolone-lacking compound **6**. P-values for each contrast test of different treatments are displayed on the right side. **(c)** Representative plot of fluorescent intensities of SYTOX™ AADvanced™ stained FAIK3-5 cells samples. Each plot displays percentages of living and dead populations in an untreated control sample and after treatment with the TS compounds pycnidione (**1**), eupenifeldin (**2**), xenovulene B (**3**) and 4-hydroxyxenovulene B (**4**) and 4-dehydroxy norpycnidione (**5**), and tropolone lacking compound 4-hydroxy norxenovulene B (**6**). Peaks of fluorescence intensities are displayed with the respective living / dead control (red) containing a heat-killed dead cell population.

## DISCUSSION

This study examines the effect of TS compound treatment to detect bioactivities of the investigated natural and unnatural newly discovered TS. Here, we report the first bioactivities for xenovulene B (**3**), 4-hydroxyxenovulene B (**4**) and 4-dehydroxy norpycnidione (**5**) in terms of cytotoxicity and effects on proliferation and cell morphology. The bioactivity of the compounds appeared to be highly dependent on the cellular model applied, as evident from EC_50_ estimations in metabolic activity assays in PC-3, Jurkat, MDA-MB-231 and FAIK3-5 cells.

The bioactivities of the tested TS compounds, were also dependent on the tropolone-moieties present, and showed a tendency to be enhanced in compounds with a tropolone dual– substitution compared to monotropolones. The results support previous research showing that methylated pycnidione derivatives lack activity in causing EPO gene expression in hepatic cell lines ^20^ and enhancing bleomycin treatment effects in Jurkat cells ^16^. This indicates that biological activity of the compounds depends on the presence of at least one unprotected tropolone moiety although the exact structure-activity relationship remains unclear.

### Half maximal effective concentration (EC_50_)

Estimations of EC_50_ values for all the investigated cell lines were spread in a wide range of concentrations in FAIK3-5, PC-3 and Jurkat cells, with a lack of response to TS treatment observed in MDA-MB-231 breast cancer cells. Given the possibility that TS compounds may act on cells through multiple unknown mechanisms, differences in the pathways affected between different cancer and non-cancer cell lines, which vary in their genetic landscape, may result in distinct sensitivities to the compounds used. For example, the malignant transformation that cancer cells must undergo to become cancerous, is a complex process involving genetic alterations, that influence cancer development and lead to clonal variations between established cancer cell lines^51^. This process in turn affects protein function and DNA methylation patterns, which influence the cell’s response to chemical compounds ^36^. In addition, MDA-MB-231 cells, known to display epithelial to mesenchymal transition (EMT) associated with breast cancer metastasis and multi-drug-resistance to anti-cancer reagents like cisplatin ^52^, doxorubicin ^53^ and paclitaxel ^54^, show a primary resistance to all of the applied TS compounds, eventually by suppressing apoptotic pathways and increasing of breast tumor stemness ^49, 50^.

### EPO induction

Besides their influence on cell viability, cell proliferation and adhesion, TS compounds were previously shown to influence EPO induction in several reporter cell lines ^20, 21^. In the current study, the nonconditionally immortalized murine REP-dervied cell line FAIK3-5 ^39^ was chosen as a model for natural EPO production *in vitro*, because, unlike previously used engineered reporter cell lines, it can produce EPO by itself at a basal level and thus more closely resembles physiological conditions. While this has the advantage of reducing assay interferences and providing more reliable results, it also exposes the model to naturally occurring regulatory mechanisms such as hypermethylation of regulatory gene sequences, thereby putatively inhibiting EPO production within the cells ^37^. Previous reports of EPO induction by TS compounds of the "epolone family" in reporter cell lines of Hep3B ^20^, HepG2, and HRCHO5 ^21^ described that certain TS compounds are able to induce EPO synthesis independent of oxygen availability, *e.g*. by inhibiting proteins containing a HIF-prolyl hydroxylase (PHD) domain. Therefore, we hypothesized that the tested TS compounds might also be capable of oxygen-independent HIF1α stabilization ^55^. In the model cell line used, however, EPO production could not be induced by external stimuli putatively stabilizing HIF1ɑ. In contrast, treatment with the DNA methyltransferase inhibitor 5-azacytidine increased basal EPO levels by approximately 2-fold in the positive controls pycnidione **1** and eupenifeldin **2**, as well as in the structurally related compounds (**3**–**6**), irrespective of the presence of a tropolone moiety, and in the vehicle controls tested. These results support the assumption that EPO production is tightly regulated at the transcriptional level, with DNA methylation of the EPO promoter partially involved in epigenetic inactivation of EPO expression ^43^. This mechanism was also previously observed in the REP cell line used, and the authors proposed the occurrence of an intrinsic transcriptional negative feedback loop responsible for the decline in EPO production within the cells, or by their putative differentiation into myofibroblasts upon prolonged culture ^39, 43, 56^.

### Cell proliferation and morphology

We found that treatment with all tested tropolone bearing compounds (**1**–**5**) resulted in a statistically significant inhibition of cell proliferation, regardless of the single or dual substitution of tropolone moieties. This was accompanied by a strong morphological alteration in the tested cell model. Cell morphology is influenced by dynamic interactions between cytoskeletal, membrane and adhesion complexes and regulatory signal transduction systems that translate external cues into biochemical signals for cell adaptation ^57^. Cell phenotypes are therefore a combination of observable characteristics with biochemical properties and the expression of genes and proteins. For example, the emergence of an enlarged, flat and multinucleated phenotype which is highly granulated can reflect senescent cell stages *in vitro* ^58^. In this study, the appearance of large and highly granular cells and a telocyte phenotype was observed in all TS-treated samples, either due to direct action of the tested compounds or indirectly via proliferative regulation and subsequent lower cell densities that are known to drive FAIK3-5 cells to evolve a telocyte phenotype ^39^. Moreover, closely related compounds induced similar phenotypic changes in treated FAIK3-5 cells. Specifically, three groups of distinct phenotypes could be identified, which could be connected to treatment with either bistropolones (**1**, **2**), monotropolones (**3**, **4**, **5**) or chemicals without tropolone moieties (DMSO control, **6**). This finding suggests that the three groups of compounds likely target similar biological pathways via specific receptors, resulting in consistent yet distinct biological responses reflected in cell morphology.

### Cytotoxicity

In previous studies Hsiao et al. and Kaneko et al. found that pycnidione **1** affects cell cycle and cell proliferation directly via cell cycle regulatory proteins, with apoptosis and reduced cell adhesion observed ^16, 17^. In our study, the treatment with all tropolone-bearing compounds (**3**-**5**) and their positive controls (**1**, **2**) also reduced cell proliferation and exhibited cytotoxic effects with observable detachment of cells, with the bistropolone positive controls pycnidione (**1**) and eupenifeldin (**2**) showing significantly higher cytotoxic potency than the tested monotropolones xenovulene B (**3**) and 4-hydroxyxenovulene B (**4**), and low cytotoxicity of 5-dehydroxy norpycnidione (**5**). Thus, we concluded that the tropolone moiety is primarily responsible for the observed cytotoxic effects, while hydroxylation of the macrocyclic core structure affects compound efficacy additionally, which is supported by previous studies on TS toxicity demonstrating higher cytotoxic potency of the tested bistropolone eupenifeldin (**2**) compared to its dehydroxy analogue dehyroxyeupenifeldin, specific tropolone-lacking meroterpenoids, as well the monotropolone noreupenifeldin B ^59^. Additionally, despite its universal use in cryopreservation or as a solvent for poorly soluble molecules, there is evidence of DMSO toxicity at low doses and its impacts on the epigenetic landscape of treated samples ^41, 60^. In line with these reports, we also observed statistically significant effects of DMSO on proliferation and cell viability, although the observed inhibition of proliferation and cytotoxicity were considerably lower than in samples treated with TS compound **1** – **5**.

## Conclusion

In conclusion, this study reveals the bioactive properties of xenovulene B (**3**), 4-hydroxyxenovulene B (**4**), and 4-dehydroxy norpycnidione (**5**), in terms of proliferation inhibition and cytotoxicity and shows a significant correlation between tropolone substitutions and cell viability. The monotropolone compound 4-dehydroxy norpycnidione (**5**) showed an improved proliferation inhibition compared to the tropolone-lacking compound 4-hydroxy norxenovulene B (**6**) and untreated control samples. However, at the same time cytotoxic effects exerted on the model cell line were lower than those of the used positive controls pycnidione (**1**) or eupenifeldin **c**(**2**). The reduced toxicity, concomitant severe decrease in proliferation and low EC_50_ values estimated by inhibition of metabolic activity induced by 4-dehydroxy norpycnidione (**5**) could be a sign of cell transition to a senescent stage or non-senescent growth arrest such as cellular quiescence. As the exact mechanism of action of these highly interesting specialized metabolites is still unknown, future investigation of their precise dose-response relationships in different cell models will help to elucidate their putative value as drug models for subsequent preclinical studies.

### Limitations of the study

There are several challenges in studying physiological EPO production *in vitro*, as there is currently no appropriate cell model for REP cells. Cultivatable immortalized murine FAIK3-5 and Replic cells have been reported to undergo myofibroblast formation and epigenetic regulation in culture ^39, 43^ associated with loss of EPO production. Consistent with this, and despite the observed effects of TS compounds on cell morphology, proliferation and cytotoxicity, the controls pycnidione **1** and eupenifeldin **2** do not show any EPO induction in the cell model used. Due to the complex and partially unknown EPO regulatory mechanism in the FAIK3-5 cell line and its lack of response to external HIF1ɑ stabilizing stimuli, we cannot make any further conclusions about the bioactive compounds (**3**-**5**) regarding their ability to increase EPO protein production.

After verifying the bioactivities of xenovulene B (**3**), 4-hydroxyxenovulene B (**4**), and 4-dehydroxy norpycnidione (**5**) in FAIK3-5, PC-3 and Jurkat cells, the next step should be to test the active compounds in a broader panel of cells, including renal (cancer)-cell lines, primary cells and hepatic models to estimate drug metabolism. Future directions in the field may involve toxicity testing in organoids and 3D models to detect possible mechanism of action and unexpected effects of TS compound treatment.

## STAR METHODS

### RESOURCE AVAILABILITY

#### Lead contact

Further information and requests for resources and information should be directed and will be fulfilled by the lead contact, Prof. Dr. Cornelia Lee-Thedieck

### Materials availability

This study did not generate new unique materials.

### Data and code availability

The R-codes generated in this study are available in the provided statistical Supplementary. Any additional information required to reanalyze the data reported in this paper is available from the lead contact upon request.

## EXPERIMENTAL MODEL AND STUDY PARTICIPANT DETAILS

### Cell lines

Bioactivitiy of TS compounds **1** – **6** was tested *in vitro* on the cancer cell lines MDA – MB – 231 (RRID:CVCL_0062), PC – 3 (RRID:CVCL_0035) and Jurkat (RRID:CVCL_0065) cell lines, and the non – cancerous fibroblastoid atypical interstitial kidney cell line FAIK3 – 5 (RRID:CVCL_A5EC). All cells were obtained from the Leibniz Institute DSMZ-German Collection of Microorganisms and Cell Cultures GmbH (Braunschweig, Germany) and were cultured according to the recommended culture protocol in Dulbecco′s Modified Eagle′s Medium (DMEM) - high Glucose (Sigma Aldrich, St. Louis, MO, United States) (MDA – MB – 231, PC – 3 and FAIK3 – 5 cell line), or in Roswell Park Memorial Institute (RPMI) 1640 medium (Sigma Aldrich, St. Louis, MO, United States) (Jurkat cell line), each supplemented with 10 % h.i. FBS (Sigma Aldrich, St. Louis, MO, United States; Lot: BCBW9923), hereafter referred to as cell culture medium. Adherent cells were split every 2^nd^ to 3^rd^ day at a confluency of 80 – 90 %, and cells in suspension were passaged every 2^nd^ to 3^rd^ day to a cell density of 1 × 10^5^ viable cells / mL. As the FAIK3 – 5 cell line was previously described to produce EPO protein for at least 30 passages ^39^, all experiments were conducted on cells below that passage number. To investigate hypoxia- independent effects of the tested TS compounds **1** – **6** on FAIK3 – 5 cells, all experiments were conducted under normoxic conditions at 37 °C and 5 % CO_2_ under saturating humidity in a CO_2_ incubator:

## METHODS DETAILS

### Biosynthesis of tropolone-sesquiterpenoid compounds

Pycnidione **1** and eupenifeldin **2** were isolated respectively from *Leptobacillium* sp. CF-236968 and *c* (formerly referred to as unidentified ascomycete F-150626) that naturally produce the compounds ^27^. For the heterologous production of the described tropolone sesquiterpenoid (TS) compounds **3 - 5** and control compound 4-hydroxy norxenovulene B **6**, a series of TS key biosynthetic gene clusters (BGC) were refactored and recombined in *A. oryzae* NSAR1 as previously described by Schotte et. al ^27^. The currently accepted stereochemistry of these compounds is displayed in **Figure 1** ^24^.

### Compound reconstitution and TS treatment of cell lines

Lyophilized TS compounds were reconstituted in dimethyl sulfoxide (DMSO, Carl Roth GmbH, Karlsruhe, Germany) to obtain 1 mM stock solutions, sterile filtered and stored at -20 °C until further use. To maintain compound stability, repeated freeze-thaw cycles of compound aliquots were avoided. Prior to treatment, aliquots were thawed at RT under light exclusion and spun down briefly. Depending on the assay, cells were seeded at the appropriate cell number as described in the following sections and allowed to adhere ON before visual inspection of cell adherence and morphology. Prior to TS treatment, cells were thoroughly washed with Dulbecco’s phosphate buffered saline (DPBS) (Sigma Aldrich, St. Louis, MO, United States) and incubated with cell culture medium containing TS compound at the appropriate concentrations for 24 hours. DMSO vehicle controls and untreated medium controls were included for each analysis.

### Measurement of metabolic activity

The half maximum effective concentration (EC_50_) of natural and unnatural TS compounds was evaluated after treatment of FAIK3-5, MDA-MB-231, PC-3, and Jurkat cells with compounds **1** – **6** using the CellTiter - Blue Cell® Viability Assay (Promega, Madison, WI, USA). The ability of living cells to convert the redox dye resazurin into the fluorescent end product resorufin, which can be detected at 590 nm, was used to monitor the metabolic activity of the cells and thus determine the viability of the treated cell lines.

The cell lines in use were seeded onto 96 well plates (F-Bottom 96-well Assay Microplates, Greiner Bio - One, Kremsmünster, Austria) in 75 µL cell culture medium at a density of 1 × 10^4^ cells per well. The cells were then cultured for recovery and adhesion ON as described before. Thereafter, cells were subjected to TS compounds **1** - **6** in a concentration series of 0.001 µM, 0.01 µM, 0.1 µM, 1 µM, 10 µM, 25 µM and 75 µM in triplicates for 24 hours. Therefore, for each well the double concentrated compounds were prepared in 75 µL of the respective cell culture medium and added to the cultured cells. Vehicle controls were prepared for each cell line and compound concentration, using the following DMSO concentrations: 0.0001 % (v/v), 0.001 % (v/v), 0.01 % (v/v), 0.1 % (v/v), 1 % (v/v), 2.5 % (v/v) and 7.5 % (v/v). In addition, untreated controls and dead controls (10 % (v/v) and 25 % (v/v) DMSO) were included. Medium and vehicle controls without cells were used for each experimental setup (N = 3) and cell line to assess background signals. After incubation, 30 µL of CellTiter - Blue Cell® Reagent was added to each well, and the cells were incubated for a further 3 hours at 37 °C before measurement on an Infinite® 200 PRO Plate reader (Tecan Austria GmbH, Groedig, Austria). The plate was automatically shaken for 10 seconds at an amplitude of 6 nm before fluorescence measurement at 590 nm. For comparison purposes the gain was set manually at 100 for each plate and experimental repetition. Statistical analysis and calculations of EC_50_ values for each cell line and compound is described in more detail in the statistics section below.

### Evaluation of optimal compound concentrations for bioactivity testing in FAIK3 – 5 cells

In preliminary tests on cell proliferation and EPO induction, optimal compound concentrations were determined by treating FAIK3-5 samples with compounds **1** – **6** at concentrations of 0 µM, 4 µM, 6 µM and 12 µM (Supplementary Figure **S 2** – **S 4**). This concentration range was based on previous studies by Cai et al. ^20^, in which EPO production was induced with 10 µM of the positive control pycnidione **1**, and by Wanner et al. ^21^ who have determined that the optimal concentration for bioactivity of pycnidione **1** is in the range of 4 µM to 8 µM under normoxic conditions. As a proof of principle for the different bioactivities of the investigated compounds on FAIK3 – 5 cells, and based on preliminary experiments, in which compound **4** and **5** showed highest effects at 12 µM (Supplementary Figure **S 3** and **S 4**), we decided to use the highest possible effective compound concentration at which the cells also show a reasonable viability for further testing (Supplementary Figure **S 6 e**). Therefore, compounds **1**, **2**, **3** and **6** were used at a concentration of 6 µM, while compounds **4** and **5** were used at a concentration of 12 µM to test EPO levels, proliferation, cytotoxicity and morphology on FAIK3 – 5 cells.

### Analysis of erythropoietin production via Enzyme-linked Immunosorbent Assay

In previous studies an enhancing effect on the EPO production of pycnidione **1** and structurally related epolones was described for Hep3B and HRCHO5 reporter cell lines ^20, 21^. To analyze, if the utilized compounds are also capable to induce a similar effect on the natural EPO production of REP - derived FAIK3 - 5 cells *in vitro*, the EPO protein content of TS compound treated FAIK3 - 5 derived cell extracts was measured via enzyme - linked immunosorbent (ELISA) assays, using the RayBio® Mouse EPO ELISA Kit (Raybiotech, Peachtree Corners, GA, United States) according to the manufacturer’s protocol.

FAIK3 – 5 cells were seeded into 6 well plates (CELLSTAR® Cell culture 6 – well plate, Greiner Bio - One, Kremsmünster, Austria) at a cell density of 5 × 10 ⁵ cells / well and kept in culture for 24 hours. The cells were then treated for 24 hours as described above, using 6 µM and 12 µM of TS compounds, and including respective vehicle controls (0.6 % *v/v* and 1.2 % *v/v* DMSO) and untreated samples. It is known that the physiological EPO production is tightly controlled by an “on - off” regulatory behavior of REP cells and by DNA methylation in the *EPO* - gene promoter that can cause the loss of EPO production capability in REP cells ^40^. Therefore, FAIK3-5 were treated with 1 µM of the hypomethylating agent 5 - Azacytidine (5 - Aza, Sigma-Aldrich, St. Louis, MO, United States) in an additional approach. The treatment was conducted for six days with complete medium changes every other day to ensure sufficient cell divisions for 5 - Aza integration and agent replenishment before degradation. After this period, 5 – Aza pre – treated cells were exposed to 6 µM or 12 µM of TS compounds, DMSO at a concentration of 0.6 % v/v and 1.2 % v/v DMSO) or left untreated for 24 hours.

Cell extracts were obtained after visual examination of growth density, by washing the samples twice in ice - cold DPBS and followed by disruption in RIPA buffer (ThermoFisher Scientific, Waltham, MA, United States) supplemented with SIGMAFAST™ Protease Inhibitor Tablets (Sigma Aldrich, St. Louis, MO, United States) to prevent protein degradation. Quantification of total protein concentrations was conducted with the Pierce™ BCA Protein Assay Kit (ThermoFisher Scientific, Waltham, MA, United States) and samples were diluted to a total protein concentration of 200 µg / mL. Finally, EPO - ELISA analysis was conducted and absorbance at 450 nm was measured immediately with the Infinite® 200 PRO Plate reader (Tecan Austria GmbH, Groedig, Austria). Experiments were analyzed with the provided Excel based freeware ELISA analysis tool v1.0 (Raybiotech, Peachtree Corners, GA, United States) and statistical analysis was performed as described in the respective section below.

### Morphological analysis

In order to handle environmental changes, cells possess the ability to attain a more desirable stress state by the activation of distinct molecular pathways, which morphologically might lead to profound changes of cell shapes. To analyze different morphological shapes of FAIK3 - 5 cells caused by TS compound treatment, brightfield pictures were taken with a ZEISS Axio Vert.A1 Inverted Microscope (Zeiss, Oberkochen, Germany) 24 hours after TS treatment and directly before subsequent analysis of the cells.

To avoid human bias and increase reproducibility of the analysis, cell segmentation of the images was performed using *Cellpose* ^48^, a generalist deep learning - based algorithm, trained to accurately recognize different cell shapes. A region of interest (ROI) was created in relation to each cell and then transferred to Fiji (ImageJ, Version 1.53c bundled with Java 1.8.0_172) ^61^ for subsequent image analysis. Images were analyzed for morphological changes for each treatment condition, which included treatment with the tested compounds **1** - **6** and an untreated control sample to replicate the natural morphology of the FAIK3-5 cell line (N=3 independent experiments for each condition). To obtain the most accurate representation of the different cell shapes in each image, all cells were included in one image, with ROIs ranging from n=194 to n=1,773 cells per image and a total of n=16,632 cells from the different test conditions, that were analyzed and compared. Subsequently, ROIs were measured in Fiji for basic morphometric variables (cell area, perimeter, major and minor axis) and standard shape descriptors (circularity, aspect ratio, roundness and solidity). To analyze differences in ramification patterns of cells treated with distinct TS compounds two-dimensional branching descriptors were calculated using the *AnalyzeSkeleton* plugin ^62^. The results were visualized using the Matlab (Version R2021b) toolbox *PhenoPlot* ^47^, an open source visualization tool for imaging data that is able to represent multiple dimensions of cellular morphology as intuitive cell-like glyphs. For a detailed description of morphological analysis and *PhenoPlot* visualization see Supplementary Figure **S 7**.

### Proliferation and Cytotoxicity Assay

To evaluate effects of proliferative behavior and cell viability caused by TS compounds, FAIK3-5 cells were cultured in cell culture medium as described above and labelled with the CellTrace™ Violet (CTV) Cell Proliferation Kit (Invitrogen, ThermoFisher Scientific, Waltham, MA, United States) at a concentration of 10 µM CTV per 1 × 10 ⁶ cells according to the manufacturer’s protocol with minor modifications. To assure homogenous CTV labelling, 900 µL of cell suspension containing 1 × 10 ⁶ cells were kept in a 15 mL tube (Greiner Bio-One, Kremsmünster, Austria), which was positioned horizontally before 100 µL of CTV staining solution was added to the upper end of the tube without mixing. The tube was kept in that position until the lid was closed carefully, and mixing was conducted on a vortex mixer (Benchmark Scientific, Sayreville, NJ, United States). After 20 minutes of staining at 37 °C in the dark, the reaction was stopped by adding 5 times the original staining volume of DPBS with 10 % of FBS and incubation for 5 minutes on ice to remove unbound dye particles. Subsequently, the supernatant staining solution was removed after centrifugation at 300 × g for 5 minutes, and FAIK3 - 5 cells were counted and seeded into 6 well plates with a cell density of 5 × 10⁵ cells per well. Cell viability was evaluated via SYTOX™ AADvanced™ Dead Cell Stain Kit (Invitrogen, ThermoFisher Scientific, Waltham, MA, United States) staining according to the manufacturer’s instructions prior to analysis with the BD FACSverse^TM^ Cell Analyzer (BD, Franklin Lakes, NJ, United States) cytometer.

Directly after staining, cell samples were analyzed with the flow cytometer to ensure homogenous dye distribution within the population and to monitor cell viability after the staining process. For each analyzed sample 10,000 events were counted. Gating strategies for living / dead discrimination and proliferation analysis are explained in more detail in the Supporting Information section (Supplementary Figure **S 8**). For each analysis unstained cells, CTV - stained cells without compound treatment and live / dead control samples, comprising a heat - killed population to set the fluorescence threshold for live / dead discrimination, were included as staining controls. To assess the optimal analysis period for the CTV proliferation tests, untreated FAIK3 - 5 cells were analyzed via flow cytometry for CTV intensities and putative effects of the CTV staining on cell viability on three subsequent days in preliminary experiments.

For following experiments with compound treated samples, measurements were taken at day 1 (d1) to obtain the basic CTV intensities of the stained samples. At day 2 TS compound treatment was started and at day 3 (d3), after 24 hours of incubation, the second measurement was conducted to evaluate changes in CTV intensities after TS treatment. For each experiment untreated cells and vehicle controls (0.6 % v/v and 1.2 % v/v DMSO) were included as treatment control samples. After cellular uptake by diffusion, CTV covalently binds to intracellular amines and is diluted proliferation dependently within the analyzed cellular populations. However, as the cellular uptake and binding capacity is contingent on multiple factors (e.g., cell viability, division stage or cellular protein level), a perfectly homogenous staining throughout experimental repetitions is not possible and hence result in distinct mean fluorescence intensities (MFI) at d1 of the experiments. Accordingly, for statistical analysis obtained values were normalized by calculating the proportions of measured MFIs at the beginning (d1) and end (d3) of the experiments and pairwise mean comparison tests were run based on mean differences, as described in more detail in the statistics section below. For descriptive statistics mean values and standard deviations were obtained from the calculated fold changes enlisted in Supplementary Figure **S 5 e** and **S 6 e**.

## QUANTIFICATION AND STATISTICAL ANALYSIS

### Statistical analysis

The statistical analysis was done based on R version 4.3.1 ^63^ on a 2020 MacBook Air using the aarch64-apple-darwin20 (64-bit) platform. Data management and plotting was done using the tidyverse packages ^64^. EC_50_s were estimated based on hierarchical nonlinear regression models fit with medrm() from the medrc package ^65^. Dunnett type comparisons between the EC50 of DMSO against the EC50s of the compounts were performed based on the multcomp package ^66^. For all other analysis, linear models, generalized linear models or ninear mixed models were used (depending on the scale of the endpoinds and the experimental design). Linear and generalized liner models were fit using the lm() and glm() functions from the stats package ^67^. Mixed effects models were fitted using lmer() from the lme4 package ^68^. Based on the fitted models either an ANOVA (for linear models) or an analysis of deviance ^69^ were run. For each of the fitted mixed effects models, a type III ANOVA was computed using Kenward-Rogeŕs approximation of degrees of freedom ^70^. Subsequently, simultaneous mean comparisons were done using the packages emmeans ^71^ and multcpomp ^66^. In order to account for the multiple-testing problem, all mean comparisons (of which most correspond to Tukeys all pairwise comparisons) were adjusted based on the multivariate t - distribution ^66^. Since the endpoints regarding the cell morphology are highly correlated (several endpoints per cell), simultaneous mean comparisons were run based on multiple marginal models ^72, 73^. An overview of the statistical analysis and the R functions used for modelling and comparing multiple means with reference to the specific part of the statistical supplementary is given in **Table 1**.

**Table 1:**
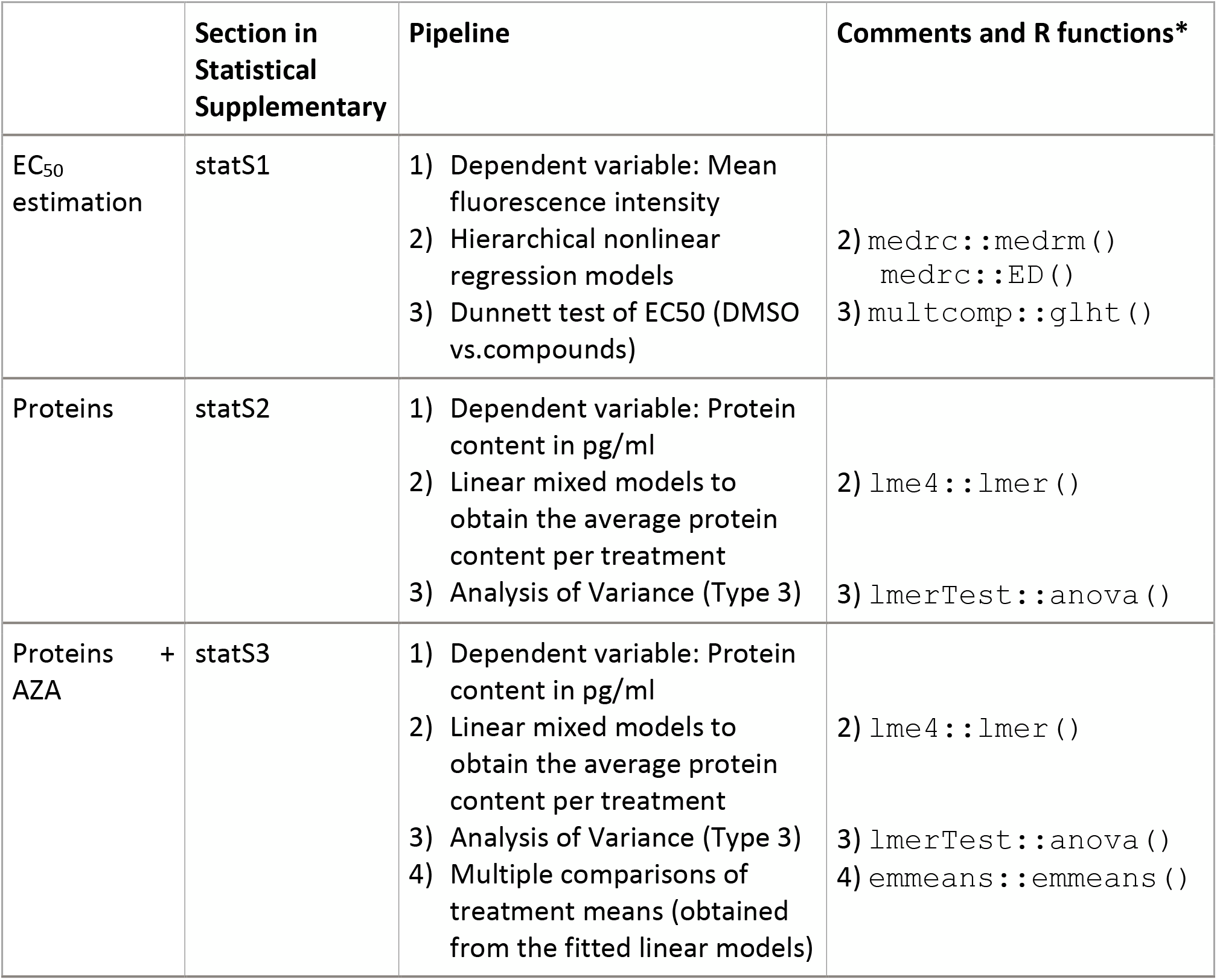
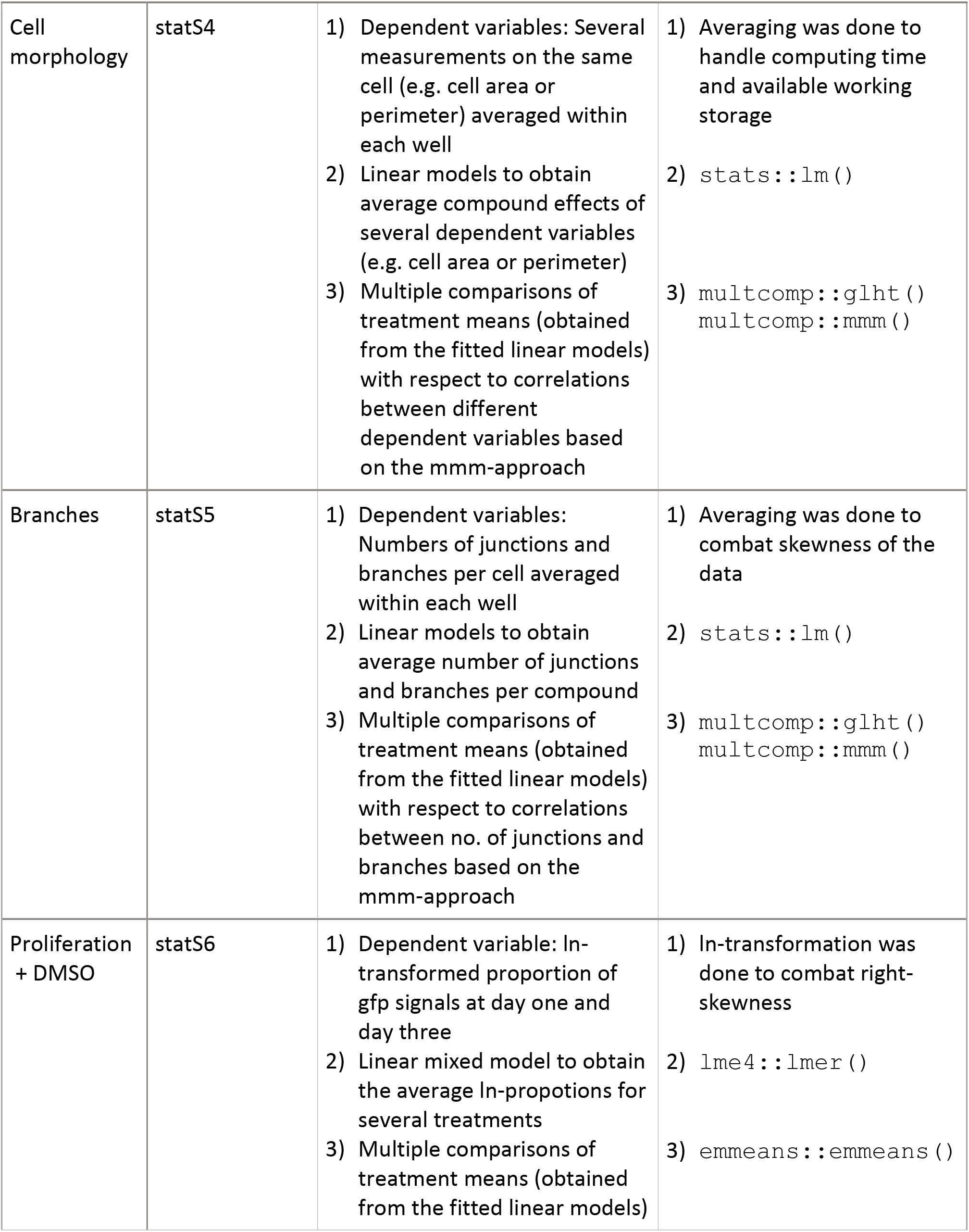
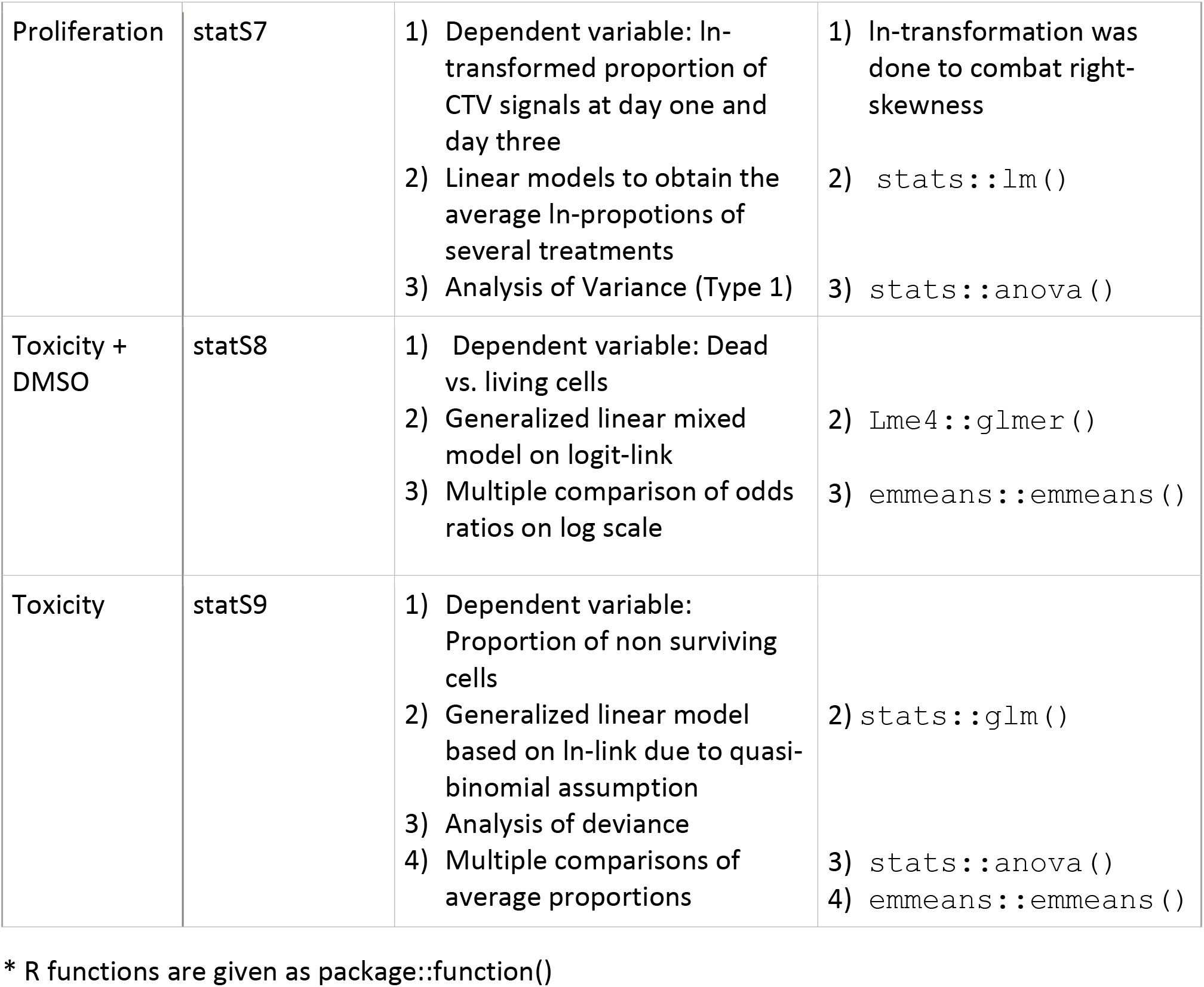
Overview of the statistical analysis.

## METHODS DETAILS

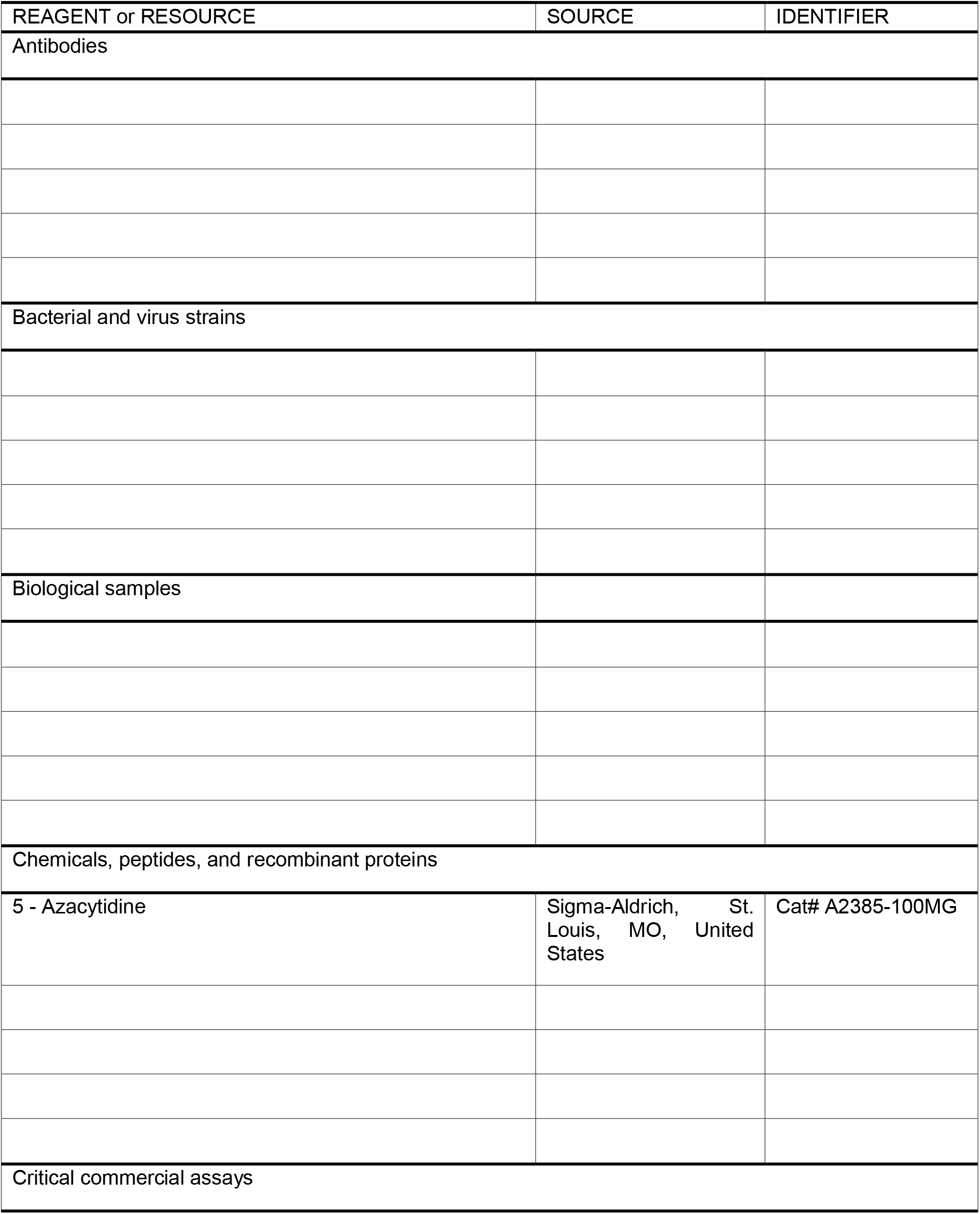
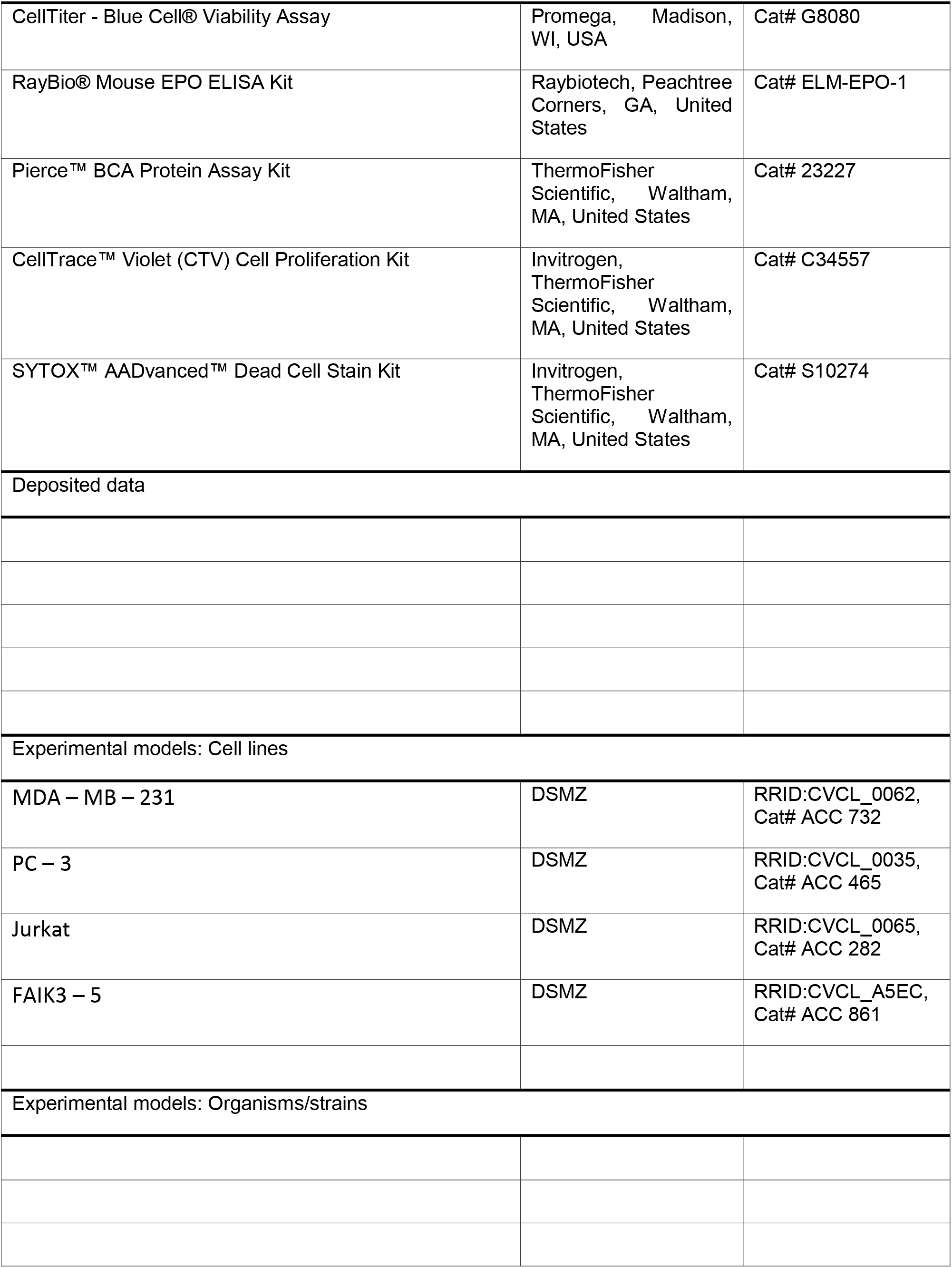
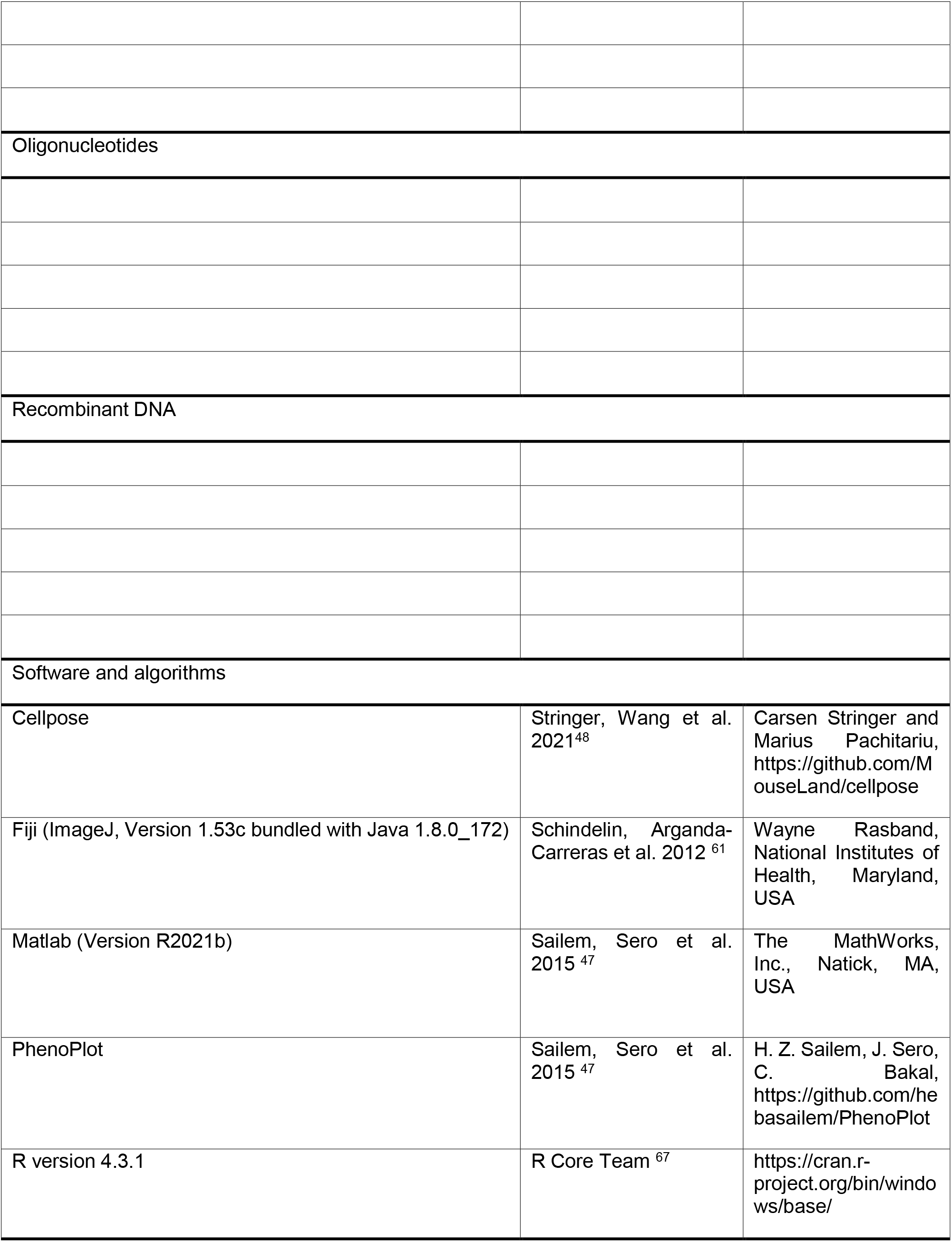
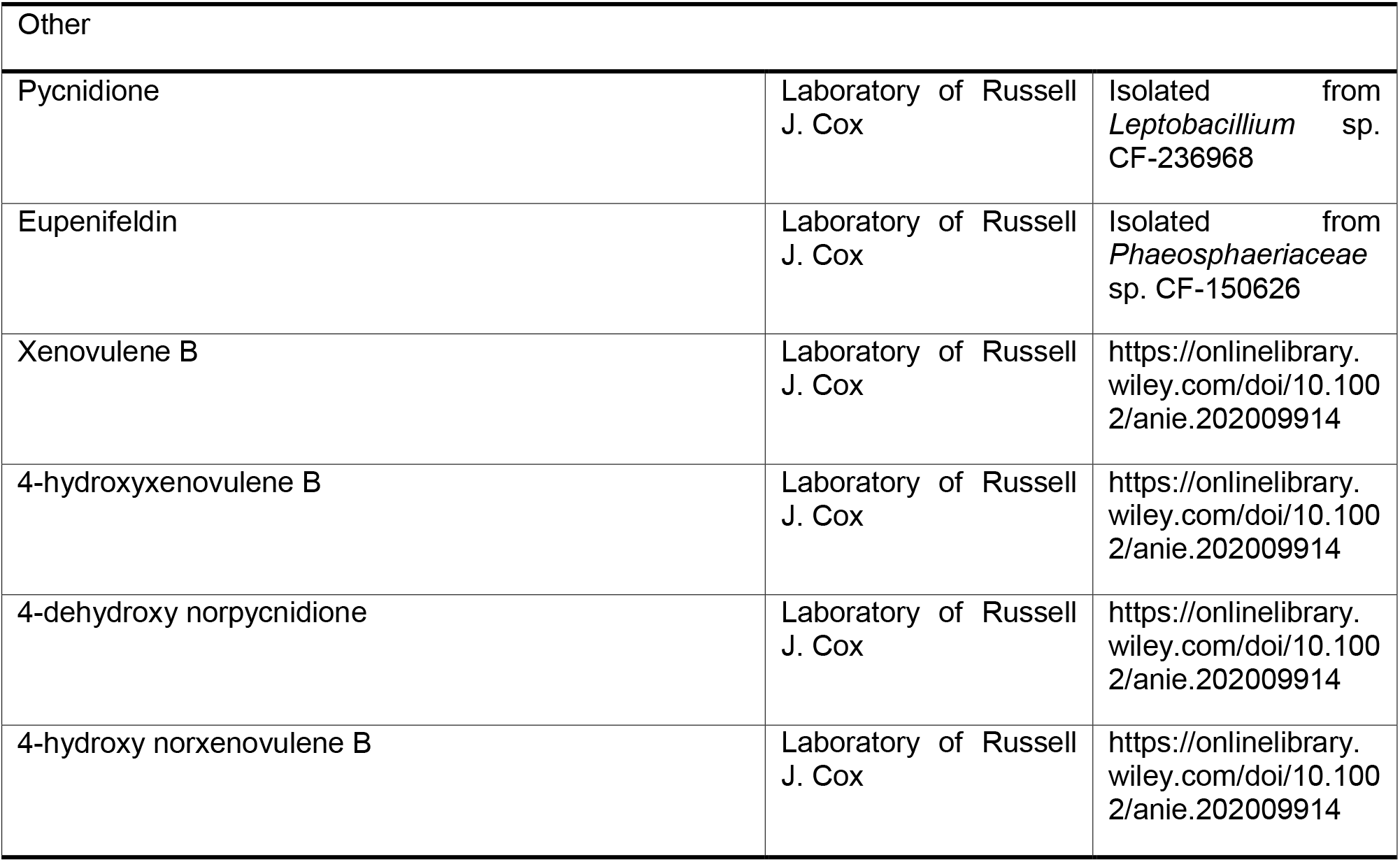

## Supporting information

Statistical Supplementary Information

## ACKNOWLEDGMENT

We would like to thank Frank Schaarschmidt for his valuable input and support in the biostatistics analysis of the experiments.

## AUTHOR CONTRIBUTIONS

T.C.B.: Investigation (lead), Methodology (lead), Formal Analysis (equal), Visualization (lead), Writing – original draft (lead). M.M.: Formal Analysis (lead), Visualization (equal), Writing – Review and Editing (equal). C.S.: Resources (lead), Writing – Review and Editing (equal). R.J.C.: Conceptualization (equal), Resources (equal), Writing – Review and Editing (equal). C.L.T.: Conceptualization (lead), Funding Acquisition (lead), Project Administration (lead), Supervision (lead), Writing – Review and Editing (lead)

## FUNDING SOURCES

This work has been carried out within the framework of the SMART BIOTECS alliance between the Technische Universität Braunschweig and the Leibniz Universität Hannover. This initiative is supported by the Ministry of Science and Culture (MWK) of Lower Saxony, Germany. This project has also received funding from the European Research Council (ERC) under the European Union’s Horizon 2020 research and innovation programme (grant agreement No 757490).

## COMPETING INTERESTS

The authors declare no competing interests.

## ABBREVIATIONS

TS: tropolone sesquiterpenoids
EPO: erythropoiestin
CTV: CellTrace™ Violet
MFI: mean fluorescence intensity

## Notes

### Competing Interest Statement

The authors have declared no competing interest.

### Summary of Updates

The study was updated with analyses on EC50 values of the tested compounds in several cell lines. The presentation of the study including text and figures was updated accordingly.

