## Supplementary Information for "Evaluating the bioactivities of natural and synthetic fungal tropolone sesquiterpenoids in various cell lines"

Dr. M. Menssen

[2] Institute of Cell Biology and Biophysics, Department of Biostatistics, Leibniz University Hannover, 30419 Hannover, Germany

Dr. C. Schotte, Prof. Dr. R.J.Cox

[3] Institute of Organic Chemistry and BMWZ, Leibniz University Hannover, 30167 Hannover, Germany

* Prof. Dr. Cornelia Lee-Thedieck, Leibniz University Hannover, Institute of Cell Biology and Biophysics, Herrenhäuserstr 2, 30419 Hannover. Phone: +49 511 762 14 972; Fax: +49 511 762 2606;

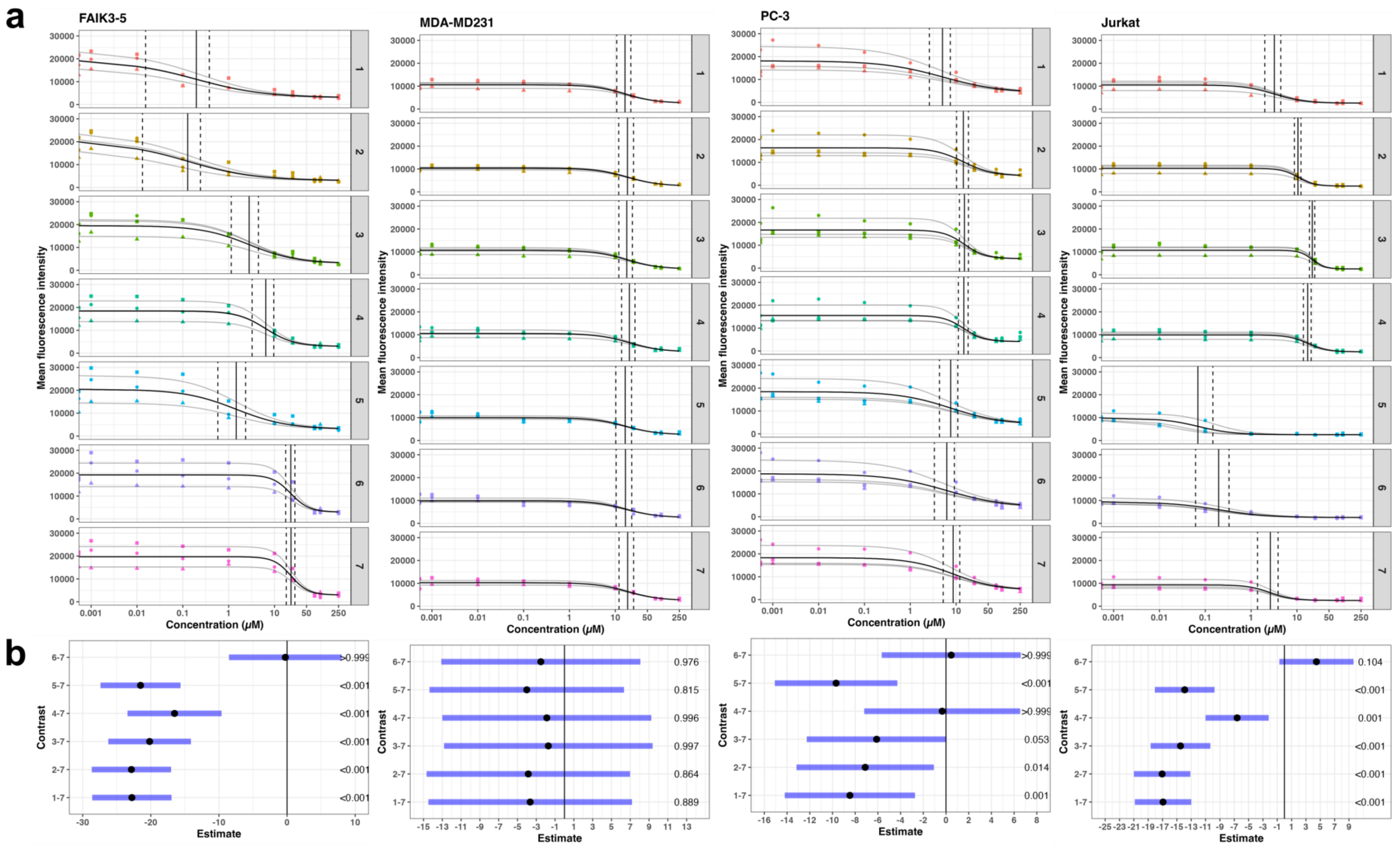


**Figure S 1. (Legend next page)**

**Figure S 1.** $EC_{50}$ of murine FAIK3–5 cells and the cancer cell lines MDA–MB–231, PC–3 and Jurkat after treatment with 0.001 µM, 0.001 µM, 0.1 µM, 1 µM, 10 µM, 25 µM and 75 µM of the tropolone sesquiterpenoids (TS) pycnidione (**1**), eupenifeldin (**2**), xenovulene B (**3**), 4-hydroxyxenovulene B (**4**), 4-dehydroxy norpycnidione (**5**) and tropolone-lacking compound 4-hydroxy norxenovulene B (**6**) and 0.0001 % (v/v), 0.001 % (v/v), 0.01 % (v/v), 0.1 % (v/v), 1 % (v/v), 2.5 % (v/v) and 7.5 % (v/v) of their respective DMSO vehicle controls (7). **(a)** For each experimental repetition one according dose–response curve (grey) was modelled, resulting in the final mean dose-response curve (black) from which $EC_{50}$ values (black vertical line) and the corresponding upper and lower confidence intervals (black dotted vertical lines) were derived. Mean fluorescence intensities were measured in three biological replicates (●=1, ▲=2, ■=3) deriving from the mean values of three technical replicates per experimental repetition. To accurately estimate the lower asymptote of each dose-response curve, dead controls without TS compounds were included comprising 10 % (v/v) and 25 % (v/v) DMSO. **(b)** Pairwise mean comparisons have been calculated to show statistically significant differences between calculated $EC_{50}$ of DMSO vehicle controls and TS compound treated samples. Black dots indicate the estimated $EC_{50}$ values and their 95 % confidence intervals (blue bars). The black vertical line indicate the Null hypothesis that the compared $EC_{50}$ values differ from each other. The corresponding p-values for each calculation are displayed on the right side. Further details on modelling and $EC_{50}$ derivation are given in the statistical supplementary statS1

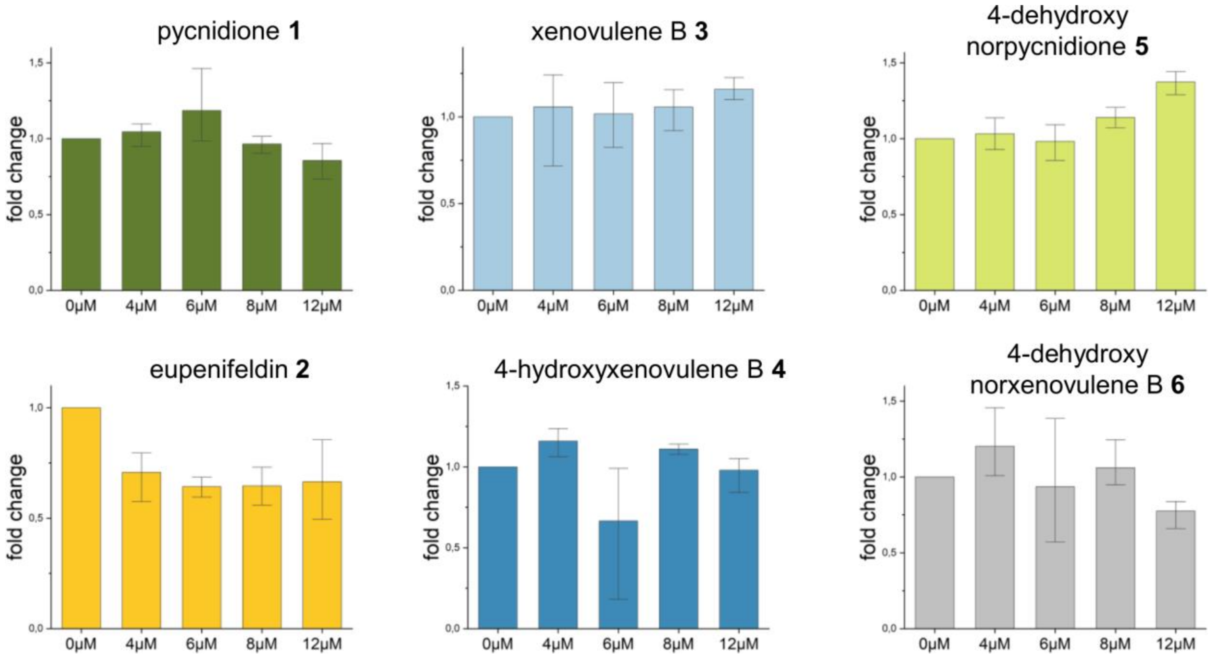


**Figure S 2.** Assessment of basic erythropoietin concentrations after administration of 4 µM, 6 µM, 8 µM and 12 µM of the tropolone sesquiterpenoids (TS) pycnidione **1**, eupenifeldin **2**, xenovulene B **3**, 4-hydroxyxenovulene B **4** and 4-dehydroxy norpycnidione **5** and the tropolone-moiety lacking compound 4-hydroxy norxenovulene B **6** for 24h (N=1 independent experiment). Initial tests on EPO concentrations were conducted to assess optimal compound concentration showing highest fold induction for pycnidione **1** at 6 µM and for 4 – dehydroxy norpycnidione **5** at 12 µM. Protein concentrations were obtained via ELISA and fold changes compared to the untreated control (0 µM) were calculated. Error bars indicate the standard deviation of the n = 3 technical replicates.

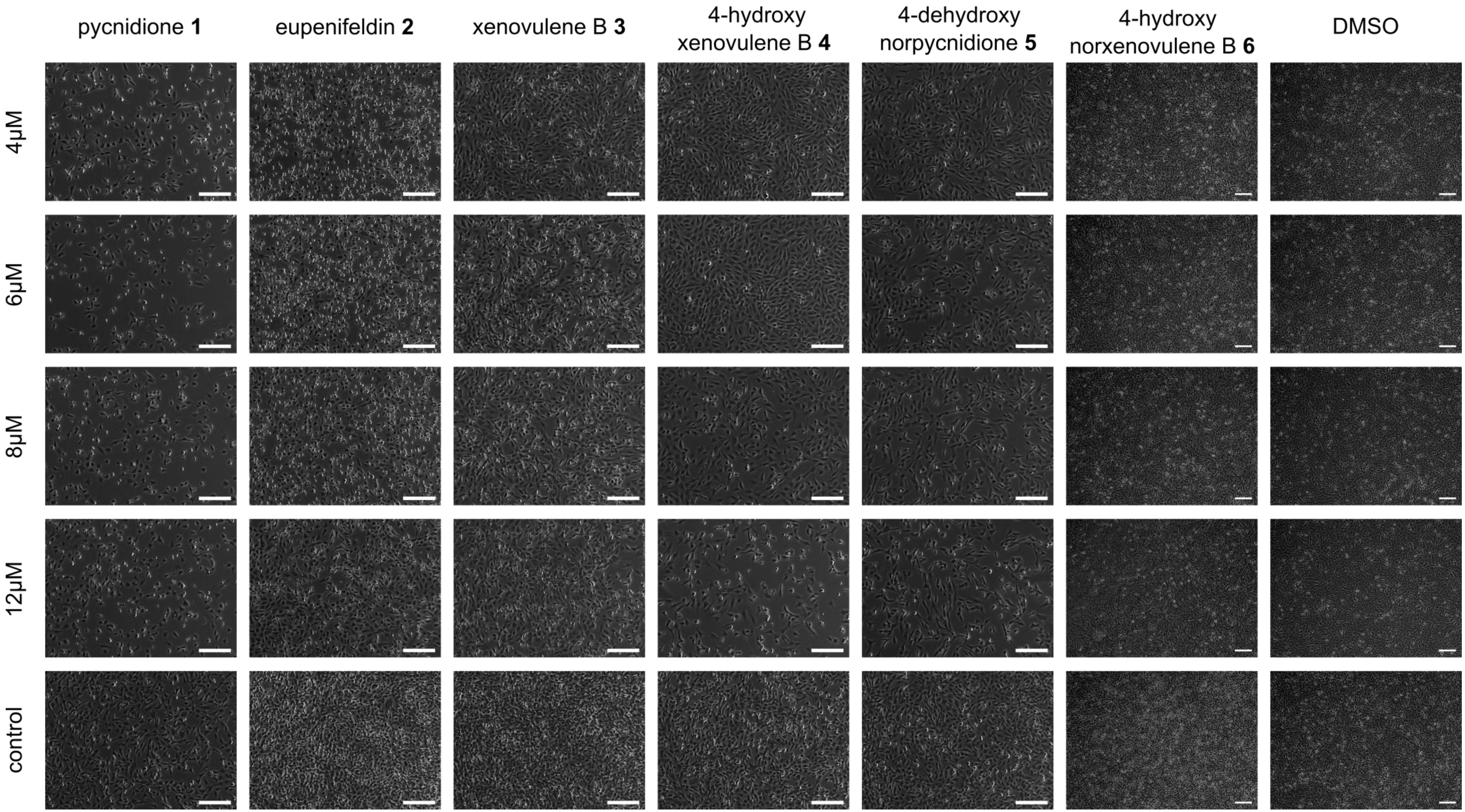


**Figure S 3. (Legend next page)**

**Figure S 3.** Phase contrast images of FAIK3-5 cell samples treated with 4 µM, 6 µM, 8 µM, and 12 µM of tropolone-sesquiterpenoids (TS) (pycnidione **1**, eupenifeldin **2**, xenovulene B **3**, 4-hydroxyxenovulene B **4** and 4-dehydroxy norpycnidione **5**), tropolone-lacking compound 4-hydoxy norxenovulene B **6**, the vehicle control dimethyl sulfoxide (DMSO) or cell culture medium (control) for 24 h. Scale bar = 100µM. TS treatment with compounds **1**, **2**, **3** and **5** induce a reduced cell density compared to the respective medium control, regardless of the compound concentration applied, whereas compound **4** shows a reduced cell confluence with higher concentrations of the compound. Treatment with tropolone-lacking compound **6** and DMSO does not influence cell densities in any applied concentration compared to the medium control.

**a**

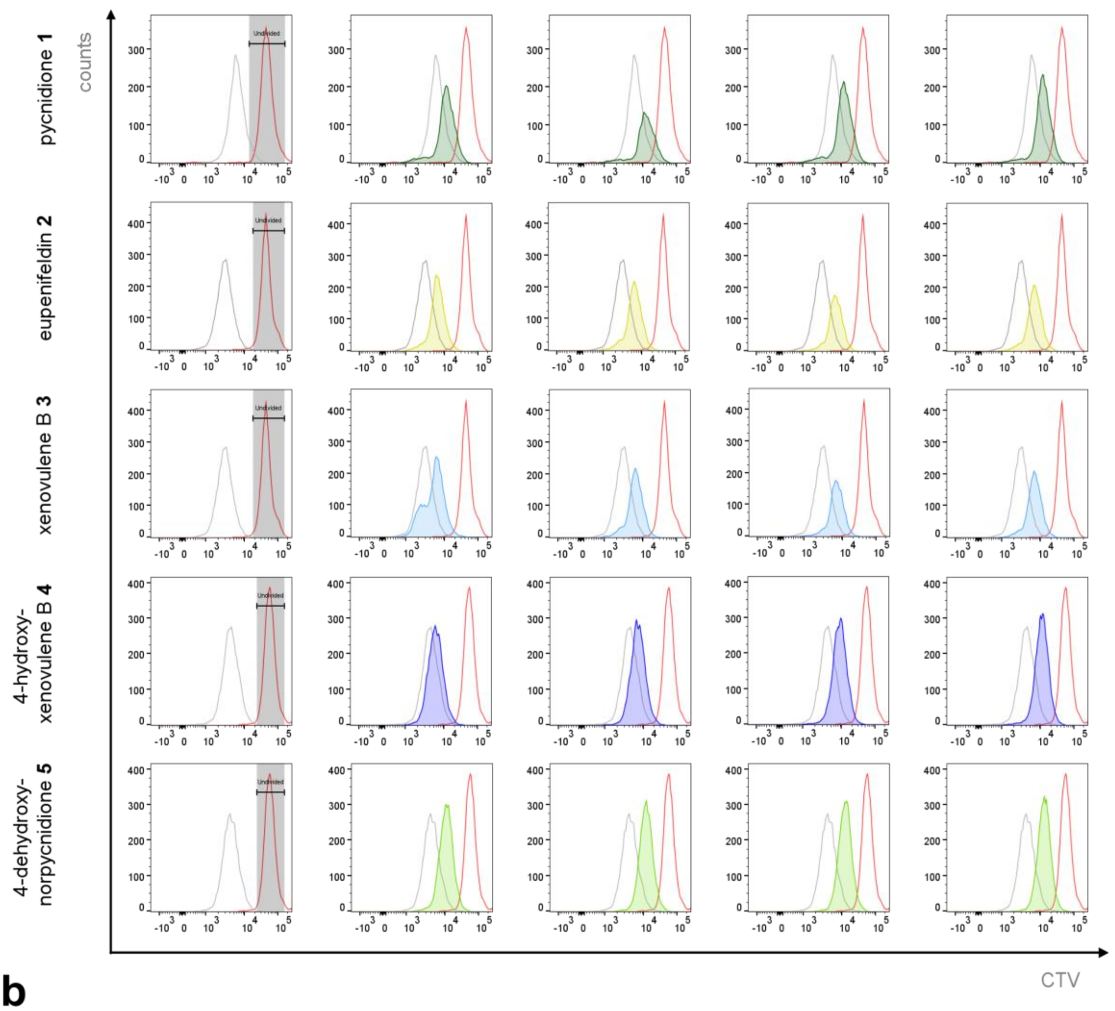


**b**

| | MFI (a.u.) | | | | | fold change | | | |
|---|---|---|---|---|---|---|---|---|---|
| | *0μM* | *4μM* | *6μM* | *8μM* | *12μM* | *4μM* | *6μM* | *8μM* | *12μM* |
| pycnidione **1** | 7124 | 12485 | 13732 | 12962 | 12985 | 1.75 | 1.93 | 1.82 | 1.82 |
| eupenifeldin **2** | 2946 | 6390 | 6689 | 6573 | 6747 | 2.17 | 2.27 | 2.23 | 2.29 |
| xenovulene B **3** | 3340 | 6937 | 6689 | 7281 | 7397 | 2.08 | 2.00 | 2.18 | 2.21 |
| 4-hydroxyxenovulene B **4** | 4108 | 5664 | 6792 | 8736 | 10747 | 1.38 | 1.65 | 2.13 | 2.62 |
| 4-dehydroxy norpycnidione **5** | 4091 | 10530 | 11534 | 12018 | 12496 | 2.57 | 2.82 | 2.94 | 3.05 |

**Figure S 4. (a)** Proliferation decrease of FAIK3-5 cells treated with 4 µM, 6 µM, 8 µM and 12 µM of tropolone-sesquiterpenoids (TS) (pycnidione **1**, eupenifeldin **2**, xenovulene B **3**, 4-hydroxyxenovulene B **4** and 4-dehydroxy norpycnidione **5**) for 24 h. Cell trace violet (CTV) stained cells were measured in a flow cytometer directly after staining to evaluate mean fluorescence intensities (MFI) of the undivided population (red) and 24 h after TS-compound treatment. MFI plots indicate a decrease of proliferation in samples treated with 4 µM – 12 µM of pycnidione **1** (dark green), eupenifeldin **2** (yellow), xenovulene B **3** (light blue) and 4-dehydroxy norpycnidione **5** (light green) assessed by consistently or increasingly higher MFI values compared to untreated controls (grey). In 4-hydroxyxenovulene B **4** (dark blue) treated samples an enhanced proliferation inhibition with increasing compound concentrations was observed, with the highest fluorescence intensities at a concentration of 12 µM **(b)** Mean fluorescence (MFI) values and fold changes of FAIK3-5 cells treated with tropolone sesquiterpenoids (TS) (pycnidione **1**, eupenifeldin **2**, xenovulene B **3**, 4-hydroxyxenovulene B **4** and 4-dehydroxy norpycnidione **5**) in a concentration rage of 0 µM, 4 µM, 6 µM, 8 µM and 12 µM. MFI values of cell trace violet stained samples indicate proliferative behaviour of cell samples, whereby higher intensities imply a downshift of cell divisions. Measurements were conducted 24 h after compound treatment and fold changes were calculated with respect to untreated control samples (0 µM).

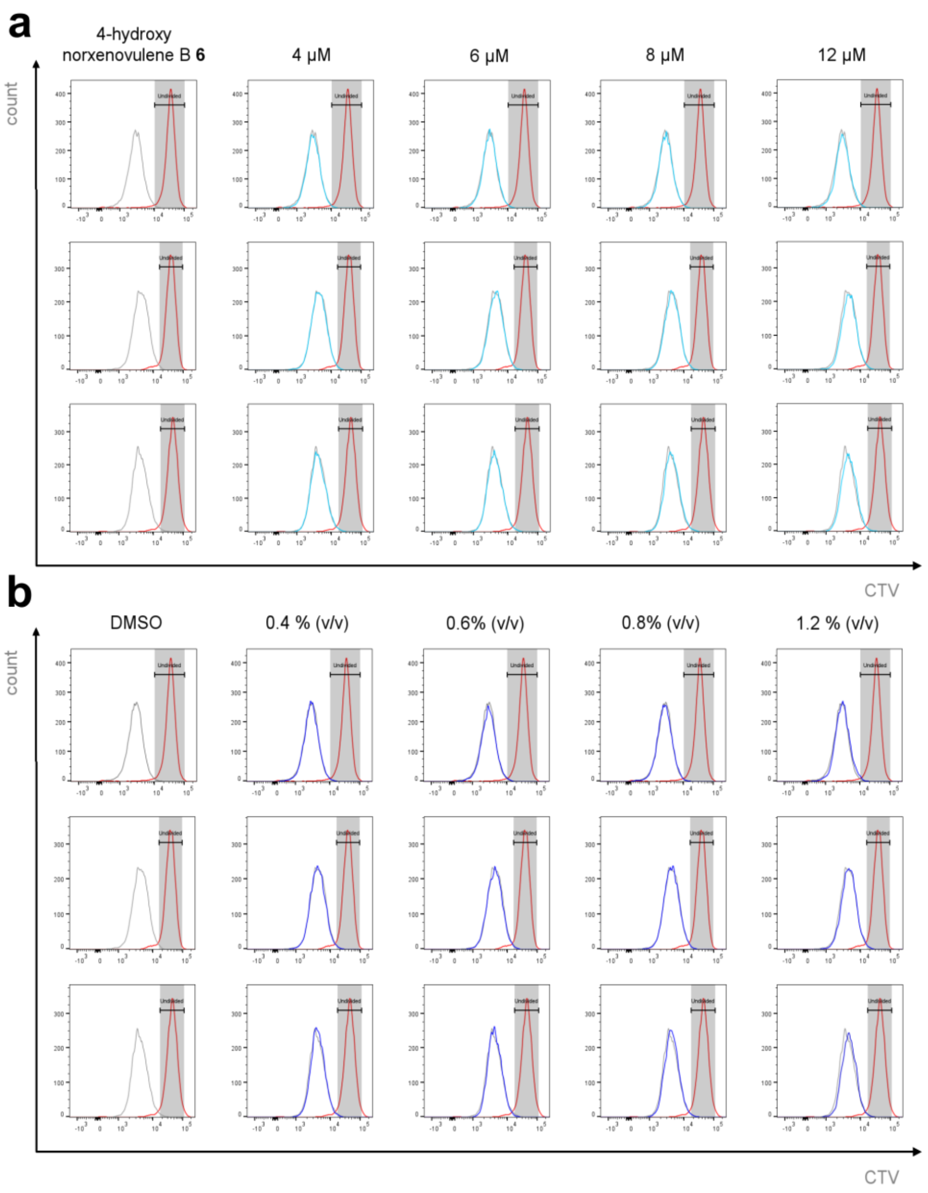


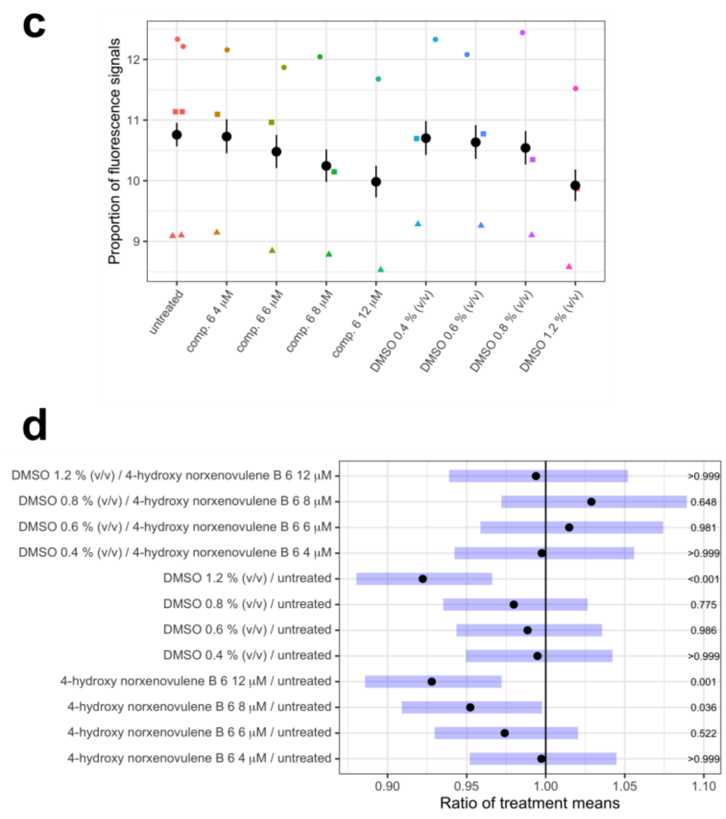


**e**

| | MFI (a.u.) | | | | | fold change | | | |
|---|---|---|---|---|---|---|---|---|---|
| | *0μM* | *4μM* | *6μM* | *8μM* | *12μM* | *4μM* | *6μM* | *8μM* | *12μM* |
| 4-hydroxy norxenovulene B **6** (1) | 2723 | 2739 | 2804 | 2761 | 2849 | 1.01 | 1.03 | 1.01 | 1.05 |
| 4-hydroxy norxenovulene B **6** (2) | 4066 | 4041 | 4185 | 4211 | 4334 | 0.99 | 1.03 | 1.04 | 1.07 |
| 4-hydroxy norxenovulene B **6** (3) | 3808 | 3823 | 3870 | 4180 | 4254 | 1.00 | 1.02 | 1.10 | 1.12 |
| | *0% (v/v)* | *0.4% (v/v)* | *0.6% (v/v)* | *0.8% (v/v)* | *1.2% (v/v)* | *0.4% (v/v)* | *0.6% (v/v)* | *0.8% (v/v)* | *1.2% (v/v)* |
| DMSO (1) | 2697 | 2697 | 2755 | 2676 | 2888 | 1.00 | 1.02 | 0.99 | 1.07 |
| DMSO (2) | 4066 | 3983 | 3999 | 4066 | 4308 | 0.98 | 0.98 | 1.00 | 1.06 |
| DMSO (3) | 3808 | 3967 | 3942 | 4099 | 4299 | 1.04 | 1.04 | 1.08 | 1.13 |

**Figure S 5 (Legend next page)**

**Figure S 5.** Proliferation decrease of FAIK3-5 cells treated with the tropolone-lacking compound 4-hydroxy norxenovulene B **6** and respective concentrations of the vehicle control DMSO for 24 h. **(a, b)** Cell trace violet (CTV) stained cells were measured in a flow cytometer directly after staining to evaluate mean fluorescence intensities (MFI) of the undivided population (red) and 24 h after incubation with 4 µM, 6 µM, 8 µM and 12 µM of 4-hydroxy norxenovulene B **6** (light blue), 0.4% (v/v), 0.6% (v/v), 0.8% (v/v) and 1.2% (v/v) of the DMSO vehicle control (dark blue) and untreated controls (grey). **(c)** For statistical analysis obtained MFI values were normalized by calculating the proportions of measured fluorescence signals on d1 and d3. The graphic shows the observed proportions of fluorescence signals ($y^1/y^3$) for each treatment and their corresponding least square means (black dots) together with 95% confidence intervals. **(d)** Pairwise mean comparisons between 4-hydroxy norxenovulene B **6**, DMSO and the untreated control. The black line indicates the null hypothesis that the ratio between the average fluorescence signals (black dots) is one. The blue horizontal bars indicate simultaneous 95 % confidence intervals for the mean ratios. If a confidence interval encompasses the one, the ratio does not significantly differ from one. The numbers on the right side of the graphic are the p-values of the corresponding test (see statistical supplementary statS6). **(e)** Mean fluorescence (MFI) values and fold changes of FAIK3-5 cells treated with tropolone moiety lacking compound 4-hydroxy norxenovulene B **6** (N=3) and DMSO vehicle controls (N=3) in a concentration rage of 0 µM, 4 µM, 6 µM, 8 µM and 12 µM or respective volume percentages (0 % (v/v), 0.4 % (v/V), 0.6 % (v/v), 0.8 % (v/v) and 1.2 % (v/v). MFI values of cell trace violet stained samples indicate proliferative behavior of cell samples, whereby higher intensities imply a downshift of cell divisions. Measurements were conducted 24 h after compound treatment and fold changes were calculated with respect to untreated control samples (0 µM and 0 % (v/v)).

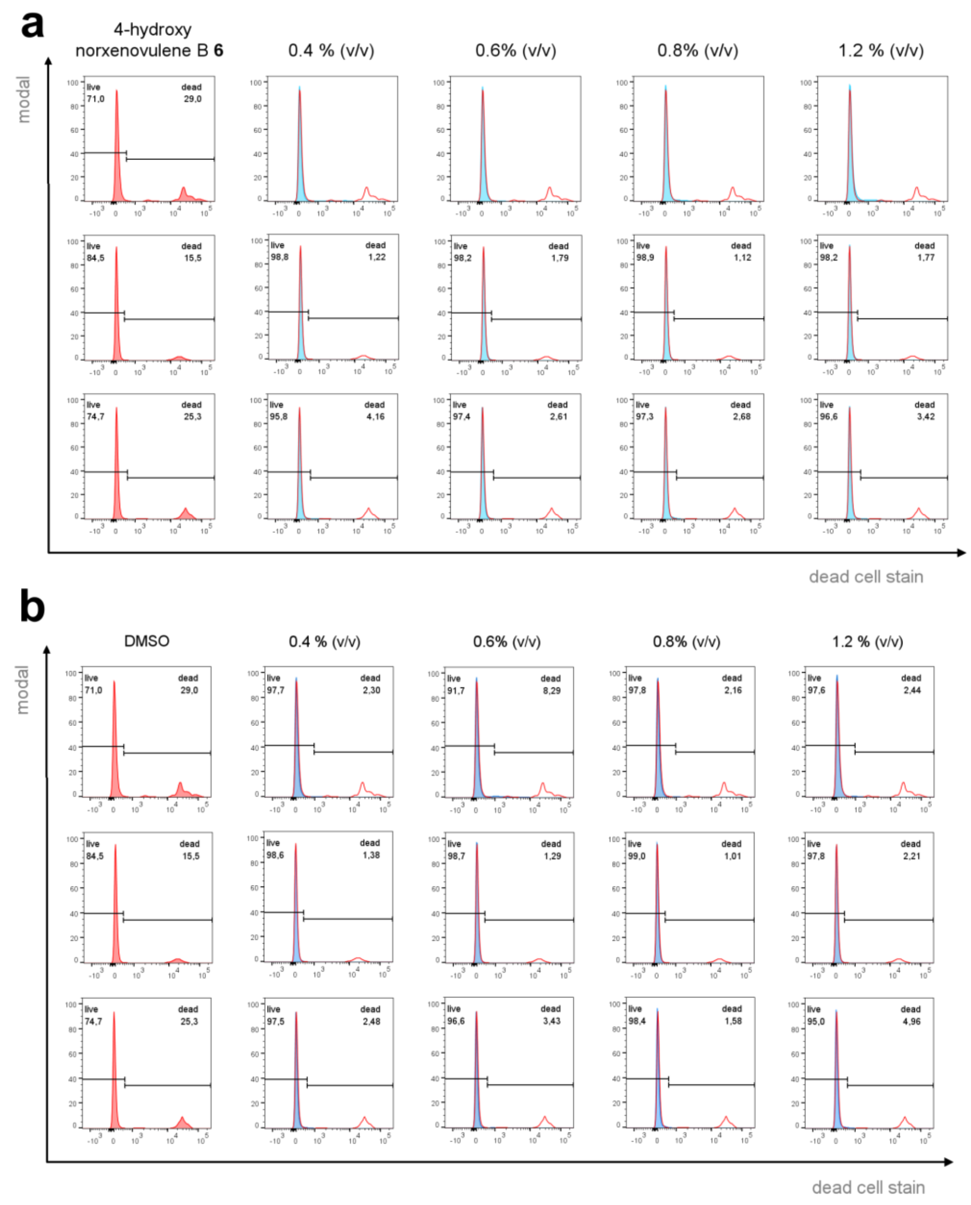


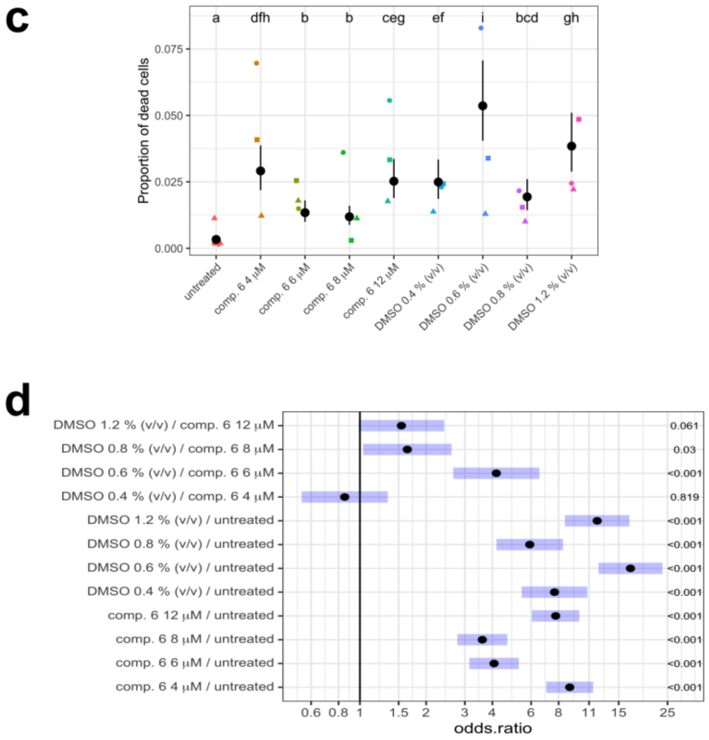


**e**

| | cell viability [%] | | | | |
|---|---|---|---|---|---|
| | *0µM* | *4µM* | *6µM* | *8µM* | *12µM* |
| pycnidione **1** | 99.3 | 68.1 | 72.2 | 77.9 | 76.5 |
| eupenifeldin **2** | 99.8 | 72.1 | 69.4 | 55.8 | 66.8 |
| xenovulene B **3** | 99.9 | 92.9 | 92.9 | 89.8 | 89.8 |
| 4-hydroxyxenovulene B **4** | 99.9 | 96.3 | 96.1 | 95.6 | 94.3 |
| 4-dehydroxy norpycnidione **5** | 99.9 | 94.7 | 92.7 | 94.1 | 94.7 |
| 4-hydroxy norxenovulene B **6** (1) | 99.8 | 93.0 | 98.5 | 96.4 | 94.4 |
| 4-hydroxy norxenovulene B **6** (2) | 98.9 | 98.8 | 98.2 | 98.9 | 98.2 |
| 4-hydroxy norxenovulene B **6** (3) | 99.7 | 95.8 | 97.4 | 97.3 | 96.6 |
| | *0% (v/v)* | *0.4% (v/v)* | *0.6% (v/v)* | *0.8% (v/v)* | *1.2% (v/v)* |
| DMSO (1) | 99.8 | 97.7 | 91.7 | 97.8 | 97.6 |
| DMSO (2) | 99.8 | 98.6 | 98.7 | 99.0 | 97.8 |
| DMSO (3) | 99.7 | 97.5 | 96.6 | 98.4 | 95.0 |

**Figure S 6. (Legend next page)**

**Figure S 6.** Assessment of cell viability of FAIK3-5 cells treated with **(a)** the tropolone moiety lacking compound 4-hydroxy norxenovulene B **6** and **(b)** DMSO vehicle control in a concentration rage of 0 µM, 4 µM, 6 µM, 8 µM and 12 µM or respective volume percentages (0 % (v/v), 0.4 % (v/v), 0.6 % (v/v), 0.8 % (v/v) and 1.2 % (v/v). Percentages of dead cells were obtained via SYTOX™ AADvanced™ dead cell staining and flow cytometric analysis of FAIK3-5 cells 24h after treatment with the compound and solvent controls. To set the threshold for dead cells' fluorescence intensities a living-dead control comprising a population of heat-killed FAIK3-5 cells was utilized and all samples were visualized in relation to the control (red). **(c)** Graphical overview of the data distribution of the proportion of dead cells in different experimental replicates (N=3) for each condition and the untreated control show model based means (black dots) and their 95% confidence intervals. Treatments marked with the same letter do not differ statistically significantly from each other ($\alpha = 0.05$). **(d)** Multiple comparison analysis was conducted based on odds-ratios of treatment means (black dots) that indicate the average chance to obtain dead cells in treatment conditions and the untreated control. Blue bars indicate simultaneous 95% confidence intervals of calculated ratios and p-values for each analysis are displayed on the ride side (see statistical supplementary statS8). (e) Cell viability of FAIK3-5 cells treated with tropolone sesquiterpenoids (TS) (pycnidione 1, eupenifeldin 2, xenovulene B 3, 4-hydroxyxenovulene B 4 and 4-dehydroxy norpycnidione 5), tropolone moiety lacking compound 4-hydroxy norxenovulene B 6 (N=3) and DMSO vehicle controls (N=3) in a concentration rage of 0 µM, 4 µM, 6 µM, 8 µM and 12 µM or respective volume percentages (0 % (v/v), 0.4 % (v/V), 0.6 % (v/v), 0.8 % (v/v) and 1.2 % (v/v). Percentages of cell viability were obtained via SYTOX™ AADvanced™ staining and subsequent flow cytometric analysis.

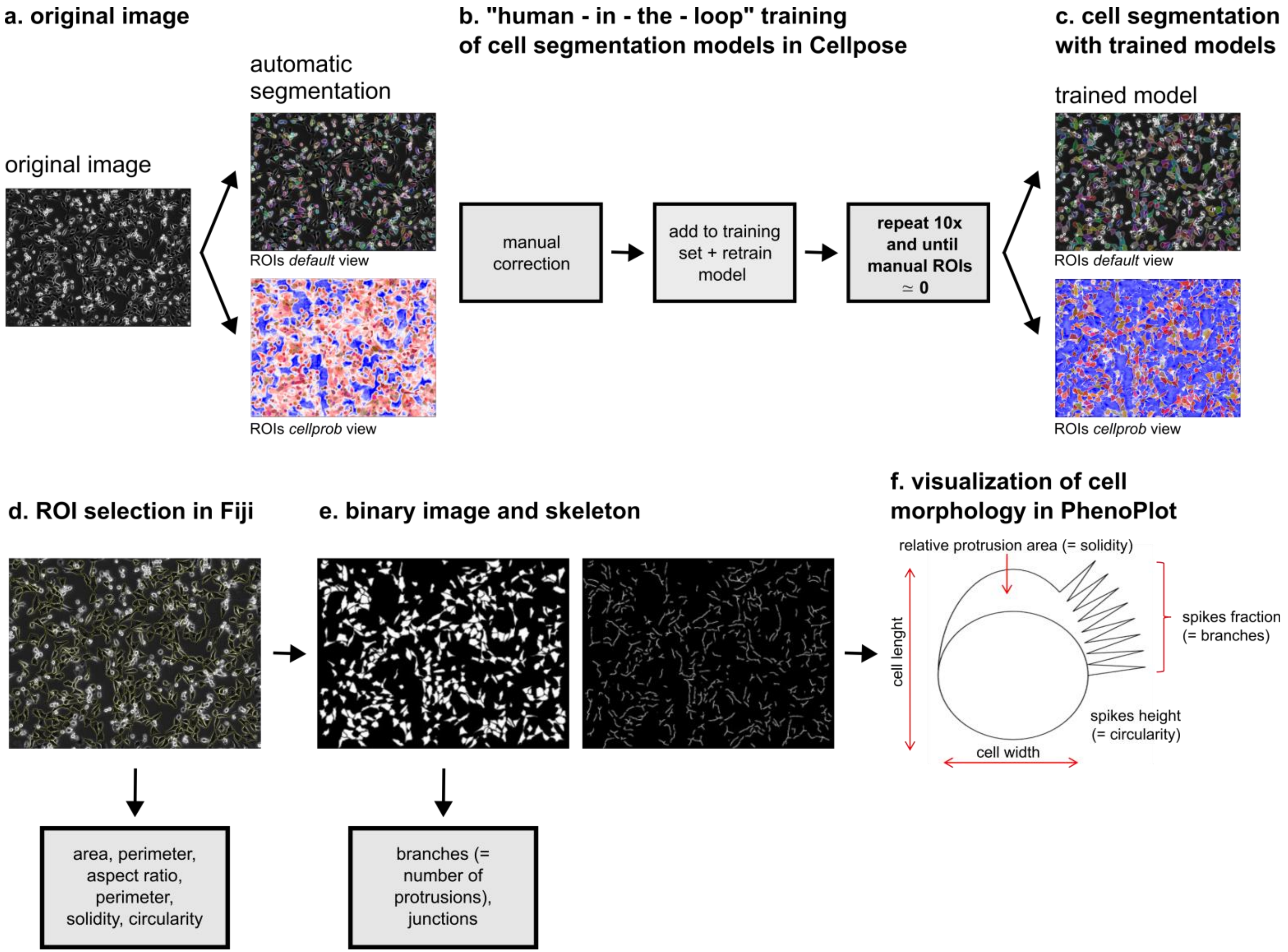


**Figure S 7. (Legend next page)**

**Figure S 7.** Workflow for analysis of morphometric parameters and ramification patterns. **(a)** For cell segmentation brightfield images were analyzed using the generalist deep-learning based algorithm *Cellpose [1]*. **(b)** Two cell segmentation models were trained to capture specific features of cell morphology accurately due to the high morphological variance between cells treated with bistropolones (**1** and **2**) (model 1) and those treated with monotropolons (**3**, **4** and **5**), the tropolone lacking compound **6** and untreated controls (model 2). This variance is caused by the increased occurrence of the telocyte-like subtype with nanotubes connecting the cells in the bistropolone samples. To operate *Cellpose* an Anaconda distribution (*Anaconda Software Distribution*. Vers. 2-2.4.0.) of Python (Version 3.8) [2] was used and a new Python environment for *Cellpose* was created in PowerShell Prompt (Version 0.0.1). (Further information for the installation of *Cellpose* can be found on https://github.com/MouseLand/cellpose). For the manual "human-in-the-loop" training of each model, we followed the same operational sequence on a minimum of 10 images. We continued until no further manual correction of the detected cells, referred to here as regions of interest (ROIs), was required. Firstly, we calibrated the cell diameter and then ran the default algorithm "*cyto2*" over the first training image. We used a flow threshold of 0.4, a cellprob threshold of 0.0, a stitch threshold of 0.0, and auto-adjusted the image saturation. ROIs were adjusted manually by removing incorrectly drawn ROIs and adding undetected ROIs to the image. To verify differences in cell morphology based on the cell's adjustments to TS compounds and controls, only living, adherent cells were included in the analysis and the models were trained to exclude oversaturated, floating cells. **(c)** For the final analysis the trained models were run on images of FAIK3-5 cells treated with the investigated compounds (**1**–**6**) and the untreated control of three independent experiments (N=3), without further manual correction of the detected ROIs. To ensure accurate measurements, we only included complete cells in our analysis. Therefore, we excluded ROIs that detected cells cut off by the image edges or overlapping regions. **(d)** Detected ROIs were transferred to Fiji (ImageJ Version 1.53c bundled with Java 1.8.0_172) [3], an overlay with the original image was created to check correct ROI selection and measurement of cell area, shape descriptors (aspect ratio, circularity, roundness and solidity), perimeter and was conducted. **(f)** For the analysis of ramification patterns, selected ROIs of separated cellular shapes were used to create a binary image, on which the "Skeletonize 2D/3D" and "Analyze Skeleton (2D/3D)" [4] plugins were run for visualization and calculation of e.g. branch numbers (via connection of end-points, end-points and junctions, or junctions and junctions) and number of branch junctions. The obtained values for geometrical features, shape descriptors and branch numbers were normalized to the highest obtained mean value of each analysed parameter for subsequent visualization in PhenoPlot [5].

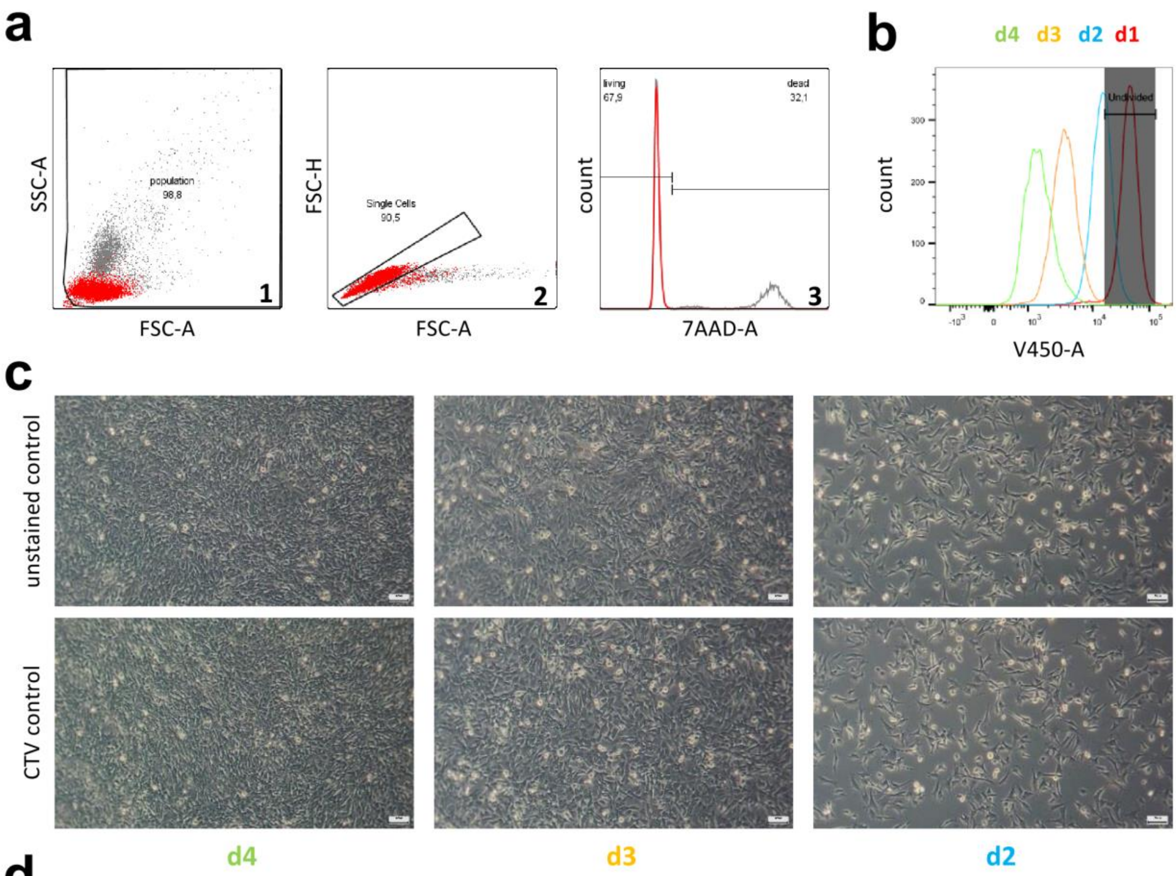


**d**

| | mean fluorescence intensity | counted living cells | viability [%] |
|---|---|---|---|
| **d1** | 46147 | 8393 | 91.7 |
| **d2** | 15444 | 9535 | 97.0 |
| **d3** | 3936 | 9432 | 97.4 |
| **d4** | 1693 | 9482 | 98.3 |

**Figure S 8.** Gating strategy for cytotoxicity assessment and proliferation analysis and preliminary tests on FAIK3-5 cells to assess optimal experimental conditions for subsequent proliferation analysis. **(a)** Backgating plots of the living cell population show the strategy for living/dead discrimination in a living/dead control sample comprising 50 % of heat-killed FAIK3-5 cells. After out-gating of (1) cell debris and (2) doublets, the (3) living and dead population is visualized in a histogram showing fluorescence intensities of Sytox AADvanced stained cells (7AAD-A channel). Thereafter, the threshold for dead cells' fluorescence intensities was set in the living/dead control and assigned to the remaining samples. **(b)** For proliferation analysis the mean fluorescence intensity (MFI) of cell trace violet (CTV, measured in V450-A channel) of samples on d1 was set as undivided population in the living cell population of the analysed samples. To assess the optimal staining period for proliferation tests, CTV intensities and cell viability of stained FAIK3-5 cells were analysed on three subsequent days. **(c, d)** Proliferative behavior of FAIK3-5 cells was evaluated by normalizing the measured fluorescence intensities at day 2 - 4 to their respective MFIs of the undivided population at d1 of the experiment. As the cells continuously proliferate (**c**), the intensity of CTV decreases with each cell division, while cell viability remains over 90 % (d). For CTV staining and living/dead analysis 10 000 events per sample were counted.
