## Supplementary material for "Evaluating the bioactivities of natural and synthetic fungal tropolone sesquiterpenoids in various cell lines": Statistical Supplementary Information

Dr. M. Menssen

[2] Institute of Cell Biology and Biophysics, Department of Biostatistics, Leibniz University Hannover, 30419 Hannover, Germany

Dr. C. Schotte, Prof. Dr. R.J.Cox

[3] Institute of Organic Chemistry and BMWZ, Leibniz University Hannover, 30167 Hannover, Germany

* Prof. Dr. Cornelia Lee-Thedieck, Leibniz University Hannover, Institute of Cell Biology and Biophysics, Herrenhäuserstr 2, 30419 Hannover. Phone: +49 511 762 14 972; Fax: +49 511 762 2606;

Statistical supplementary statS6: Proliferation for 4-hydroxy norxenovulene B 6 and DMSO. 25

### Statistical supplementary statS1: Estimations of EC50

Bergmann et al. 2024

#### Data

```
# First rows of the data
head(dat)
```

```
##      line     compound repetition  conc  mean_int
## 1 FAIK3_5 pycnidione_1          1 0e+00 17218.000
## 2 FAIK3_5 pycnidione_1          1 1e-03 19811.667
## 3 FAIK3_5 pycnidione_1          1 1e-02 20228.667
## 4 FAIK3_5 pycnidione_1          1 1e-01  8383.333
## 5 FAIK3_5 pycnidione_1          1 1e+00  7260.333
## 6 FAIK3_5 pycnidione_1          1 1e+01  4559.333
```

- line: Indecator for the cell lines
- compound: Indecator for the six compounds and DMSO without a compound as control
- repetition: Indecator wich experiment belongs to the specific repetition (three repetitions in all)
- mean_int: Mean fluorescence intensity, average over 12 observations (four per each of three wells)
- conc: Concentration

#### Hierarchical non linear modelling

For each cell line, the EC50s of the compounds were estimated based on hierarchical nonlinear regression models. All fitted models were based on the four-parameter log-logistic function with parameters

- b: Steepness of the curve at EC50
- c: Lower asymptote
- e: EC50
- d: Upper asymptote

In each of the fitted models (one for each cell line), the lower asymptote was estimated from all observations, regardless to which compound they belong. For MDA-MD 231 cells, also the steepness parameter was held fixed for all compounds (due to convergance problems), but e, d were allowed to differ between the compounds. For the other three cell lines, the parameters b, e, d were allowed to differ between the compounds. For all four models, the parameters d and e were modelled as random effects.

EC50s were derived from the fitted models and for each cell line, the estimated EC50s for the compounds were tested against the EC50 of DMSO in a Dunnett type test procedure (p-value adjustment following Hothoern et al. 2008).

##### FAIK3-5

```
# Subset for FAIK3-5 cells
dat_faik <- filter(dat, line=="FAIK3_5", compound != "DMSO_without_cells")
%>%
```

```
        mutate(comrep=interaction(compound, repetition)) %>%
        select(mean_int, conc, comrep, compound) %>%
        droplevels() %>%
        data.frame()

# Hierarchical nonlinear model
fit_faik <- medrm(mean_int~conc,
                  data=dat_faik,
                  fct=LL.4(),
                  curveid=b + e + d ~ compound,
                  random= d + e ~ 1|comrep)

# Estimated EC50s, their standard errors and 95% confidence intervals (lowe
r, upper)
as.data.frame(ED(fit_faik, 50, interval="delta", display = FALSE))

##                                      Estimate Std. Error       Lower
Upper
## e:4_dehydroxy_norpycnidione_5:50  1.4648800 0.44754053  0.58174587  2.34
80142
## e:4_hydrox_xenovulene_B_4:50      6.4576130 1.62448228  3.25201326  9.66
32127
## e:4_hydroxy_norxenovulene_B_6:50 22.7257179 2.62369080 17.54837461 27.90
30612
## e:DMSO_with_cells:50             22.9778857 2.48126862 18.08158488 27.87
41866
## e:eupenifeldin_2:50               0.1278044 0.05808026  0.01319430  0.24
24145
## e:pycnidione_1:50                 0.1973568 0.09219915  0.01541975  0.37
92939
## e:xenovulene_B_3:50               2.8011319 0.84234664  1.13892467  4.46
33391

# Dunnett test of the EC50s

ED_multcomp <- ED(fit_faik,
                  50,
                  interval="delta",
                  multcomp = TRUE,
                  display=FALSE)

nvec <- 1:7
names(nvec) <- names(ED_multcomp$EDmultcomp$coef)

summary(glht(ED_multcomp[["EDmultcomp"]],
                                linfct=contrMat(nvec,
                                                "Dunnett",
                                                base=4)))

##    lhs1 rhs    estimate        lwr        upr
## 1   5-7   0 -21.5130057 -27.389563 -15.636449
## 2   4-7   0 -16.5202728 -23.428357  -9.612189
## 3   6-7   0  -0.2521678  -8.532694   8.028358
## 4   2-7   0 -22.8500813 -28.675879 -17.024283
## 5   1-7   0 -22.7805289 -28.605027 -16.956030
## 6   3-7   0 -20.1767539 -26.245270 -14.108238
```

Jurkat

```
# Subset for Jurkat cells
dat_jurkat <- filter(dat, line=="Jurkat", compound != "DMSO_without_cells")
 %>%
        mutate(comrep=interaction(compound, repetition)) %>%
        select(mean_int, conc, comrep, compound) %>%
        droplevels() %>%
        data.frame()

# Hierarchical nonlinear model
fit_jurkat <- medrm(mean_int~conc,
                   data=dat_jurkat,
                   fct=LL.4(),
                   curveid=b + e + d ~ compound,
                   random= d + e ~ 1|comrep)

# Estimated EC50s, their standard errors and 95% confidence intervals (lowe
r, upper)
as.data.frame(ED(fit_jurkat, 50, interval="delta", display = FALSE))

##                                    Estimate Std. Error       Lower
Upper
## e:4_dehydroxy_norpycnidione_5:50  3.2130030 0.62143604  1.98671986  4.43
92861
## e:4_hydrox_xenovulene_B_4:50     10.5252475 0.87274227  8.80306043 12.24
74346
## e:4_hydroxy_norxenovulene_B_6:50 21.5883406 1.45271227 18.72169554 24.45
49856
## e:DMSO_with_cells:50             17.1239168 1.67252971 13.82350480 20.42
43287
## e:eupenifeldin_2:50               0.0689933 0.03927945 -0.00851705  0.14
65037
## e:pycnidione_1:50                 0.1957781 0.06811034  0.06137559  0.33
01806
## e:xenovulene_B_3:50               2.6283387 0.63275187  1.37972599  3.87
69514

# Dunnett test of the EC50s

ED_multcomp <- ED(fit_jurkat,
                  50,
                  interval="delta",
                  multcomp = TRUE,
                  display=FALSE)

nvec <- 1:7
names(nvec) <- names(ED_multcomp$EDmultcomp$coef)

summary(glht(ED_multcomp[["EDmultcomp"]],
                                linfct=contrMat(nvec,
                                                "Dunnett",
                                                base=4)))

##    lhs1 rhs    estimate         lwr        upr
## 1   5-7   0 -13.910914 -18.0571301  -9.764697
```

```
## 2  4-7   0  -6.598669 -10.9864917  -2.210847
## 3  6-7   0   4.464424  -0.6890397   9.617887
## 4  2-7   0 -17.054923 -20.9775078 -13.132339
## 5  1-7   0 -16.928139 -20.8502719 -13.006005
## 6  3-7   0 -14.495578 -18.6441720 -10.346984
```

##### MDA-MD 231

```
# Subset for MDA-MD 231 cells
dat_mda_md <- filter(dat, line=="MDA_MD_231", compound != "DMSO_without_cel
ls") %>%
        mutate(comrep=interaction(compound, repetition)) %>%
        select(mean_int, conc, comrep, compound) %>%
        droplevels() %>%
        data.frame()

# Hierarchical nonlinear model
fit_mda_md <- medrm(mean_int~conc,
                     data=dat_mda_md,
                     fct=LL.4(),
                     curveid= e + d ~ compound,
                     random= d + e ~ 1|comrep)

## Warning in nlme.formula(mean_int ~ meLL.4(conc, b, c, d, e), fixed = lis
t(b ~ :
## Iteration 1, LME step: nlminb() did not converge (code = 1). Do increase
## 'msMaxIter'!

# Estimated EC50s, their standard errors and 95% confidence intervals (lowe
r, upper)
as.data.frame(ED(fit_mda_md, 50, interval="delta", display = FALSE))

##                                    Estimate Std. Error    Lower    Upper
## e:4_dehydroxy_norpycnidione_5:50 16.28259   2.804129 10.74918 21.81599
## e:4_hydrox_xenovulene_B_4:50     18.41708   3.328798 11.84835 24.98582
## e:4_hydroxy_norxenovulene_B_6:50 17.78042   2.965391 11.92879 23.63204
## e:DMSO_with_cells:50             20.29586   3.337690 13.70958 26.88214
## e:eupenifeldin_2:50              16.44847   3.144468 10.24348 22.65347
## e:pycnidione_1:50                16.64705   3.143708 10.44355 22.85055
## e:xenovulene_B_3:50              18.58067   3.309810 12.04940 25.11194

# Dunnett test of the EC50s

ED_multcomp <- ED(fit_mda_md,
                  50,
                  interval="delta",
                  multcomp = TRUE,
                  display=FALSE)

nvec <- 1:7
names(nvec) <- names(ED_multcomp$EDmultcomp$coef)

summary(glht(ED_multcomp[["EDmultcomp"]],
                              linfct=contrMat(nvec,
                                              "Dunnett",
                                              base=4)))
```

```
##   lhs1 rhs  estimate       lwr      upr
## 1  5-7   0 -4.013272 -14.37244 6.345894
## 2  4-7   0 -1.878778 -13.00179 9.244231
## 3  6-7   0 -2.515441 -13.11113 8.080248
## 4  2-7   0 -3.847386 -14.67960 6.984828
## 5  1-7   0 -3.648808 -14.48749 7.189874
## 6  3-7   0 -1.715189 -12.81746 9.387079
```

PC-3

```
# Subset for Jurkat cells
dat_pc3 <- filter(dat, line=="PC_3", compound != "DMSO_without_cells") %>%
        mutate(comrep=interaction(compound, repetition)) %>%
        select(mean_int, conc, comrep, compound) %>%
        droplevels() %>%
        data.frame()

# Hierarchical nonlinear model
fit_pc3 <- medrm(mean_int~conc,
                 data=dat_pc3,
                 fct=LL.4(),
                 curveid=b + e + d ~ compound,
                 random= d + e ~ 1|comrep)

# Estimated EC50s, their standard errors and 95% confidence intervals (lowe
r, upper)
as.data.frame(ED(fit_pc3, 50, interval="delta", display = FALSE))

##                                   Estimate Std. Error      Lower     Uppe
r
## e:4_dehydroxy_norpycnidione_5:50  5.041838   1.231958  2.610808  7.47286
8
## e:4_hydrox_xenovulene_B_4:50     14.387172   2.112586 10.218394 18.55595
0
## e:4_hydroxy_norxenovulene_B_6:50 15.182357   1.716479 11.795220 18.56949
4
## e:DMSO_with_cells:50             14.722614   1.792643 11.185182 18.26004
6
## e:eupenifeldin_2:50               7.605650   1.662656  4.324722 10.88657
8
## e:pycnidione_1:50                 6.257120   1.469021  3.358292  9.15594
8
## e:xenovulene_B_3:50               8.610989   1.717159  5.222510 11.99946
7

# Dunnett test of the EC50s

ED_multcomp <- ED(fit_pc3,
                  50,
                  interval="delta",
                  multcomp = TRUE,
                  display=FALSE)

nvec <- 1:7
names(nvec) <- names(ED_multcomp$EDmultcomp$coef)
```

```
summary(glht(ED_multcomp[["EDmultcomp"]],
                                 linfct=contrMat(nvec,
                                                 "Dunnett",
                                                 base=4)))
```

```
##   lhs1 rhs   estimate        lwr        upr
## 1  5-7   0 -9.6807758 -15.075650 -4.2859020
## 2  4-7   0 -0.3354424  -7.199931  6.5290457
## 3  6-7   0  0.4597432  -5.661179  6.5806654
## 4  2-7   0 -7.1169640 -13.164776 -1.0691518
## 5  1-7   0 -8.4654941 -14.198617 -2.7323707
## 6  3-7   0 -6.1116252 -12.265527  0.0422762
```

### Statistical supplementary statS2: Proteins

Bergmann et al. 2024

```r
# Load the packages used for evaluation
library(tidyverse)
library(lme4)
library(lmerTest)
library(tidyverse)
library(emmeans)
```

#### Overview about the data

Description of the variables

- Rep: Replication
- Plate: Plate in each replication
- Well: Well in each plate in each replication
- Compund: Chemical compounds (treatment variable)
- Protein: Protein content in pg/ml

```r
# First six rows of the data set
head(prote)

##   Rep Plate Well       Compound  Protein
## 1   1     1    1   pycnidione 1 5.819760
## 2   1     1    1   pycnidione 1 4.158405
## 3   1     1    1   pycnidione 1 4.302978
## 4   1     1    2 xenovulene B 3 7.161895
## 5   1     1    2 xenovulene B 3 6.590697
## 6   1     1    2 xenovulene B 3 7.172868

# 69 observations at all (rows)
str(prote)

## 'data.frame':    69 obs. of  5 variables:
##  $ Rep     : Factor w/ 3 levels "1","2","3": 1 1 1 1 1 1 1 1 1 1 ...
##  $ Plate   : Factor w/ 2 levels "1","2": 1 1 1 1 1 1 1 1 1 1 ...
##  $ Well    : Factor w/ 6 levels "1","2","3","4",..: 1 1 1 2 2 2 3 3 3 4
...
##  $ Compound: Factor w/ 8 levels "DMSO 0.6 % (v/v)",..: 3 3 3 5 5 5 4 4 4
 7 ...
##  $ Protein : num  5.82 4.16 4.3 7.16 6.59 ...
```

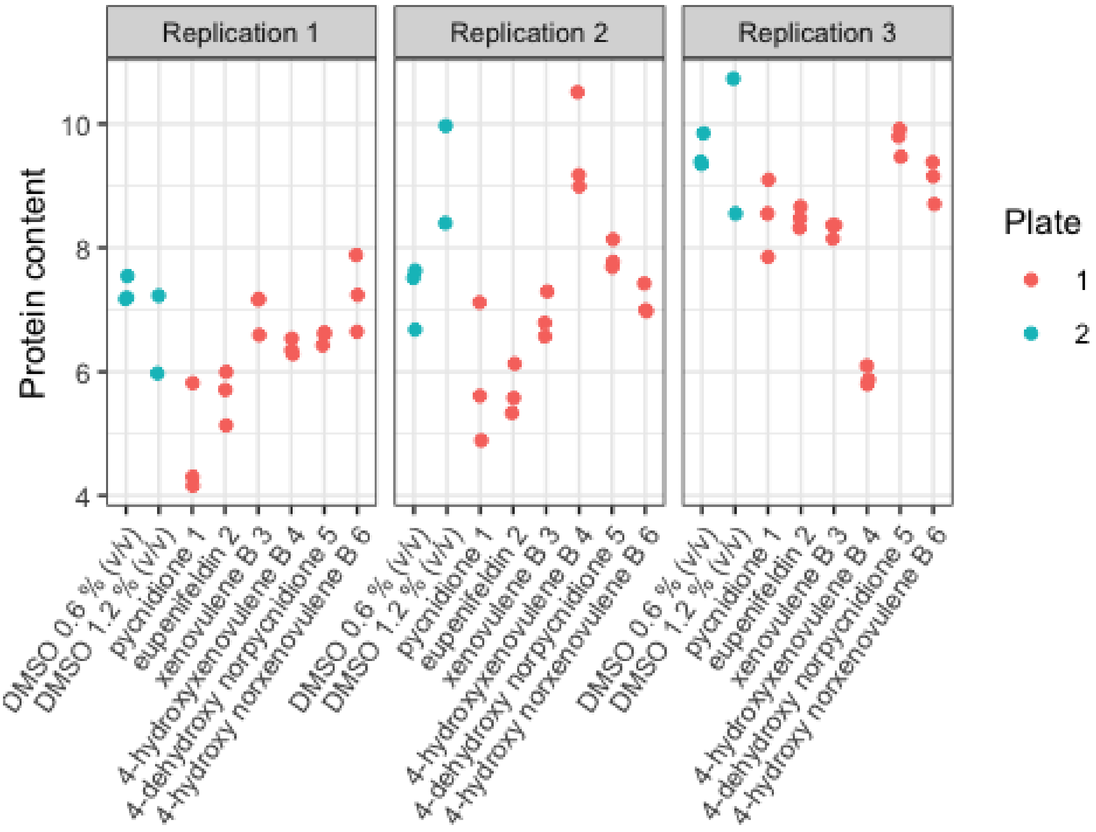

#### Statistical model

Hierarchical experimental design, since the plates are nested in the replications and the wells are nested in each plate.

- Linear mixed model
- Protein as response variable
- Compounds as fixed effects
- Rep, Plates and Wells as random effects

```
fit <- lmer(Protein ~ Compound + (1|Rep) + (1|Rep:Plate) + (1|Rep:Plate:Wel
l),
            data=prote)
```

##### Estimated variance components

```
print(VarCorr(fit),comp=c("Variance"),digits=2)

##  Groups          Name        Variance
##  Rep:Plate:Well (Intercept) 1.2e+00
##  Rep:Plate      (Intercept) 1.5e-09
##  Rep            (Intercept) 9.7e-01
##  Residual                   4.0e-01
```

- Most of the total variance can be explained by the wells.
- The plates do not explain any variance

```
anova(fit,
      ddf="Kenward-Roger")
```

```
## Type III Analysis of Variance Table with Kenward-Roger's method
##          Sum Sq Mean Sq NumDF  DenDF F value Pr(>F)
## Compound 3.3223 0.47462     7 10.514  1.1038 0.4261
```

- The p-value of 0.4261 is bigger than $\alpha = 0.05$
- No different mean protein contents between the compounds

### Statistical supplementary statS3: Proteins after 5-Azacytidine treatment

Bergmann et al. 2024

```
# Load the packages used for evaluation
library(tidyverse)
library(lme4)
library(lmerTest)
library(tidyverse)
library(emmeans)
library(multcomp)
```

#### Overview about the data

Description of the variables

- Rep: Replication
- Plate: Plate in each replication
- Well: Well in each plate in each replication
- Compund: Chemical compounds (treatment variable)
- Protein: Protein content in pg/ml

```
# First six rows of the data set
head(aza)

##   Rep Plate Well       Compound  Protein
## 1   1     1    1   pycnidione 1 22.10247
## 2   1     1    1   pycnidione 1 19.06987
## 3   1     1    1   pycnidione 1 18.17886
## 4   1     1    2 xenovulene B 3 17.96557
## 5   1     1    2 xenovulene B 3 14.10570
## 6   1     1    2 xenovulene B 3 17.36827

# 72 observations at all (rows)
str(aza)

## 'data.frame':    72 obs. of  5 variables:
##  $ Rep     : Factor w/ 3 levels "1","2","3": 1 1 1 1 1 1 1 1 1 1 ...
##  $ Plate   : Factor w/ 2 levels "1","2": 1 1 1 1 1 1 1 1 1 1 ...
##  $ Well    : Factor w/ 6 levels "1","2","3","4",..: 1 1 1 2 2 2 3 3 3 4
...
##  $ Compound: Factor w/ 8 levels "DMSO 0.6 % (v/v)",..: 3 3 3 5 5 5 4 4 4
 7 ...
##  $ Protein : num  22.1 19.1 18.2 18 14.1 ...
```

#### Statistical model

Hierarchical experimental design, since the plates are nested in the replications and the wells are nested in each plate.

- Linear mixed model
- Protein as response variable
- Compounds as fixed effects
- Rep, Plates and Wells as random effects

```
fit <- lmer(Protein ~ Compound + (1|Rep) + (1|Rep:Plate) + (1|Rep:Plate:Wel
l),
            data=aza)
```

##### Estimated variance components

```
print(VarCorr(fit),comp=c("Variance"),digits=2)

##  Groups         Name        Variance
##  Rep:Plate:Well (Intercept) 0.26
##  Rep:Plate      (Intercept) 0.24
##  Rep            (Intercept) 0.13
##  Residual                   3.96
```

- The residual variance is relatively high, compared to the variance that can be explained due to the experimental design.

```
anova(fit,
      ddf="Kenward-Roger")

## Type III Analysis of Variance Table with Kenward-Roger's method
##          Sum Sq Mean Sq NumDF  DenDF F value   Pr(>F)
## Compound 160.94  22.991     7 10.845  5.3773 0.007245 **
## ---
## Signif. codes:  0 '***' 0.001 '**' 0.01 '*' 0.05 '.' 0.1 ' ' 1
```

- The p-value of 0.007245 is smaller than $\alpha = 0.05$

#### Model based least square means and their 95 % confidence intervals

```
means <- emmeans(fit,
                 specs="Compound",
                 adjust="mvt",
                 type="response")

means

##  Compound                        emmean    SE   df lower.CL upper.CL
##  DMSO 0.6 % (v/v)                  16.9 0.807 14.5     14.4     19.5
##  DMSO 1.2 % (v/v)                  18.9 0.807 14.5     16.3     21.4
##  pycnidione 1                      17.8 0.807 14.5     15.3     20.4
##  eupenifeldin 2                    13.4 0.807 14.5     10.9     15.9
##  xenovulene B 3                    15.1 0.807 14.5     12.5     17.6
##  4-hydroxyxenovulene B 4           14.0 0.807 14.5     11.4     16.5
##  4-dehydroxy norpycnidione 5       15.0 0.807 14.5     12.4     17.5
##  4-hydroxy norxenovulene B 6       15.5 0.807 14.5     13.0     18.1
##
## Degrees-of-freedom method: kenward-roger
## Confidence level used: 0.95
## Conf-level adjustment: mvt method for 8 estimates
```

A graphical overview about this table is given in fig. 2 c.

#### Mean comparisons

Only the mean comparisons defined in the list below were tested

```
cont_list
```

```
## $`DMSO 0.6 % (v/v) - pycnidione 1`
## [1]  1  0 -1  0  0  0  0  0
##
## $`DMSO 0.6 % (v/v) - eupenifeldin 2`
## [1]  1  0  0 -1  0  0  0  0
##
## $`DMSO 0.6 % (v/v) - xenovulene B 3`
## [1]  1  0  0  0 -1  0  0  0
##
## $`DMSO 0.6 % (v/v) - 4-hydroxy norxenovulene B 6`
## [1]  1  0  0  0  0  0  0 -1
##
## $`DMSO 1.2 % (v/v) - 4-hydroxyxenovulene B 4`
## [1]  0  1  0  0  0 -1  0  0
##
## $`DMSO 1.2 % (v/v) - 4-dehydroxy norpycnidione 5`
## [1]  0  1  0  0  0  0 -1  0

comp <- contrast(means,
                 method=cont_list,
                 adjust="mvt")
```

```
comp

##                                          contrast  estimate p.value   si
g
## 1 DMSO 0.6 % (v/v) - 4-hydroxy norxenovulene B 6  1.4027470   0.669 FALS
E
## 2                DMSO 0.6 % (v/v) - eupenifeldin 2  3.5509246   0.046  TRU
E
## 3                  DMSO 0.6 % (v/v) - pycnidione 1 -0.8899113   0.918 FALS
E
## 4                DMSO 0.6 % (v/v) - xenovulene B 3  1.8763319   0.418 FALS
E
## 5 DMSO 1.2 % (v/v) - 4-dehydroxy norpycnidione 5  3.9179278   0.028  TRU
E
## 6     DMSO 1.2 % (v/v) - 4-hydroxyxenovulene B 4  4.9004060   0.008  TRU
E
```

A graphical overview about this table is given in fig. 2 d.

### Statistical supplementary statS4: Cell morphology

Bergmann et al. 2024

#### Packages used for evaluation

```
library(tidyverse)
library(emmeans)
library(lme4)
library(multcomp)
library(tukeytrend)
library(ggpubr)
```

#### Overview about the data

The complete experiment was run in three separate repetitions (Variable trial). For treatment (six compounds plus untreated control), cells were cultured on one well per treatment within each of the trials. For each well, several morphological characteristics were measured on several hundreds of cells (variable n) resulting in a total of 16632 analysed cells. Since several of the measurements resulted in highly right or left skewed observations that were impossible to center by transformation, modelling was done based on the average measurement per well.
This data sets (dat) is depicted below and contains the following variables:

- Comp_ID: ID for the compounds of interest (see below)
- Trial: Indicator variable for the three repetitions
- Well: Indicator variable for the wells in each repetition
- Area_m: Average cell area per well
- Perim_m: Average perimeter per well
- AR_m: Average aspect ratio per well
- Circ_m: Average circularity per well
- Solidity_m: Average solidity per well
- n: Number of cells per well

Compound IDs

- 1 = pycnidione
- 2 = eupenifeldin
- 3 = xenovulene B
- 4 = 4-hydroxyxenovulene B
- 5 = 4-dehydroxy norpycnidione
- 6 = 4-hydroxy norxenovulene B
- U = Untreated

```
dat

##    Comp_ID Trial Well    Area_m   Perim_m     AR_m    Circ_m Solidity_m
   n
## 1        1     1    1  670.6581 127.58322 2.092708 0.5236193  0.8185791
 373
## 2        2     1    2  791.9424 151.75865 1.955226 0.4436452  0.7430968
 310
## 3        3     1    3  975.8344 139.81239 1.913272 0.6078634  0.8770748
 615
```

```
## 4         4     1     4 1048.7376 151.33379 2.057143 0.5621454  0.8521684
 392
## 5         5     1     5  773.6091 130.99050 2.283691 0.5629228  0.8659365
 803
## 6         6     1     6  560.7641 103.78866 1.816892 0.6410123  0.8926472
1301
## 7         U     1     7  490.0072  97.16836 1.840930 0.6410694  0.8923834
1484
## 8         1     2     8  529.2838 117.34503 1.961732 0.4868156  0.7694134
 358
## 9         2     2     9  541.9511 115.49518 2.010395 0.5094737  0.7923684
 228
## 10        3     2    10  832.7332 130.05037 1.912398 0.6061806  0.8706806
 576
## 11        4     2    11 1046.1742 142.11862 1.718805 0.6384778  0.8894157
 676
## 12        5     2    12  700.1952 129.86816 2.498716 0.5276609  0.8444358
 693
## 13        6     2    13  481.7690  97.00257 1.886395 0.6352602  0.8901686
1518
## 14        U     2    14  396.9946  87.20479 1.816864 0.6458562  0.8904478
1773
## 15        1     3    15  489.1608 109.31691 2.004433 0.5105155  0.7915464
 194
## 16        2     3    16  529.8521 115.10853 2.021799 0.5102360  0.7929794
 339
## 17        3     3    17  838.6017 130.16683 2.062396 0.6070312  0.8842899
 545
## 18        4     3    18 1038.0826 142.75380 1.798205 0.6269541  0.8847202
 654
## 19        5     3    19  713.8594 121.16117 2.073190 0.6047402  0.8793225
 893
## 20        6     3    20  523.0936 101.81040 1.991062 0.6268624  0.8916426
1410
## 21        U     3    21  470.2807  95.31744 1.845806 0.6397602  0.8896560
1497
```

#### Modeling

For each of the measured variables, a separate linear model was fit, with Trial and Compound ID as predictor variables

```
# Area
area_fit <- lm(Area_m ~ Trial + Comp_ID, data=dat)

# Perimeter
perim_fit <- lm(Perim_m ~ Trial + Comp_ID, data=dat)

# Aspect ratio
ar_fit <- lm(AR_m ~ Trial + Comp_ID, data=dat)

# Circularity
circ_fit <- lm(Circ_m ~ Trial + Comp_ID, data=dat)
```

```r
# Solidity
Solidity_fit <- lm(Solidity_m ~ Trial + Comp_ID, data=dat)
```

#### Multiple comparisons

Multiple contrast tests are done with respect to the correlations between the dependent variables of the individual models using multiple marginal models.

```r
ni <- as.numeric(table(dat$Comp_ID))
cmt <- contrMat(ni, type="Tukey")

# Multiple marginal models
meandiff <- glht(mmm(area_fit=area_fit,
                     perim_fit=perim_fit,
                     ar_fit=ar_fit,
                     circ_fit=circ_fit,
                     Solidity_fit=Solidity_fit),
                 mlf(mcp(Comp_ID = "Tukey")))

#
meandiff_pval <- fortify(summary(meandiff)) %>%
        mutate(sig=p<0.05) %>%
        dplyr::select(-se, -t) %>%
        data.frame(row.names = NULL)

meandiff_pval

##                      lhs rhs      estimate            p   sig
## 1       area_fit: 2 - 1   0  5.821430e+01 9.567286e-01 FALSE
## 2       area_fit: 3 - 1   0  3.193556e+02 4.022338e-13  TRUE
## 3       area_fit: 4 - 1   0  4.812972e+02 0.000000e+00  TRUE
## 4       area_fit: 5 - 1   0  1.661870e+02 5.624894e-03  TRUE
## 5       area_fit: 6 - 1   0 -4.115868e+01 9.970425e-01 FALSE
## 6       area_fit: U - 1   0 -1.106067e+02 2.625072e-01 FALSE
## 7       area_fit: 3 - 2   0  2.611413e+02 4.043728e-07  TRUE
## 8       area_fit: 4 - 2   0  4.230829e+02 1.409983e-14  TRUE
## 9       area_fit: 5 - 2   0  1.079727e+02 4.786193e-01 FALSE
## 10      area_fit: 6 - 2   0 -9.937298e+01 6.440247e-01 FALSE
## 11      area_fit: U - 2   0 -1.688210e+02 4.168936e-02  TRUE
## 12      area_fit: 4 - 3   0  1.619416e+02 7.112023e-03  TRUE
## 13      area_fit: 5 - 3   0 -1.531686e+02 5.039825e-07  TRUE
## 14      area_fit: 6 - 3   0 -3.605142e+02 0.000000e+00  TRUE
## 15      area_fit: U - 3   0 -4.299623e+02 0.000000e+00  TRUE
## 16      area_fit: 5 - 4   0 -3.151102e+02 1.444178e-12  TRUE
## 17      area_fit: 6 - 4   0 -5.224559e+02 0.000000e+00  TRUE
## 18      area_fit: U - 4   0 -5.919039e+02 0.000000e+00  TRUE
## 19      area_fit: 6 - 5   0 -2.073457e+02 8.782701e-09  TRUE
## 20      area_fit: U - 5   0 -2.767937e+02 4.204970e-12  TRUE
## 21      area_fit: U - 6   0 -6.944802e+01 8.355530e-01 FALSE
## 22     perim_fit: 2 - 1   0  9.372398e+00 8.835281e-01 FALSE
## 23     perim_fit: 3 - 1   0  1.526148e+01 2.544424e-01 FALSE
## 24     perim_fit: 4 - 1   0  2.732035e+01 3.993781e-05  TRUE
## 25     perim_fit: 5 - 1   0  9.258229e+00 8.913982e-01 FALSE
## 26     perim_fit: 6 - 1   0 -1.721450e+01 1.187468e-01 FALSE
## 27     perim_fit: U - 1   0 -2.485152e+01 1.455427e-03  TRUE
```

```
## 28      perim_fit: 3 - 2   0  5.889078e+00 9.993070e-01 FALSE
## 29      perim_fit: 4 - 2   0  1.794796e+01 3.178668e-01 FALSE
## 30      perim_fit: 5 - 2   0 -1.141689e-01 1.000000e+00 FALSE
## 31      perim_fit: 6 - 2   0 -2.658690e+01 2.352845e-02  TRUE
## 32      perim_fit: U - 2   0 -3.422392e+01 3.636029e-04  TRUE
## 33      perim_fit: 4 - 3   0  1.205888e+01 1.428601e-02  TRUE
## 34      perim_fit: 5 - 3   0 -6.003247e+00 9.932483e-01 FALSE
## 35      perim_fit: 6 - 3   0 -3.247598e+01 2.168632e-08  TRUE
## 36      perim_fit: U - 3   0 -4.011300e+01 4.918288e-12  TRUE
## 37      perim_fit: 5 - 4   0 -1.806213e+01 6.855940e-02 FALSE
## 38      perim_fit: 6 - 4   0 -4.453486e+01 1.643130e-14  TRUE
## 39      perim_fit: U - 4   0 -5.217188e+01 0.000000e+00  TRUE
## 40      perim_fit: 6 - 5   0 -2.647273e+01 3.959769e-03  TRUE
## 41      perim_fit: U - 5   0 -3.410975e+01 6.358882e-06  TRUE
## 42      perim_fit: U - 6   0 -7.637018e+00 9.883507e-01 FALSE
## 43         ar_fit: 2 - 1   0 -2.381755e-02 1.000000e+00 FALSE
## 44         ar_fit: 3 - 1   0 -5.693572e-02 9.999872e-01 FALSE
## 45         ar_fit: 4 - 1   0 -1.615734e-01 9.005095e-01 FALSE
## 46         ar_fit: 5 - 1   0  2.655749e-01 2.741016e-01 FALSE
## 47         ar_fit: 6 - 1   0 -1.215077e-01 9.856159e-01 FALSE
## 48         ar_fit: U - 1   0 -1.850910e-01 7.897582e-01 FALSE
## 49         ar_fit: 3 - 2   0 -3.311817e-02 1.000000e+00 FALSE
## 50         ar_fit: 4 - 2   0 -1.377558e-01 9.926260e-01 FALSE
## 51         ar_fit: 5 - 2   0  2.893924e-01 5.159686e-01 FALSE
## 52         ar_fit: 6 - 2   0 -9.769018e-02 9.991979e-01 FALSE
## 53         ar_fit: U - 2   0 -1.612734e-01 6.096741e-01 FALSE
## 54         ar_fit: 4 - 3   0 -1.046377e-01 9.995593e-01 FALSE
## 55         ar_fit: 5 - 3   0  3.225106e-01 4.063017e-01 FALSE
## 56         ar_fit: 6 - 3   0 -6.457201e-02 9.999947e-01 FALSE
## 57         ar_fit: U - 3   0 -1.281552e-01 9.813431e-01 FALSE
## 58         ar_fit: 5 - 4   0  4.271483e-01 9.221319e-02 FALSE
## 59         ar_fit: 6 - 4   0  4.006564e-02 1.000000e+00 FALSE
## 60         ar_fit: U - 4   0 -2.351759e-02 1.000000e+00 FALSE
## 61         ar_fit: 6 - 5   0 -3.870826e-01 1.573353e-01 FALSE
## 62         ar_fit: U - 5   0 -4.506658e-01 2.169425e-02  TRUE
## 63         ar_fit: U - 6   0 -6.358323e-02 9.999716e-01 FALSE
## 64       circ_fit: 2 - 1   0 -1.919853e-02 9.987607e-01 FALSE
## 65       circ_fit: 3 - 1   0  1.000416e-01 1.667472e-04  TRUE
## 66       circ_fit: 4 - 1   0  1.022090e-01 2.988963e-04  TRUE
## 67       circ_fit: 5 - 1   0  5.812449e-02 2.300002e-01 FALSE
## 68       circ_fit: 6 - 1   0  1.273948e-01 5.310850e-08  TRUE
## 69       circ_fit: U - 1   0  1.352451e-01 5.615386e-09  TRUE
## 70       circ_fit: 3 - 2   0  1.192401e-01 3.600350e-08  TRUE
## 71       circ_fit: 4 - 2   0  1.214075e-01 1.079894e-05  TRUE
## 72       circ_fit: 5 - 2   0  7.732302e-02 7.307088e-02 FALSE
## 73       circ_fit: 6 - 2   0  1.465934e-01 1.305033e-10  TRUE
## 74       circ_fit: U - 2   0  1.544436e-01 3.895773e-13  TRUE
## 75       circ_fit: 4 - 3   0  2.167395e-03 1.000000e+00 FALSE
## 76       circ_fit: 5 - 3   0 -4.191709e-02 6.690935e-01 FALSE
## 77       circ_fit: 6 - 3   0  2.735325e-02 8.217284e-01 FALSE
## 78       circ_fit: U - 3   0  3.520354e-02 2.056416e-01 FALSE
## 79       circ_fit: 5 - 4   0 -4.408449e-02 8.120621e-01 FALSE
## 80       circ_fit: 6 - 4   0  2.518586e-02 9.904561e-01 FALSE
## 81       circ_fit: U - 4   0  3.303614e-02 9.060071e-01 FALSE
```

```
## 82       circ_fit: 6 - 5   0  6.927034e-02 8.670402e-02 FALSE
## 83       circ_fit: U - 5   0  7.712063e-02 1.507736e-02  TRUE
## 84       circ_fit: U - 6   0  7.850283e-03 9.999964e-01 FALSE
## 85   Solidity_fit: 2 - 1   0 -1.703145e-02 9.872190e-01 FALSE
## 86   Solidity_fit: 3 - 1   0  8.416879e-02 1.281805e-07  TRUE
## 87   Solidity_fit: 4 - 1   0  8.225511e-02 2.536434e-07  TRUE
## 88   Solidity_fit: 5 - 1   0  7.005196e-02 2.622702e-05  TRUE
## 89   Solidity_fit: 6 - 1   0  9.830650e-02 2.221284e-10  TRUE
## 90   Solidity_fit: U - 1   0  9.764945e-02 3.176529e-10  TRUE
## 91   Solidity_fit: 3 - 2   0  1.012002e-01 0.000000e+00  TRUE
## 92   Solidity_fit: 4 - 2   0  9.928656e-02 1.559530e-12  TRUE
## 93   Solidity_fit: 5 - 2   0  8.708341e-02 6.960377e-11  TRUE
## 94   Solidity_fit: 6 - 2   0  1.153379e-01 0.000000e+00  TRUE
## 95   Solidity_fit: U - 2   0  1.146809e-01 0.000000e+00  TRUE
## 96   Solidity_fit: 4 - 3   0 -1.913676e-03 1.000000e+00 FALSE
## 97   Solidity_fit: 5 - 3   0 -1.411683e-02 7.116117e-01 FALSE
## 98   Solidity_fit: 6 - 3   0  1.413771e-02 1.164414e-02  TRUE
## 99   Solidity_fit: U - 3   0  1.348066e-02 7.182251e-02 FALSE
## 100  Solidity_fit: 5 - 4   0 -1.220315e-02 9.907477e-01 FALSE
## 101  Solidity_fit: 6 - 4   0  1.605139e-02 8.381229e-01 FALSE
## 102  Solidity_fit: U - 4   0  1.539433e-02 8.820907e-01 FALSE
## 103  Solidity_fit: 6 - 5   0  2.825454e-02 1.518039e-02  TRUE
## 104  Solidity_fit: U - 5   0  2.759748e-02 3.003614e-02  TRUE
## 105  Solidity_fit: U - 6   0 -6.570534e-04 1.000000e+00 FALSE
```

#### Graphical overview about the multiple comparisons

The graphic displays the estimated differences between the different means estimated based on the models shown above $\hat{\mu}_i - \hat{\mu}_{i'}$ (black dots). The error bars indicate 95 % simultaneous confidence intervals for the mean differences. The black line indicates the null hypothesis $H_0: \mu_i - \mu_{i'} = 0$. Hence, mean differences do not significantly differ from zero, if their confidence intervals encompass zero.

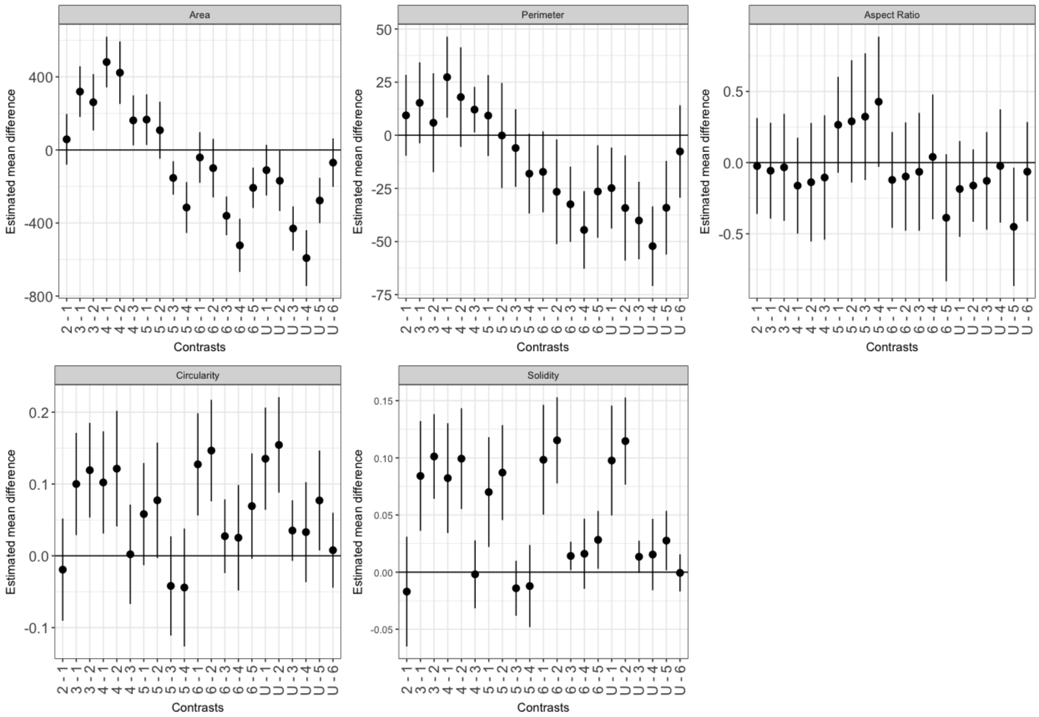

#### Statistical supplementary statS5: Branches

Bergmann et al. 2024

##### Packages used for evaluation

```
library(tidyverse)
library(emmeans)
library(lme4)
library(multcomp)
library(tukeytrend)
library(ggpubr)
```

##### Overview about the data

The complete experiment was run in three separate repetitions (Variable trial). For treatment (six compounds plus untreated control), cells were cultured on one well per treatment within each of the trials. For each well, the number of branches and the number of junctions was counted on several hundreds of cells (variable n) resulting in a total of 16632 analysed cells. Modelling was done based on the average measurement per well.
This data sets (dat) is depicted below and contains the following variables:

- Compound: Compound of interest
- Comp_ID: ID for the compounds of interest
- Trial: Indicator variable for the three repetitions
- Well: Indicator variable for the wells in each repetition
- Branches_m: Average number of branches area per well
- Junctions_m: Average number of junctions per well
- n: Number of cells per well

```
##                    Compound Comp_ID Trial Well Branches_m Junctions_m
n
## 1                pycnidione       1     1    1   1.828418  0.41286863

373
## 2              eupenifeldin       2     1    2   3.035484  1.01290323

310
## 3              xenovulene B       3     1    3   1.304065  0.15121951

615
## 4       hydroxyxenovulene B       4     1    4   1.612245  0.30612245

392
## 5   dehydroxy norpycnidione       5     1    5   1.264010  0.13325031

803
## 6   hydroxy norxenovulene B       6     1    6   1.138355  0.07225211

1301
## 7                 untreated       U     1    7   1.105795  0.05727763

1484
## 8                pycnidione       1     2    8   1.997207  0.49441341
```

```
358
## 9               eupenifeldin       2     2    9    2.043860   0.52192982

228
## 10              xenovulene B       3     2   10    1.291667   0.14583333

576
## 11      hydroxyxenovulene B       4     2   11    1.196746   0.09763314

676
## 12 dehydroxy norpycnidione       5     2   12    1.248196   0.12409812

693
## 13 hydroxy norxenovulene B       6     2   13    1.113966   0.05928854

1518
## 14                 untreated      U     2   14    1.120135   0.06373378

1773
## 15                pycnidione      1     3   15    1.891753   0.44329897

194
## 16              eupenifeldin      2     3   16    2.029499   0.51032448

339
## 17              xenovulene B      3     3   17    1.255046   0.12660550

545
## 18       hydroxyxenovulene B      4     3   18    1.238532   0.12079511

654
## 19 dehydroxy norpycnidione       5     3   19    1.178052   0.09182531

893
## 20 hydroxy norxenovulene B       6     3   20    1.097872   0.05319149

1410
## 21                 untreated      U     3   21    1.122244   0.06680027

1497
```

#### Modeling

For each of the measured variables, a separate linear model was fit, with Trial and Compound ID as predictor variables

```
# Area
branches_fit <- lm(Branches_m ~ Trial + Comp_ID, data=dat)

# Perimeter
junctions_fit <- lm(Junctions_m ~ Trial + Comp_ID, data=dat)
```

#### Multiple comparisons

Multiple contrast tests are done with respect to the correlations between the dependent variables of the individual models using multiple marginal models.

```r
ni <- as.numeric(table(dat$Comp_ID))
cmt <- contrMat(ni, type="Tukey")

# Multiple marginal models
meandiff <- glht(mmm(branches_fit=branches_fit,
                     junctions_fit=junctions_fit),
                 mlf(mcp(Comp_ID = "Tukey")))

#
meandiff_pval <- fortify(summary(meandiff)) %>%
        mutate(sig=p<0.05) %>%
        dplyr::select(-se, -t) %>%
        data.frame(row.names = NULL)

meandiff_pval
```

```
##                       lhs rhs      estimate            p   sig
## 1   branches_fit: 2 - 1   0  0.4638215111 1.428023e-01 FALSE
## 2   branches_fit: 3 - 1   0 -0.6221999776 1.122805e-02  TRUE
## 3   branches_fit: 4 - 1   0 -0.5566183139 3.676095e-02  TRUE
## 4   branches_fit: 5 - 1   0 -0.6757065964 4.376842e-03  TRUE
## 5   branches_fit: 6 - 1   0 -0.7890614385 3.379284e-04  TRUE
## 6   branches_fit: U - 1   0 -0.7897341703 3.398045e-04  TRUE
## 7   branches_fit: 3 - 2   0 -1.0860214888 2.849172e-06  TRUE
## 8   branches_fit: 4 - 2   0 -1.0204398250 1.139823e-04  TRUE
## 9   branches_fit: 5 - 2   0 -1.1395281075 7.583477e-07  TRUE
## 10  branches_fit: 6 - 2   0 -1.2528829496 5.874713e-08  TRUE
## 11  branches_fit: U - 2   0 -1.2535556814 4.073595e-08  TRUE
## 12  branches_fit: 4 - 3   0  0.0655816638 9.988450e-01 FALSE
## 13  branches_fit: 5 - 3   0 -0.0535066188 9.992345e-01 FALSE
## 14  branches_fit: 6 - 3   0 -0.1668614609 7.929603e-01 FALSE
## 15  branches_fit: U - 3   0 -0.1675341926 8.639582e-01 FALSE
## 16  branches_fit: 5 - 4   0 -0.1190882825 9.651053e-01 FALSE
## 17  branches_fit: 6 - 4   0 -0.2324431246 5.490846e-01 FALSE
## 18  branches_fit: U - 4   0 -0.2331158564 6.490952e-01 FALSE
## 19  branches_fit: 6 - 5   0 -0.1133548421 9.538080e-01 FALSE
## 20  branches_fit: U - 5   0 -0.1140275739 9.737858e-01 FALSE
## 21  branches_fit: U - 6   0 -0.0006727318 1.000000e+00 FALSE
## 22 junctions_fit: 2 - 1   0  0.2315255082 1.387972e-01 FALSE
## 23 junctions_fit: 3 - 1   0 -0.3089742198 1.162076e-02  TRUE
## 24 junctions_fit: 4 - 1   0 -0.2753434392 3.862750e-02  TRUE
## 25 junctions_fit: 5 - 1   0 -0.3338024221 4.386203e-03  TRUE
## 26 junctions_fit: 6 - 1   0 -0.3886162896 2.324123e-04  TRUE
## 27 junctions_fit: U - 1   0 -0.3875897766 3.191971e-04  TRUE
## 28 junctions_fit: 3 - 2   0 -0.5404997280 3.152466e-06  TRUE
## 29 junctions_fit: 4 - 2   0 -0.5068689473 2.144680e-05  TRUE
## 30 junctions_fit: 5 - 2   0 -0.5653279303 7.437242e-07  TRUE
## 31 junctions_fit: 6 - 2   0 -0.6201417978 3.168100e-08  TRUE
## 32 junctions_fit: U - 2   0 -0.6191152848 6.963035e-08  TRUE
## 33 junctions_fit: 4 - 3   0  0.0336307807 9.986919e-01 FALSE
## 34 junctions_fit: 5 - 3   0 -0.0248282022 9.995081e-01 FALSE
## 35 junctions_fit: 6 - 3   0 -0.0796420698 8.313510e-01 FALSE
## 36 junctions_fit: U - 3   0 -0.0786155568 8.985300e-01 FALSE
## 37 junctions_fit: 5 - 4   0 -0.0584589829 9.689262e-01 FALSE
```

```
## 38 junctions_fit: 6 - 4   0 -0.1132728505 5.877227e-01 FALSE
## 39 junctions_fit: U - 4   0 -0.1122463374 6.955207e-01 FALSE
## 40 junctions_fit: 6 - 5   0 -0.0548138676 9.618961e-01 FALSE
## 41 junctions_fit: U - 5   0 -0.0537873545 9.810913e-01 FALSE
## 42 junctions_fit: U - 6   0  0.0010265130 1.000000e+00 FALSE
```

#### Graphical overview about the multiple comparisons

The graphic displays the estimated differences between the different means estimated based on the models shown above $\hat{\mu}_i - \hat{\mu}_{i'}$ (black dots). The error bars indicate 95 % simultaneous confidence intervals for the mean differences. The black line indicates the null hypothesis $H_0: \mu_i - \mu_{i'} = 0$. Hence, mean differences do not significantly differ from zero, if their confidence intervals encompass zero.

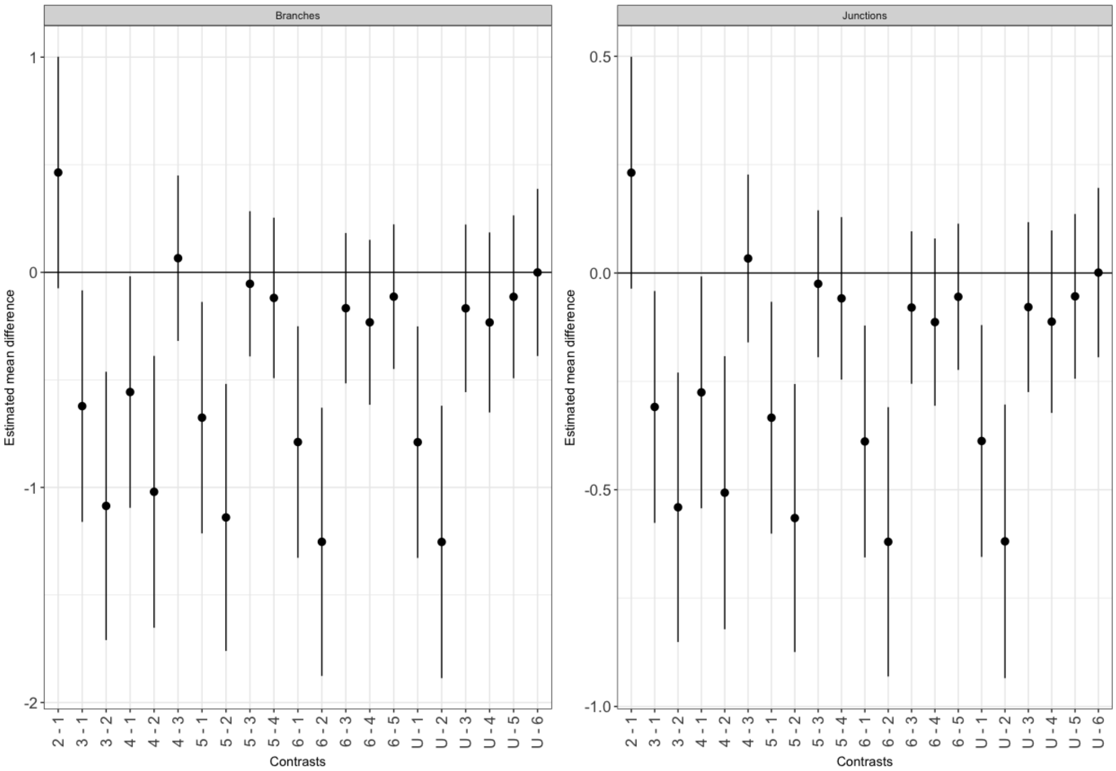

#### Statistical supplementary statS6: Proliferation for 4-hydroxy norxenovulene B 6 and DMSO

Bergmann et al. 2024

```
# Load the packages used for evaluation
library(lme4)
library(lmerTest)
library(tidyverse)
library(emmeans)
library(multcomp)
```

##### Data set

- Treatment: Name of the treatments (combination of compound and dose)
- Rep: Repetition (timepoint)
- Plate: Plate per timepoint
- Prop: Proportion of the fluorescence signals at the start of the experiment and its end
- Fluo1: Fluorescence signals at the start of the experiments (day one)
- Fluo3: Fluorescence signals at the end of the experiments (day three)

```
proli7

##                           Treatment Rep Plate      Prop Fluo1 Fluo3
## 1                         untreated   1     1 12.220345 33276  2723
## 2   4-hydroxy norxenovulene B 6:4   1     1 12.148959 33276  2739
## 3   4-hydroxy norxenovulene B 6:6   1     1 11.867332 33276  2804
## 4   4-hydroxy norxenovulene B 6:8   1     1 12.052155 33276  2761
## 5  4-hydroxy norxenovulene B 6:12   1     1 11.679888 33276  2849
## 6                         untreated   1     2 12.338154 33276  2697
## 7                          DMSO:0.4   1     2 12.338154 33276  2697
## 8                          DMSO:0.6   1     2 12.078403 33276  2755
## 9                          DMSO:0.8   1     2 12.434978 33276  2676
## 10                         DMSO:1.2   1     2 11.522161 33276  2888
## 11                        untreated   2     3  9.097393 36990  4066
## 12  4-hydroxy norxenovulene B 6:4   2     3  9.153675 36990  4041
## 13  4-hydroxy norxenovulene B 6:6   2     3  8.838710 36990  4185
## 14  4-hydroxy norxenovulene B 6:8   2     3  8.784137 36990  4211
## 15 4-hydroxy norxenovulene B 6:12   2     3  8.534841 36990  4334
## 16                        untreated   2     4  9.097393 36990  4066
## 17                         DMSO:0.4   2     4  9.286970 36990  3983
## 18                         DMSO:0.6   2     4  9.249812 36990  3999
## 19                         DMSO:0.8   2     4  9.097393 36990  4066
## 20                         DMSO:1.2   2     4  8.586351 36990  4308
## 21                        untreated   3     5 11.146008 42444  3808
## 22  4-hydroxy norxenovulene B 6:4   3     5 11.102276 42444  3823
## 23  4-hydroxy norxenovulene B 6:6   3     5 10.967442 42444  3870
## 24  4-hydroxy norxenovulene B 6:8   3     5 10.154067 42444  4180
## 25 4-hydroxy norxenovulene B 6:12   3     5  9.977433 42444  4254
## 26                        untreated   3     6 11.146008 42444  3808
## 27                         DMSO:0.4   3     6 10.699269 42444  3967
## 28                         DMSO:0.6   3     6 10.767123 42444  3942
## 29                         DMSO:0.8   3     6 10.354721 42444  4099
## 30                         DMSO:1.2   3     6  9.872994 42444  4299
```

##### Statistical model

- Linear mixed effects model
- Fluorescence proportion as response variable (ln-transformed)
- Repetition and Treatment as fixed effects
- Plates nested in Repetition as normal distributed random effects

```
fit_p7 <- lmer(log(Prop) ~ Rep + Treatment  + (1|Rep:Plate), data=proli7)
```

##### Model based means and their pointwise 95% confidence intervals

- All mean comparisons were performed on the ln-scale
- The means and their CI are back transformed from the ln-scale to the response scale

```
means <- emmeans(fit_p7,
                 specs="Treatment",
                 type="response")

means

##                          Treatment  response         SE  lower.CL  upper.CL
## 1                        untreated 10.758070 0.09157412 10.566103 10.95353
## 2  4-hydroxy norxenovulene B 6:4 10.728754 0.13274648 10.452111 11.01272
## 3  4-hydroxy norxenovulene B 6:6 10.478912 0.12965520 10.208712 10.75626
## 4  4-hydroxy norxenovulene B 6:8 10.244758 0.12675801  9.980595 10.51591
## 5 4-hydroxy norxenovulene B 6:12  9.982778 0.12351654  9.725370 10.24700
## 6                         DMSO:0.4 10.701854 0.13241364 10.425904 10.98511
## 7                         DMSO:0.6 10.634415 0.13157923 10.360205 10.91588
## 8                         DMSO:0.8 10.540625 0.13041876 10.268833 10.81961
## 9                         DMSO:1.2  9.921181 0.12275441  9.665362 10.18377
```

A graphical overview about this table is given in the supplementary figure S 5 c.

##### Mean comparisons (contrast tests)

Pairwise mean comparisons, that are defined below by `cont_list`, were run based on mean differences $ln(\hat{\mu}_j) - ln(\hat{\mu}_{j'})$. The tested null-hypothesis for each comparison was $H_0: ln(\mu_j) - ln(\mu_{j'}) = 0$. The table below shows results from the tests that are already back transformed to the response scale. Hence the back transformed contrasts become proportions on the response scale $exp\left(ln(\hat{\mu}_j) - ln(\hat{\mu}_{j'})\right) = \hat{\mu}_j/\hat{\mu}_{j'}$. Similarly, the $H_0$ for the back transformed contrasts changes to $H_0: \mu_j/\mu_{j'} = 1$

```
# List of contrast coefficients
cont_list

## $`4-hydroxy norxenovulene B 6:4 - untreated`
## [1] -1  1  0  0  0  0  0  0  0
##
## $`4-hydroxy norxenovulene B 6:6 - untreated`
## [1] -1  0  1  0  0  0  0  0  0
##
## $`4-hydroxy norxenovulene B 6:8 - untreated`
## [1] -1  0  0  1  0  0  0  0  0
##
## $`4-hydroxy norxenovulene B 6:12 - untreated`
```

```
## [1] -1  0  0  0  1  0  0  0  0
##
## $`DMSO:0.4 - untreated`
## [1] -1  0  0  0  0  1  0  0  0
##
## $`DMSO:0.6 - untreated`
## [1] -1  0  0  0  0  0  1  0  0
##
## $`DMSO:0.8 - untreated`
## [1] -1  0  0  0  0  0  0  1  0
##
## $`DMSO:1.2 - untreated`
## [1] -1  0  0  0  0  0  0  0  1
##
## $`DMSO:0.4 - 4-hydroxy norxenovulene B 6:4`
## [1]  0 -1  0  0  0  1  0  0  0
##
## $`DMSO:0.6 - 4-hydroxy norxenovulene B 6:6`
## [1]  0  0 -1  0  0  0  1  0  0
##
## $`DMSO:0.8 - 4-hydroxy norxenovulene B 6:8`
## [1]  0  0  0 -1  0  0  0  1  0
##
## $`DMSO:1.2 - 4-hydroxy norxenovulene B 6:12`
## [1]  0  0  0  0 -1  0  0  0  1
```

```
# Multiple contrast tests
comp <- contrast(means,
                 method=cont_list,
                 adjust="mvt")
```

```
##                                      contrast     ratio p.value   sig
## 1  4-hydroxy norxenovulene B 6:12 / untreated 0.9279339   0.001  TRUE
## 2   4-hydroxy norxenovulene B 6:4 / untreated 0.9972749  >0.999 FALSE
## 3   4-hydroxy norxenovulene B 6:6 / untreated 0.9740513   0.522 FALSE
## 4   4-hydroxy norxenovulene B 6:8 / untreated 0.9522858   0.036  TRUE
## 5    DMSO:0.4 / 4-hydroxy norxenovulene B 6:4 0.9974927  >0.999 FALSE
## 6                        DMSO:0.4 / untreated 0.9947745  >0.999 FALSE
## 7    DMSO:0.6 / 4-hydroxy norxenovulene B 6:6 1.0148396   0.981 FALSE
## 8                        DMSO:0.6 / untreated 0.9885058   0.986 FALSE
## 9    DMSO:0.8 / 4-hydroxy norxenovulene B 6:8 1.0288798   0.648 FALSE
## 10                       DMSO:0.8 / untreated 0.9797877   0.775 FALSE
## 11  DMSO:1.2 / 4-hydroxy norxenovulene B 6:12 0.9938297  >0.999 FALSE
## 12                       DMSO:1.2 / untreated 0.9222082  <0.001  TRUE
```

The contrast ratios, their corresponding p-values and their confidence intervals (table below) are shown in supplementary figure S 5 d.

```
confint(comp)
```

```
##  contrast                                   ratio     SE   df lower.CL u
pper.CL
##  4-hydroxy norxenovulene B 6:4 / untreated  0.997 0.0147 18.6    0.952
  1.045
##  4-hydroxy norxenovulene B 6:6 / untreated  0.974 0.0144 18.6    0.930
```

```
  1.020
##  4-hydroxy norxenovulene B 6:8 / untreated  0.952 0.0141 18.6    0.909
  0.998
##  4-hydroxy norxenovulene B 6:12 / untreated 0.928 0.0137 18.6    0.886
  0.972
##  DMSO:0.4 / untreated                       0.995 0.0147 18.6    0.950
  1.042
##  DMSO:0.6 / untreated                       0.989 0.0146 18.6    0.944
  1.036
##  DMSO:0.8 / untreated                       0.980 0.0145 18.6    0.935
  1.026
##  DMSO:1.2 / untreated                       0.922 0.0136 18.6    0.880
  0.966
##  DMSO:0.4 / 4-hydroxy norxenovulene B 6:4   0.997 0.0179 17.2    0.942
  1.056
##  DMSO:0.6 / 4-hydroxy norxenovulene B 6:6   1.015 0.0182 17.2    0.959
  1.074
##  DMSO:0.8 / 4-hydroxy norxenovulene B 6:8   1.029 0.0185 17.2    0.972
  1.089
##  DMSO:1.2 / 4-hydroxy norxenovulene B 6:12  0.994 0.0178 17.2    0.939
  1.052
##
## Results are averaged over the levels of: Rep
## Degrees-of-freedom method: kenward-roger
## Confidence level used: 0.95
## Conf-level adjustment: mvt method for 12 estimates
## Intervals are back-transformed from the log scale
```

### Statistical supplementary statS7: Proliferation

Bergmann et al. 2024

```
# Load the packages used for evaluation
library(tidyverse)
library(emmeans)
library(multcomp)
```

#### Data set

- Compound: Name of the treatments
- Rep: Repetition (timepoint)
- Prop: Proportions of the fluorescence signals at the start of the experiment and its end
- Fluo1: Fluorescence signals at the start of the experiments (day one)
- Fluo3: Fluorescence signals at the end of the experiments (day three)

```
proli

##                         Compound Rep      Prop Fluo1 Fluo3
## 1                   xenovulene B 3   1  5.611080 59758 10650
## 2                   eupenifeldin 2   1  5.509681 59758 10846
## 3   4-dehydroxy norpycnidione 5   1  5.765920 59758 10364
## 4       4-hydroxyxenovulene B 4   1  5.857479 59758 10202
## 5                     pycnidione 1   1  5.870714 59758 10179
## 6   4-hydroxy norxenovulene B 6   1 12.898338 59758  4633
## 7                      untreated   1 14.221323 59758  4202
## 8                   xenovulene B 3   2  6.730180 64603  9599
## 9                   eupenifeldin 2   2  6.730180 64603  9599
## 10  4-dehydroxy norpycnidione 5   2  6.346694 64603 10179
## 11      4-hydroxyxenovulene B 4   2  7.140820 64603  9047
## 12                    pycnidione 1   2  6.899082 64603  9364
## 13  4-hydroxy norxenovulene B 6   2 14.968258 64603  4316
## 14                     untreated   2 15.888588 64603  4066
## 15                  xenovulene B 3   3  6.417296 50164  7817
## 16                  eupenifeldin 2   3  9.086035 50164  5521
## 17  4-dehydroxy norpycnidione 5   3  6.590121 50164  7612
## 18      4-hydroxyxenovulene B 4   3  6.796369 50164  7381
## 19                    pycnidione 1   3  7.598304 50164  6602
## 20  4-hydroxy norxenovulene B 6   3 15.178215 50164  3305
## 21                     untreated   3 15.607965 50164  3214
```

#### Linear model

The data can be treated as a block-design with replicates as blocks and compund as variable of interest.

- Rep as blocks
- Compund as treatments
- Prop (ln-transformed proportions) as dependent variable

```
fit <- lm(log(Prop) ~ Rep + Compound, data=proli)
```

The fitted model is $ln(y_{ij}) = \hat{\mu} + \hat{r}_i + \hat{c}_j + \hat{e}_{ij}$ with:

- $ln(y_{ij})$: Ln-transformed proportions (signal on day 1 / signal on day 3)
- $\hat{\mu}$: Fitted overall mean proportion
- $\hat{r}_i$: Observed effects due to the replication, with $i = 1,2$
- $\hat{c}_j$: Observed effects of the compounds, with $j = 1, \dots, 7$
- $\hat{e}_{ij}$: Observed residuals

#### Analysis of variance

```
anova(fit)

## Analysis of Variance Table
##
## Response: log(Prop)
##           Df  Sum Sq Mean Sq F value    Pr(>F)
## Rep        2 0.16006 0.08003  13.226 0.0009237 ***
## Compound   6 2.87086 0.47848  79.078 6.044e-09 ***
## Residuals 12 0.07261 0.00605
## ---
## Signif. codes:  0 '***' 0.001 '**' 0.01 '*' 0.05 '.' 0.1 ' ' 1
```

- The compound is significant ($\alpha = 0.05 > p = 6.044e - 09$)

#### Model based least square means and their pointwise 95% confidence intervals

- All mean comparisons were performed on the ln-scale
- The means and their CI are back transformed from the ln-scale to the response scale

```
means <- emmeans(fit,
                 specs="Compound",
                 type="response")

##                     Compound  response        SE df  lower.CL  upper.CL
## 1                  untreated 15.221491 0.6835946 12 13.802617 16.786222
## 2               pycnidione 1  6.751490 0.3032083 12  6.122148  7.445526
## 3             eupenifeldin 2  6.958397 0.3125004 12  6.309768  7.673702
## 4             xenovulene B 3  6.234597 0.2799947 12  5.653438  6.875498
## 5     4-hydroxyxenovulene B 4  6.575245 0.2952931 12  5.962332  7.251163
## 6 4-dehydroxy norpycnidione 5  6.224482 0.2795404 12  5.644266  6.864343
## 7 4-hydroxy norxenovulene B 6 14.310075 0.6426630 12 12.976158 15.781114
```

A graphical overview about this table is given in the manuscript in figure 4 a.

#### Mean comparisons (contrast tests)

Pairwise mean comparisons were run based on mean differences $ln(\hat{\mu}_j) - ln(\hat{\mu}_{j'})$. The tested null-hypothesis for each comparison was $H_0: ln(\mu_j) - ln(\mu_{j'}) = 0$. The table below shows results from the tests that are already back transformed to the response scale. Hence, the back transformed contrasts become proportions on the response scale

$exp\left(ln(\hat{\mu}_j) - ln(\hat{\mu}_{j'})\right) = \hat{\mu}_j/\hat{\mu}_{j'}$. Similarly, the $H_0$ for the back transformed contrasts changes to $H_0$: $\mu_j/\mu_{j'} = 1$

```
# Contrasts are given on the response scale
comp <- contrast(means,
                method="pairwise",
                adjust="mvt")
```

```
##                                                          contrast ratio p
.value
## 1                                          untreated / pycnidione 1  2.25
<0.001
## 2                                        untreated / eupenifeldin 2  2.19
<0.001
## 3                                        untreated / xenovulene B 3  2.44
<0.001
## 4                             untreated / (4-hydroxyxenovulene B 4)  2.31
<0.001
## 5                         untreated / (4-dehydroxy norpycnidione 5)  2.45
<0.001
## 6                         untreated / (4-hydroxy norxenovulene B 6)  1.06
 0.951
## 7                                     pycnidione 1 / eupenifeldin 2  0.97
 0.999
## 8                                     pycnidione 1 / xenovulene B 3  1.08
  0.86
## 9                          pycnidione 1 / (4-hydroxyxenovulene B 4)  1.03
 0.999
## 10                     pycnidione 1 / (4-dehydroxy norpycnidione 5)  1.08
 0.849
## 11                     pycnidione 1 / (4-hydroxy norxenovulene B 6)  0.47
<0.001
## 12                                  eupenifeldin 2 / xenovulene B 3  1.12
 0.612
## 13                       eupenifeldin 2 / (4-hydroxyxenovulene B 4)  1.06
 0.967
## 14                   eupenifeldin 2 / (4-dehydroxy norpycnidione 5)  1.12
 0.597
## 15                   eupenifeldin 2 / (4-hydroxy norxenovulene B 6)  0.49
<0.001
## 16                       xenovulene B 3 / (4-hydroxyxenovulene B 4)  0.95
 0.976
## 17                   xenovulene B 3 / (4-dehydroxy norpycnidione 5)  1.00
>0.999
## 18                   xenovulene B 3 / (4-hydroxy norxenovulene B 6)  0.44
<0.001
## 19      (4-hydroxyxenovulene B 4) / (4-dehydroxy norpycnidione 5)  1.06
 0.972
## 20      (4-hydroxyxenovulene B 4) / (4-hydroxy norxenovulene B 6)  0.46
<0.001
## 21 (4-dehydroxy norpycnidione 5) / (4-hydroxy norxenovulene B 6)  0.43
<0.001
```

Results of the mean comparisons are summarized in the manuscript in figure 4 a. The contrast ratios, their corresponding p-values and their confidence intervals (table below) are shown in figure 4 b in the manuscript.

```
# 95% simultanious confidence intervals for the contrasts on response scale
confint(contrast(means,
                 method="pairwise",
                 adjust="mvt"))

##                                                            contrast  LCL  U
CL
## 1                                       untreated / pycnidione 1 1.81 2.
82
## 2                                     untreated / eupenifeldin 2 1.75 2.
73
## 3                                     untreated / xenovulene B 3 1.95 3.
05
## 4                          untreated / (4-hydroxyxenovulene B 4) 1.85 2.
89
## 5                      untreated / (4-dehydroxy norpycnidione 5) 1.96 3.
05
## 6                      untreated / (4-hydroxy norxenovulene B 6) 0.85 1.
33
## 7                                  pycnidione 1 / eupenifeldin 2 0.78 1.
21
## 8                                  pycnidione 1 / xenovulene B 3 0.87 1.
35
## 9                       pycnidione 1 / (4-hydroxyxenovulene B 4) 0.82 1.
28
## 10                  pycnidione 1 / (4-dehydroxy norpycnidione 5) 0.87 1.
35
## 11                  pycnidione 1 / (4-hydroxy norxenovulene B 6) 0.38 0.
59
## 12                               eupenifeldin 2 / xenovulene B 3 0.89 1.
39
## 13                    eupenifeldin 2 / (4-hydroxyxenovulene B 4) 0.85 1.
32
## 14                eupenifeldin 2 / (4-dehydroxy norpycnidione 5) 0.90 1.
40
## 15                eupenifeldin 2 / (4-hydroxy norxenovulene B 6) 0.39 0.
61
## 16                    xenovulene B 3 / (4-hydroxyxenovulene B 4) 0.76 1.
18
## 17                xenovulene B 3 / (4-dehydroxy norpycnidione 5) 0.80 1.
25
## 18                xenovulene B 3 / (4-hydroxy norxenovulene B 6) 0.35 0.
54
## 19     (4-hydroxyxenovulene B 4) / (4-dehydroxy norpycnidione 5) 0.85 1.
32
## 20     (4-hydroxyxenovulene B 4) / (4-hydroxy norxenovulene B 6) 0.37 0.
57
## 21 (4-dehydroxy norpycnidione 5) / (4-hydroxy norxenovulene B 6) 0.35 0.
54
```

### Statistical supplementary statS8: Toxicity for 4-hydroxy norxenovulene B 6 and DMSO

Bergmann et al 2024

```r
# Load the packages used for evaluation
library(tidyverse)
library(lme4)
library(lmerTest)
library(emmeans)
library(multcomp)
library(multcompView)
```

#### Data set

- Treatment: Name of the treatments (combination compound and dose)
- Rep: Repetition (timepoint)
- Plate: Plate per timepoint
- Prop: Proportion of the dead cells
- Dead: Number of dead cells,
- Alive: Number of living cells
- Total: Total number of cells

```
##                         Treatment Rep Plate Alive        Prop Total Dead
## 1                       untreated   1     1  9968 0.001502554  9983   15
## 2   4-hydroxy norxenovulene B 6:4   1     1  9295 0.069662696  9991  696
## 3   4-hydroxy norxenovulene B 6:6   1     1  9840 0.014916408  9989  149
## 4   4-hydroxy norxenovulene B 6:8   1     1  9627 0.036046861  9987  360
## 5  4-hydroxy norxenovulene B 6:12   1     1  9422 0.055627944  9977  555
## 6                       untreated   1     2  9973 0.001801621  9991   18
## 7                        DMSO:0.4   1     2  9760 0.023023023  9990  230
## 8                        DMSO:0.6   1     2  9161 0.082891180  9989  828
## 9                        DMSO:0.8   1     2  9761 0.021649795  9977  216
## 10                       DMSO:1.2   1     2  9739 0.024441551  9983  244
## 11                      untreated   2     3  9558 0.011275473  9667  109
## 12  4-hydroxy norxenovulene B 6:4   2     3  9379 0.012216956  9495  116
## 13  4-hydroxy norxenovulene B 6:6   2     3  9097 0.017920760  9263  166
## 14  4-hydroxy norxenovulene B 6:8   2     3  8968 0.011245865  9070  102
## 15 4-hydroxy norxenovulene B 6:12   2     3  8547 0.017699115  8701  154
## 16                      untreated   2     4  9374 0.001916525  9392   18
## 17                       DMSO:0.4   2     4  9313 0.013766811  9443  130
## 18                       DMSO:0.6   2     4  9153 0.012940796  9273  120
## 19                       DMSO:0.8   2     4  9448 0.010058676  9544   96
## 20                       DMSO:1.2   2     4  9051 0.022147796  9256  205
## 21                      untreated   3     5  9569 0.002709745  9595   26
## 22  4-hydroxy norxenovulene B 6:4   3     5  9107 0.040863612  9495  388
## 23  4-hydroxy norxenovulene B 6:6   3     5  9105 0.025473617  9343  238
## 24  4-hydroxy norxenovulene B 6:8   3     5  9425 0.002962023  9453   28
## 25 4-hydroxy norxenovulene B 6:12   3     5  8907 0.033318863  9214  307
## 26                      untreated   3     6  9504 0.003460208  9537   33
## 27                       DMSO:0.4   3     6  9327 0.024270321  9559  232
## 28                       DMSO:0.6   3     6  9261 0.033903609  9586  325
## 29                       DMSO:0.8   3     6  9306 0.015446466  9452  146
## 30                       DMSO:1.2   3     6  8899 0.048540575  9353  454
```

#### Statistical model

- Generalized linear mixed model
- Avereage proportions of dead cells $\hat{\pi}_j$ are modelled
- Binomial assumption (overdispersed binomial is impossible in `glmer()`)
- Modelling of the response variable was done on logit link $ln\left(\frac{\hat{\pi}_j}{1-\hat{\pi}_j}\right)$
- Repetition and Treatment as fixed effects
- Plates nested in Repetition as random effects

```
fit_t7 <- glmer(cbind(Dead, Alive) ~ Rep + Treatment + (1|Rep:Plate),
                family="binomial",
                data=tox7_num)
```

#### Model based means and their pointwise 95% confidence intervals

- Average proportions of des cells $\hat{\pi}_j$

```
means <- emmeans(fit_t7,
                 specs="Treatment",
                 type="response")

##                        Treatment   prob     SE asymp.LCL asymp.UCL
## 1                      untreated 0.0033 0.0005    0.0025    0.0044
## 4  4-hydroxy norxenovulene B 6:8 0.0119 0.0018    0.0088    0.0160
## 3  4-hydroxy norxenovulene B 6:6 0.0134 0.0020    0.0100    0.0180
## 8                       DMSO:0.8 0.0194 0.0029    0.0144    0.0260
## 6                       DMSO:0.4 0.0249 0.0037    0.0186    0.0334
## 5 4-hydroxy norxenovulene B 6:12 0.0252 0.0037    0.0189    0.0336
## 2  4-hydroxy norxenovulene B 6:4 0.0291 0.0042    0.0219    0.0387
## 9                       DMSO:1.2 0.0384 0.0056    0.0289    0.0510
## 7                       DMSO:0.6 0.0536 0.0076    0.0405    0.0706
```

A graphical overview about this table is given in the manuscript in supplementary figure S 6 c.

#### Multiple comparisons

Pairwise mean comparisons, that are defined below by `cont_list`, were run based on log odds-ratios $ln\left(\frac{\hat{\pi}_j}{1-\hat{\pi}_j}/\frac{\hat{\pi}_{j\prime}}{1-\hat{\pi}_{j\prime}}\right) = ln\left(\frac{\hat{\pi}_j}{1-\hat{\pi}_j}\right) - ln\left(\frac{\hat{\pi}_{j\prime}}{1-\hat{\pi}_{j\prime}}\right)$. The tested null-hypothesis for each comparison was $H_0$: $ln\left(\frac{\pi_j}{1-\pi_j}\right) - ln\left(\frac{\pi_{j\prime}}{1-\pi_{j\prime}}\right) = 0$.

```
# List of contrast coefficients
cont_list

## $`4-hydroxy norxenovulene B 6:4 - untreated`
## [1] -1  1  0  0  0  0  0  0  0
##
## $`4-hydroxy norxenovulene B 6:6 - untreated`
## [1] -1  0  1  0  0  0  0  0  0
##
## $`4-hydroxy norxenovulene B 6:8 - untreated`
## [1] -1  0  0  1  0  0  0  0  0
```

```
##
## $`4-hydroxy norxenovulene B 6:12 - untreated`
## [1] -1  0  0  0  1  0  0  0  0
##
## $`DMSO:0.4 - untreated`
## [1] -1  0  0  0  0  1  0  0  0
##
## $`DMSO:0.6 - untreated`
## [1] -1  0  0  0  0  0  1  0  0
##
## $`DMSO:0.8 - untreated`
## [1] -1  0  0  0  0  0  0  1  0
##
## $`DMSO:1.2 - untreated`
## [1] -1  0  0  0  0  0  0  0  1
##
## $`DMSO:0.4 - 4-hydroxy norxenovulene B 6:4`
## [1]  0 -1  0  0  0  1  0  0  0
##
## $`DMSO:0.6 - 4-hydroxy norxenovulene B 6:6`
## [1]  0  0 -1  0  0  0  1  0  0
##
## $`DMSO:0.8 - 4-hydroxy norxenovulene B 6:8`
## [1]  0  0  0 -1  0  0  0  1  0
##
## $`DMSO:1.2 - 4-hydroxy norxenovulene B 6:12`
## [1]  0  0  0  0 -1  0  0  0  1
```

The table below shows results from the tests that are back transformed to the response scale. Hence the back transformed contrasts become proportions on the response scale $exp\left[ln\left(\frac{\hat{\pi}_j}{1-\hat{\pi}_j}\right) - ln\left(\frac{\hat{\pi}_{j'}}{1-\hat{\pi}_{j'}}\right)\right] = \frac{\hat{\pi}_j}{1-\hat{\pi}_j} / \frac{\hat{\pi}_{j'}}{1-\hat{\pi}_{j'}}$. Similarly, the $H_0$ for the back transformed contrasts changes to $H_0$: $\frac{\pi_{j'}}{1-\pi_{j'}} / \frac{\pi_{j'}}{1-\pi_{j'}} = 1$

```
# Multiple contrast tests
comp <- contrast(means,
                 method=cont_list,
                 adjust="mvt")

##                                      contrast odds.ratio p.value   sig
## 1  4-hydroxy norxenovulene B 6:12 / untreated  7.7557037  <0.001  TRUE
## 2   4-hydroxy norxenovulene B 6:4 / untreated  8.9839760  <0.001  TRUE
## 3   4-hydroxy norxenovulene B 6:6 / untreated  4.0712481  <0.001  TRUE
## 4   4-hydroxy norxenovulene B 6:8 / untreated  3.6005985  <0.001  TRUE
## 5    DMSO:0.4 / 4-hydroxy norxenovulene B 6:4  0.8528535   0.819 FALSE
## 6                        DMSO:0.4 / untreated  7.6620153  <0.001  TRUE
## 7    DMSO:0.6 / 4-hydroxy norxenovulene B 6:6  4.1679982  <0.001  TRUE
## 8                        DMSO:0.6 / untreated 16.9689545  <0.001  TRUE
## 9    DMSO:0.8 / 4-hydroxy norxenovulene B 6:8  1.6423417    0.03  TRUE
## 10                       DMSO:0.8 / untreated  5.9134131  <0.001  TRUE
## 11  DMSO:1.2 / 4-hydroxy norxenovulene B 6:12  1.5441243    0.06 FALSE
## 12                       DMSO:1.2 / untreated 11.9757703  <0.001  TRUE
```

A graphical overview about this table is given in the manuscript in supplementary figure S 6 d.

### Statistical supplementary statS9: Toxicity

Bergmann et al. 2024

```
# Load the packages used for evaluation
library(tidyverse)
library(emmeans)
```

#### Data set

- Compound: Name of the treatments
- Rep: Repetition (timepoint)
- Total: Total number of cells
- Dead: Number of dead cells
- Prop: Proportion of dead cells (for plotting only)

```
toxy

##    Rep                     Compound Total Dead Alive        Prop
## 1    1                 xenovulene B 3  9520  963  8557 0.101155462
## 2    1                 eupenifeldin 2  9358 1431  7927 0.152917290
## 3    1 4-dehydroxy norpycnidione 5  9675  516  9159 0.053333333
## 4    1       4-hydroxyxenovulene B 4  9240 1937  7303 0.209632035
## 5    1                  pycnidione 1  9370 1661  7709 0.177267876
## 6    1 4-hydroxy norxenovulene B 6  9611  695  8916 0.072312975
## 7    1                    untreated  9751   51  9700 0.005230233
## 8    2                 xenovulene B 3  8951 1000  7951 0.111719361
## 9    2                 eupenifeldin 2  9451 1557  7894 0.164744471
## 10   2 4-dehydroxy norpycnidione 5  8932  411  8521 0.046014330
## 11   2       4-hydroxyxenovulene B 4  9261  729  8532 0.078717201
## 12   2                  pycnidione 1  9164 1694  7470 0.184853776
## 13   2 4-hydroxy norxenovulene B 6  9354  326  9028 0.034851400
## 14   2                    untreated  9382   33  9349 0.003517374
## 15   3                 xenovulene B 3  9585  886  8699 0.092436098
## 16   3                 eupenifeldin 2  9767 2313  7454 0.236817856
## 17   3 4-dehydroxy norpycnidione 5  9700  411  9289 0.042371134
## 18   3       4-hydroxyxenovulene B 4  9519  786  8733 0.082571699
## 19   3                  pycnidione 1  9342 2442  6900 0.261400128
## 20   3 4-hydroxy norxenovulene B 6  9779  280  9499 0.028632785
## 21   3                    untreated  9772  136  9636 0.013917315
```

#### Statistical model

- Block design: One observation per compound in each replication
- Generalized linear model
- Binomial proportion for dead cells depend on Rep and Compound
- Logit-Link

```
fit <-  glm(cbind(Dead, Alive) ~ Rep + Compound,
        family=quasibinomial(link="logit"),
        data=toxy)

summary(fit)$dispersion
```

```
## [1] 129.9988
```

- Huge amound of overdispersion:
- The variance is almost 130 times higher than under simple binomial distribution

#### Deviance analysis

- Deviance analysis based on F-Test due to quasi-binomial assumption

```
anova(fit,
      test="F")

## Analysis of Deviance Table
##
## Model: quasibinomial, link: logit
##
## Response: cbind(Dead, Alive)
##
## Terms added sequentially (first to last)
##
##
##          Df Deviance Resid. Df Resid. Dev       F    Pr(>F)
## NULL                        20    13351.9
## Rep       2    178.2        18    13173.7  0.6855    0.5225
## Compound  6  11645.6        12     1528.1 14.9304 6.148e-05 ***
## ---
## Signif. codes:  0 '***' 0.001 '**' 0.01 '*' 0.05 '.' 0.1 ' ' 1
```

- The p-value for the variable compound is far smaller than $\alpha = 0.05$
- At least two compounds differ from each other with regard to their average proportion of dead cells

#### Estimated average proportions and their comparisons

Pairwise comparisons of average proportions $\hat{\pi}_j$ were run on logit scale. Hence, the tested null-hypothesis for each comparison was $H_0: ln\left(\frac{\pi_j}{1-\pi_j}\right) - ln\left(\frac{\pi_{j'}}{1-\pi_{j'}}\right) = 0$. Back transformation to the response scale results in odds ratios $\frac{\hat{\pi}_j}{1-\hat{\pi}_j} / \frac{\hat{\pi}_{j'}}{1-\hat{\pi}_{j'}}$. Hence, back transformation from logit to response scale alters the $H_0$ such that $H_0: \frac{\pi_j}{1-\pi_j} / \frac{\pi_{j'}}{1-\pi_{j'}} = 1$.

```
comp <- emmeans(fit,
                specs="Compound",
                contr="pairwise",
                type="response",
                adjust="mvt")
```

##### Estimated mean proportions of dead cells

```
##                         Compound        prob          SE  df  asymp.LCL   as
ymp.UCL
## 1                      untreated 0.007550266 0.005782888 Inf 0.00167346 0.0
3337510
## 2                   pycnidione 1 0.207168230 0.027687215 Inf 0.15809553 0.2
6664871
```

```
## 3              eupenifeldin 2 0.184836578 0.026188748 Inf 0.13888478 0.2
4172426
## 4              xenovulene B 3 0.100795816 0.020458394 Inf 0.06718123 0.1
4855122
## 5     4-hydroxyxenovulene B 4 0.122629081 0.022328644 Inf 0.08513631 0.1
7350178
## 6 4-dehydroxy norpycnidione 5 0.046832482 0.014268929 Inf 0.02558786 0.0
8419170
## 7 4-hydroxy norxenovulene B 6 0.044942643 0.013896502 Inf 0.02434141 0.0
8152277
```

A graphical overview about this table is given in the manuscript in figure 5 a.

##### Mean comparisons (contrast tests)

```
##                                                          contrast ratio p
.value
## 1                                         untreated / pycnidione 1 0.029
<0.001
## 2                                       untreated / eupenifeldin 2 0.034
<0.001
## 3                                       untreated / xenovulene B 3 0.068
 0.012
## 4                            untreated / (4-hydroxyxenovulene B 4) 0.054
 0.004
## 5                        untreated / (4-dehydroxy norpycnidione 5) 0.155
 0.252
## 6                        untreated / (4-hydroxy norxenovulene B 6) 0.162
 0.281
## 7                                    pycnidione 1 / eupenifeldin 2 1.152
 0.997
## 8                                    pycnidione 1 / xenovulene B 3 2.331
 0.036
## 9                         pycnidione 1 / (4-hydroxyxenovulene B 4) 1.870
 0.202
## 10                    pycnidione 1 / (4-dehydroxy norpycnidione 5) 5.318
<0.001
## 11                    pycnidione 1 / (4-hydroxy norxenovulene B 6) 5.553
<0.001
## 12                                 eupenifeldin 2 / xenovulene B 3 2.023
  0.15
## 13                      eupenifeldin 2 / (4-hydroxyxenovulene B 4) 1.622
 0.525
## 14                  eupenifeldin 2 / (4-dehydroxy norpycnidione 5) 4.615
<0.001
## 15                  eupenifeldin 2 / (4-hydroxy norxenovulene B 6) 4.819
<0.001
## 16                      xenovulene B 3 / (4-hydroxyxenovulene B 4) 0.802
  0.99
## 17                  xenovulene B 3 / (4-dehydroxy norpycnidione 5) 2.281
 0.319
## 18                  xenovulene B 3 / (4-hydroxy norxenovulene B 6) 2.382
 0.269
## 19     (4-hydroxyxenovulene B 4) / (4-dehydroxy norpycnidione 5) 2.845
 0.076
## 20     (4-hydroxyxenovulene B 4) / (4-hydroxy norxenovulene B 6) 2.970
```

```
 0.061
## 21 (4-dehydroxy norpycnidione 5) / (4-hydroxy norxenovulene B 6) 1.044
>0.999
```

A graphical overview about this table is given in the manuscript in figure 5 b.
